# Effector-host networks in the barley powdery mildew pathosystem reveal significant targeting of ubiquitin-associated proteins and transcriptional regulators

**DOI:** 10.64898/2026.10.01.756047

**Authors:** J. Mitch Elmore, Valeria Velásquez-Zapata, Gregory Fuerst, Schuyler Smith, Megan Mueller, Zoi Goebel, Ashton Byal, Matthew J. Moscou, Roger P. Wise

## Abstract

- The ascomycete fungal pathogen *Blumeria hordei* (*Bh*) is predicted to secrete over 530 effector proteins to infect barley and cause powdery mildew disease, yet their individual and collective functions are largely unexplored. We used yeast two-hybrid next-generation interaction screening (Y2H-NGIS) to probe the host targets of 48 *Bh* candidate secreted effectors (CSEPs), including multiple AVR_A_ effectors that are recognized by MLA nucleotide-binding leucine-rich repeat (NLR) receptors.
- Y2H-NGIS and subsequent binary retesting identified 356 high-confidence protein-protein interactions (PPI) among *Bh* CSEPs, MLA NLR fragments, and barley proteins. We uncovered unique and shared host targets, including numerous proteins involved in ubiquitin-associated processes, gene expression, immune and kinase signaling, metabolism, and vesicle trafficking.
- We identified a host protein (*Hv*CAP2) that interacts with MLA and at least five *Bh* effectors, as well as additional effector-host PPI that correlate with loss of recognition of AVR_A13_ and AVR_A6_ by MLA13 and MLA6, respectively.
- Results were integrated with an interolog-based barley interactome (HvInt) to assemble a host-pathogen network of 1248 proteins and 1701 interactions. This PPI resource can serve as a platform for functional investigations into the molecular basis of powdery mildew pathogenesis and host recognition.

## Introduction

Plant pathogens deploy effectors to manipulate the cell signaling and physiology of potential hosts (He *et al*., 2020). Many fungal pathogens are predicted to secrete hundreds of protein effectors during infection (Thordal-Christensen *et al*., 2018; Seong & Krasileva, 2023), yet their individual and collective functions remain largely unknown. Accordingly, key challenges in molecular plant pathology are understanding how each pathogen’s suite of effectors promotes host susceptibility to facilitate parasitic growth and reproduction, and how hosts respond to these pressures (Toruño *et al*., 2016; Arroyo-Velez *et al*., 2020).

The obligate biotrophic fungus *Blumeria hordei* (*Bh)* causes barley powdery mildew, a significant disease that reduces crop yield and grain quality (Smedegaard-Petersen & Stolen, 1981). Genomic analyses of *Bh* have identified over 530 candidate secreted effector proteins (CSEPs), many of which are specifically expressed during infection in intracellular feeding structures known as haustoria (Bindschedler *et al*., 2009; Spanu *et al*., 2010; Pedersen *et al*., 2012; Frantzeskakis *et al*., 2018; Bilstein-Schloemer *et al*., 2025; Sabelleck *et al*., 2025). A conspicuous feature of the *Bh* CSEP repertoire is the expansion of a class of effectors called ribonuclease (RNase)-like proteins associated with haustoria (RALPHs), which comprise up to 70% of the annotated CSEPs in *Blumeria* (Spanu, 2017; Seong & Krasileva, 2023). Despite generally low homology in DNA/protein sequences, RALPHs exhibit structural similarity to fungal RNases (Spanu, 2017; Pennington *et al*., 2019; Cao *et al*., 2023; Lawson *et al*., 2025). However, RALPHs typically lack key catalytic residues and those tested have undetectable RNase activity, suggesting they have undergone neofunctionalization (Spanu, 2017; Pennington *et al*., 2019; Bauer *et al*., 2021; Cao *et al*., 2023). One unresolved issue is to what degree the RNase-like fold has served as a common scaffold for the diversification of host targets and virulence activities, and related, whether or not CSEPs have maintained common host targets during their expansion (Outram *et al*., 2022; Cao *et al*., 2023; Bilstein-Schloemer *et al*., 2025; Knight *et al*., 2026).

The effector-host target landscape is complex, with multiple effectors converging on the same host target and individual effectors targeting multiple host proteins (Mukhtar *et al*., 2011; Weßling *et al*., 2014). For example, the RALPH effector CSEP0064/BEC1054 targets pathogenesis-related protein 10 (PR10), a glutathione S-transferase, a malate dehydrogenase, a thaumatin-like protein PR5, and elongation factor 1 gamma (Pennington *et al*., 2016, 2019). CSEP0264, which belongs to the same subfamily as CSEP0064/BEC1054, also interacts with PR10 *in planta* (Pennington *et al*., 2019). Divergent CSEP0105 and CSEP0162 target small heat shock proteins to compromise their chaperone activity (Ahmed *et al*., 2015). CSEP0162, but not CSEP0105, additionally targets barley Monensin Sensitivity 1 (Liao *et al*., 2023). An ER J-domain protein (HvERdj3B) involved in plant immunity is targeted by sequence-unrelated CSEP0008/AVR_A1_ and CSEP0491/BEC1016 (Li *et al*., 2024b). These observations indicate that CSEPs can have a diversity of targets *in planta* but relatively divergent CSEPs might also converge on a limited number of host processes. However, because interactors for only 14 effectors (less than 3% of the *Bh* effector complement) have been reported, it is challenging to draw overarching conclusions about the molecular basis of *Bh* virulence (Bilstein-Schloemer *et al*., 2025).

Nucleotide-binding leucine-rich-repeat (NLR) receptors encoded at *Mildew locus a* (*Mla*) in barley can activate full immunity against particular *Bh* isolates by recognizing specific avirulence (AVR) effectors (Wei *et al*., 1999; Halterman *et al*., 2001; Zhou *et al*., 2001; Seeholzer *et al*., 2010). Cognate pairs of MLA-AVR have been identified, such as MLA1-CSEP0008/AVR_A1_, MLA6-BLGH_00708/AVR_A6_, MLA7-CSEP0059/AVR_A7_, MLA9-CSEP0174/AVR_A9_, MLA10-CSEP0141/AVR_A10_, and MLA13-CSEP0372/AVR_A13_ (Lu *et al*., 2016; Saur *et al*., 2019; Bauer *et al*., 2021). Current evidence indicates that the MLA receptors physically interact with the AVR to activate immunity, although additional host proteins may regulate complex formation (Saur *et al*., 2019; Lawson *et al*., 2025). *In silico* and experimental structural characterization indicates that *Bh* AVR_A_ effectors cloned to date belong to the RALPH class of CSEPs (Bauer *et al*., 2021; Cao *et al*., 2023; Lawson *et al*., 2025). While RALPH experimental structures share a common core of two β-sheets with a central α-helix, there is considerable variability in the number and length of the β-strands comprising each β-sheet, the length of the α-helix, and the loop regions connecting these elements, indicating substantial differentiation across RALPH subfamilies. It is unknown to what extent this variation influences the spectrum of cellular targets (Pennington *et al*., 2019; Cao *et al*., 2023; Lawson *et al*., 2025).

While a limited number of individual effectors and their host targets have been investigated (Bilstein-Schloemer *et al*., 2025), the overwhelming majority of the *Bh* effector arsenal remains functionally uncharacterized, leaving the broader framework of the cereal-*Blumeria* interactome uncharted. In this study, we present a large-scale protein-protein interaction (PPI) map between 48 *Bh* effectors and barley using yeast two-hybrid next generation interaction screening (Y2H-NGIS) (Lewis *et al*., 2012; Suter *et al*., 2015; Pashkova *et al*., 2016; Velásquez-Zapata *et al*., 2021, 2023; Elmore *et al*., 2023). We present evidence that *Bh* effectors target critical biological processes, such as post-translational modifications, transcriptional regulation, immune signaling, metabolism, and vesicle trafficking (Li *et al*., 2024b; Sabelleck *et al*., 2025; Thordal-Christensen *et al*., 2026; Deb *et al*., 2026). We identified examples of related CSEPs targeting the same or similar host proteins, as well as divergent CSEPs interacting with the same host targets. Notably, two host proteins interact with AVRs and the CC/NB domains of MLA; a calcium-dependent protein kinase 1 adaptor protein 2 (*Hv*CAP2) and a ubiquitin CUE domain protein. Finally, we found that effector residues important for AVR_A6_ and AVR_A13_ recognition are also critical for interactions with select host targets.

## Materials and Methods

### Plant and fungal materials

Barley Cereal Introduction (CI) 16151 (*Mla6*) and derived mutants are described in (Chapman *et al*., 2021). *Bh* isolate 5874 (*AVR_a1_, AVR_a3_, AVR_a6_, AVR_a12_*) was maintained on susceptible *H. vulgare* cv. Morex in a growth chamber at 18 °C (16/8 hour light/darkness).

### Differential transcript accumulation

Transcriptome data were obtained from an infection time course of CI 16151 and the mutant *mla6*-m18982 at 0, 16, 20, 24, 32 and 48 hours after inoculation (HAI) with *Bh* isolate 5874 (NCBI-GEO GSE101304) and analyzed as described in (Chapman *et al*., 2021; Velásquez-Zapata *et al*., 2022, 2024). Genes differentially expressed at an adjusted (adj.) *p* < 0.001 for barley and adj. *p* < 0.003 for *Bh* were considered significant.

### Yeast two-hybrid, next-generation interaction screening (Y2H-NGIS)

The Y2H system consisted of pDEST-AD (Activation Domain, pPC86-based, *TRP1*) and pDEST-DB (DNA-Binding domain, pPC97-based, *LEU2*) and yeast strains Y8800 and Y8930 (Dreze *et al*., 2010; Mukhtar *et al*., 2011; Weßling *et al*., 2014). A size-selected cDNA library generated from RNA from the infection time course was cloned into pDONR222 in three frames using the CloneMiner II kit (Invitrogen, Carlsbad, CA, USA) and subsequently transferred to pDEST-AD using Gateway cloning (Invitrogen, Carlsbad, CA, USA). The prey library was transformed into strain Y8800, yielding 1.1 x 10^7^ primary clones (Gietz & Schiestl, 2007). Y2H bait sequences were cloned into pENTR221 or pCR8, transferred to pDEST-DB, and transformed into Y8930. Three biological replicates of each Y8930 bait strain were screened by mating to the Y8800 prey library as described in (Elmore *et al*., 2023). Diploid yeast were grown in synthetic complete liquid media lacking leucine and tryptophan (SC-LW, library control) and sub-cultured twice under selective SC-LWH media lacking histidine to enrich for interactions that activate the *HIS3* reporter gene (Pashkova *et al*., 2016; Elmore *et al*., 2023). After subculturing, yeast were harvested *en masse* and plasmids were purified from each cell population (Elmore *et al*., 2023). Prey gene fragments were then amplified by low-cycle PCR and each amplicon library was sequenced to a depth of 20 million reads by GENEWIZ (Morrisville, NC, USA). NGPINT v3 software was used to process sequence data (Banerjee *et al*., 2021; Smith *et al*., 2025). Candidate interactors were ranked via enrichment, in-frame, and specificity scores using Y2H-SCORES (Velásquez-Zapata *et al*., 2021, 2023).

### Binary Y2H confirmation

In-frame sequences of interacting prey fragments were determined using read alignments obtained from NGPINT. Custom sequences were synthesized by Gene Universal (Newark, DE, USA). Binary Y2H tests were performed according to (Dreze *et al*., 2010; Velásquez-Zapata *et al*., 2022; Li *et al*., 2024b; Deb *et al*., 2026) by mating bait and prey strains, and plating diploids on media with selection for diploids (SC-LW), or interactions at four selective stringencies (SC-LWH; SC-LWH + 0.1 mM 3-AT; SC-LWH + 0.5 mM 3-AT, SC-LWH + 1.0 mM 3-AT). The following interaction score was assigned based on yeast growth: 0 = no colonies present on any selection plates; 1 = colonies present on SC-LWH; 2 = colonies present on SC-LWH + 0.1 mM 3-Amino-1,2,4-triazole (3-AT); 3 = colonies present on SC-LWH + 0.5 mM 3-AT; 4 = colonies present on SC-LWH + 1.0 mM 3-AT; and 5 = >15 colonies present on SC-LWH + 1.0 mM 3-AT.

### Determination of a high-confidence set of PPI

From the binary Y2H assays, an interaction score ≥ 2 was considered a positive interaction. ROC curves were generated in R using this binary outcome as a function of Enrichment score. To determine an ideal threshold for the complete dataset, the benchmark data was used to plot performance metrics accuracy, precision, sensitivity (TPR, True Positive Rate), specificity (1 - FPR, False Positive Rate), and Youden Index (TPR - FPR) at possible Enrichment scores every 0.001 from 0.000 - 1.000. The enrichment score at the highest Youden Index was considered the optimal threshold. From this analysis, interactions with an Enrichment score ≥ 0.80 were assigned as high-confidence interactions and named the “Enrich80” set.

### Gene and protein functional classifications

Protein domains within the barley cultivar Morex v3 proteome were annotated using InterProScan v5.73-104.0 (Jones *et al*., 2014; Mascher *et al*., 2021; Blum *et al*., 2025). Short functional descriptions were generated using AHRD v3.1.1 (Mascher *et al*., 2021). Protein kinases and transcription factors were annotated using iTAK v2.0.2 (Zheng *et al*., 2016). Ubiquitin-associated proteins were classified using previously published domain annotations, along with the addition of some plant-specific domains. (Zhou *et al*., 2018; Castellanos *et al*., 2020). Barley gene orthology and homology to wheat, rice, and *Arabidopsis* was determined using the EnsemblCompara database release 113 (Herrero *et al*., 2016) and BLAST to the respective genomes (Camacho *et al*., 2009).

### Protein structure clustering and multimer predictions

For CSEP structural predictions, signal peptide-trimmed CSEP sequences were used as input to AlphaFold2 v2.3.2 with parameters: three recycles and max template date of May 2^nd^ 2026 (Jumper *et al*., 2021). Select structures were predicted with the AlphaFold3 webserver v3.0.1 as indicated in the text (Abramson *et al*., 2024). Top-ranked models was used for subsequent analyses. To create structural dendrograms of CSEP relatedness, AlphaFold2-predicted structures were subjected to all-by-all alignment using Reseek v2.8 (Edgar, 2024). Pairwise alignment quality (AQ) scores were used to create a distance matrix (1 - AQ) and hierarchical clustering was performed using the R v4.6.0 stats::hclust function with average linkage. Dendrograms were visualized with associated metadata using the R packages ggtree v4.2.0 and ggtreeExtra v1.22.1 (Yu *et al*., 2017; Xu *et al*., 2021). For AlphaFold2-Multimer analyses, prey sequences corresponding to the NGPINT-predicted interaction fragment with the most in-frame reads were modeled with the corresponding CSEP bait using AlphaPulldown v2.9.0 (Evans *et al*., 2022; Yu *et al*., 2023). The best-scoring multimer model based on highest ipTM-pTM score was used as input to AlphaJudge v1.4.2 (https://github.com/KosinskiLab/AlphaJudge) to calculate per-model and per-interface scores. Differences in the score distributions were evaluated using Wilcoxon rank-sum tests with Benjamini-Hochberg multiple test correction and visualized with violin plots using ggplot2 v4.03.

### InterPro domain enrichment

InterPro domain enrichment was performed in R using the enricher function from clusterProfiler v.4.20.0 using a one-sided Fisher’s exact test with Benjamini-Hochberg correction (Yu *et al*., 2012). InterPro term reduction was performed within all terms for all baits from the enricher output using a Szymkiewicz– Simpson coefficient approach based on the overlap in genes for each term in the proteome-wide InterProScan annotation using a similarity threshold of 0.85 (Simpson, 1949). Representative terms for each overlap cluster were selected using the maximum number of genes associated with each term.

### Network graphs

Network visualizations were produced in Cytoscape v3.10.4 through the RCy3 interface in R (Shannon *et al*., 2003; Gustavsen *et al*., 2019). To contextualize the interactions within the broader barley interactome, we integrated them with the predicted interolog network HvInt by retaining the first-neighbor nodes of all preys in the Enrich80 set (Velásquez-Zapata *et al*., 2022).

## Results

### Selection of CSEPs for Y2H-NGIS

For identification of *Bh*-host PPI, CSEPs were selected based on: **1**) documented virulence activities based on transient overexpression or host-induced gene silencing (HIGS), or recognition by *Mla* in the case of CSEP0008/AVR_A1_, BLGH_00708/AVR_A6_, CSEP0059/AVR_A7_, CSEP0174/AVR_A9_, and CSEP0372/AVR_A13_-1 (Zhang *et al*., 2012; Pliego *et al*., 2013; Schmidt *et al*., 2014; Aguilar *et al*., 2016; Lu *et al*., 2016; Saur *et al*., 2019; Bauer *et al*., 2021), **2**) degree of gene conservation across eight *Blumeria graminis* formae speciales (Frantzeskakis *et al*., 2018), **3**) differential gene expression in *Bh* isolate 5874 (*AVR_a6_*) during infection of resistant CI 16151 harboring the *Mla6* immune receptor and susceptible mutant *mla6*-m18982 (Velásquez-Zapata *et al*., 2024), and **4**) peak expression at different stages of infection (**Fig. 1, Supporting Information Fig. S1, Fig. S2**). CSEP structures were predicted with AlphaFold2 and clustered based on their structural similarity, which agreed with previous classifications (**Fig. 1, Fig. S1**) (Seong & Krasileva, 2023). The majority of screened CSEPs (∼80%) belong to the RNase-like superfamily (Seong & Krasileva, 2023), and we sampled CSEPs from different subclades within this group (**Fig. S1**). Some effectors are expressed highest at early stages of infection (16 - 20 HAI), while most have peak expression during haustorial development (32 - 48 HAI) in the *mla6*-m18982 genotype. Eight of the 48 displayed higher expression in incompatible, as opposed to compatible interactions (**Fig. S2**). In addition to CSEPs, we screened the conserved, non-CSEP effector protease BEC1019, fragments of the MLA6 NLR comprising the CC (MLA6_1-161_), CC-NB (MLA6_1-225_), and LRR (MLA6_550-956_) domains, and the cytosolic juxtamembrane kinase (JMK) domain of the receptor-like kinase *Hv*SRF3_L1 (HORVU.MOREX.r3.7HG0739520), which we found to interact with CSEP0372/AVR_A13_-1 in initial tests. Luciferase was used as a negative control bait.

**Fig. 1.**
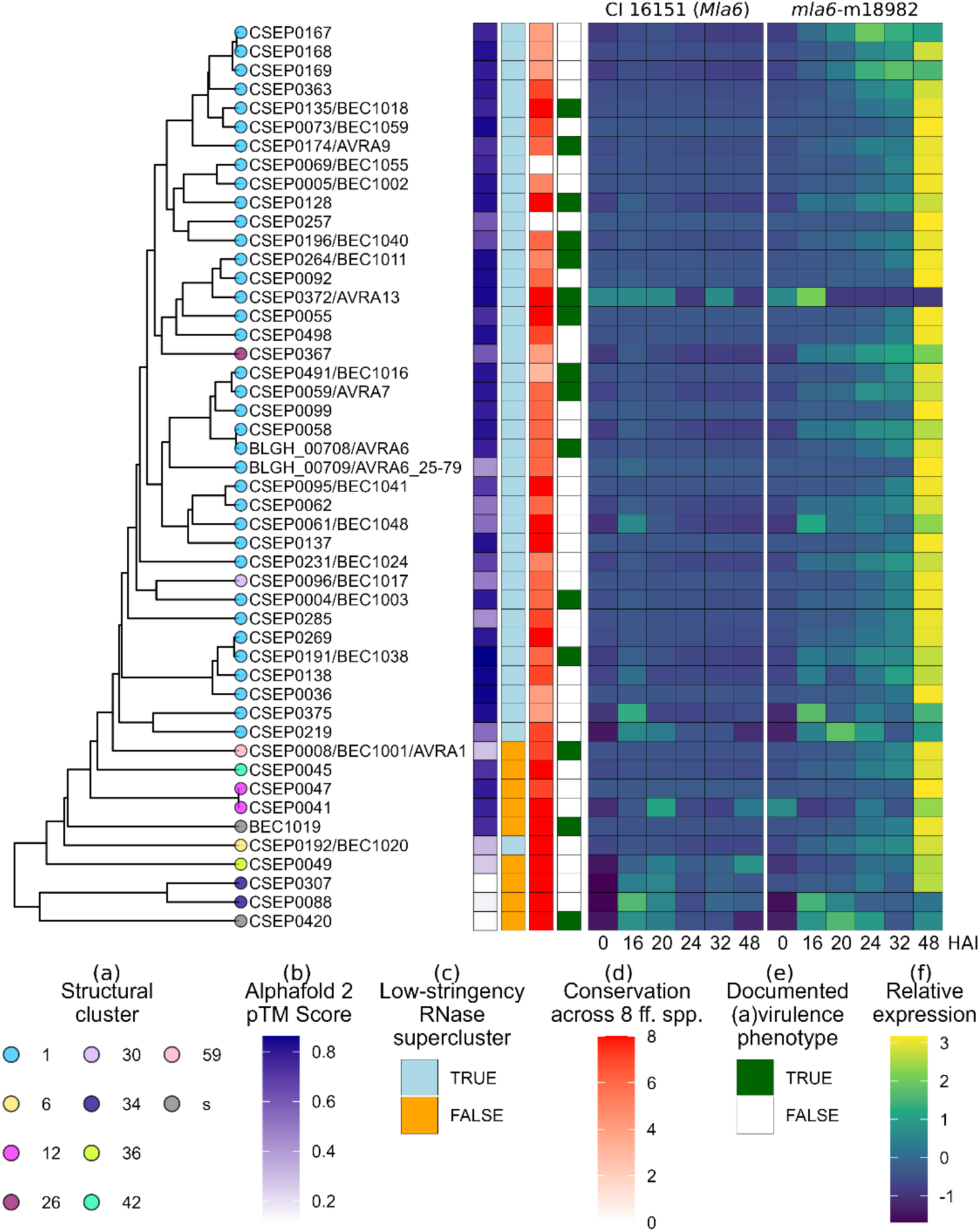
Overview of *Bh* effectors screened by Y2H-NGIS. (**a**) Dendrogram based on pairwise alignments of AlphaFold2-predicted structures. Tip labels correspond to the structural family classification of (Seong & Krasileva, 2023). (**b**) AlphaFold2 predicted Template Modeling (pTM) score. (**c**) RNase-like supercluster designation from (Seong & Krasileva, 2023). (**d**) Conservation of gene orthologs across 8 *Blumeria* formae speciales (ff. spp.) published in (Frantzeskakis *et al*., 2018). (**e**) Documented avirulence or virulence phenotypes (Bilstein-Schloemer *et al*., 2025). (f) Relative expression (z-score) of effectors in *Blumeria hordei* isolate 5874 (*AVR_a6_)* during infection of barley cultivar CI 16151 (*Mla6*) (incompatible interaction) and CI 16151-derived mutant *mla6*-m18982 (compatible interaction). HAI, hours after inoculation. Transcriptome data were published in (Velásquez-Zapata *et al*., 2024).

### Y2H-NGIS and defining a high-confidence set of effector-host PPI

To discover *Bh*-host interactions, we adapted the DEEPN protocol to screen the selected baits against a cDNA library derived from a *Bh* infection time course (Pashkova *et al*., 2016; Elmore *et al*., 2023; Velásquez-Zapata *et al*., 2024). The sequencing data were analyzed with NGPINT to align to the *Bh* and barley genomes, obtain gene counts and test for enrichment, identify fusion reads (chimeric reads that contain Y2H plasmid and prey cDNA sequences), and determine whether cDNA fragments were in-frame with the Y2H GAL4 AD (Banerjee *et al*., 2021; Velásquez-Zapata *et al*., 2023; Smith *et al*., 2025).

From the NGPINT output, each bait had an average of 20.1 significantly enriched preys (adj. p < 0.01; median: 14, range: 0 - 122) relative to the non-selected controls across three independent replicates. Y2H-SCORES ranks candidate interactions by calculating enrichment, specificity, and in-frame scores (each on a 0 - 1 scale) (Velásquez-Zapata *et al*., 2021, 2023). Using the Y2H-SCORES rankings, we performed binary Y2H (i.e., one-to-one) retesting to confirm positive interactions (Dreze *et al*., 2010), while including lower scoring and off-target interactions (**Table S3**). All prey fragments were tested against the bait empty vector and Luciferase as negative controls, which identified two auto-active and/or non-specific preys in the cDNA library: the transcription factor VOZ1 (HORVU.MOREX.r3.3HG0289300) and a receptor-like kinase (RLK) HORVU.MOREX.r3.2HG0132360. These preys were removed from further consideration.

Using the binary retesting as benchmark data (n = 122), we plotted a Receiver Operating Characteristic (ROC) curve to evaluate the diagnostic ability of the Y2H-SCORES ranking and determine optimal thresholds for the full dataset. This analysis demonstrated that the Enrichment score had good predictive ability, with an AUC (area under the curve) value of 0.8179 (**Fig. 2a**). We next calculated Accuracy, Precision, Sensitivity (TPR), Specificity (1 - FPR), and Youden Index (TPR - FPR) at possible Enrichment score thresholds (**Fig. 2b**). The Youden Index identified an optimal Enrichment score of 0.802 for predicting positive interactions in the full dataset. This threshold achieved a sensitivity of 0.75, a specificity of 0.81, and precision of 0.882 (i.e., 88.2% of binary retests above this threshold yielded a positive interaction). For simplicity, we selected an Enrichment score cutoff of 0.80 (sensitivity: 0.75, specificity: 0.79, precision: 0.87) which predicts a total of 356 high-confidence interactions in the dataset, and we refer to these interactions as the “Enrich80” set. At this threshold, each bait had an average of 6.8 interactions (median: 3.5, range: 0 - 57) (**Fig. 2c**). CSEP0047 and CSEP0062 had the most interactions with 47 and 57 preys, respectively. No interactions with an enrichment score greater than 0.80 were identified for CSEP0041, CSEP0059/AVR_A7_, CSEP0191, CSEP0285, or BEC1019 although some positive tests were recovered below this threshold. On average, each prey was targeted by 1.31 baits (median: 1, range: 1 - 15), with ∼82% of the preys interacting with a single bait (**Fig 2d)**. Notably, the ubiquitin domain protein HORVU.MOREX.r3.3HG0231510 was targeted by 15 CSEPs but did not yield a positive binary Y2H result with BD-EV, Luciferase, CSEP0045, or CSEP0420, so it was retained (**Table S3**).

**Fig 2.**
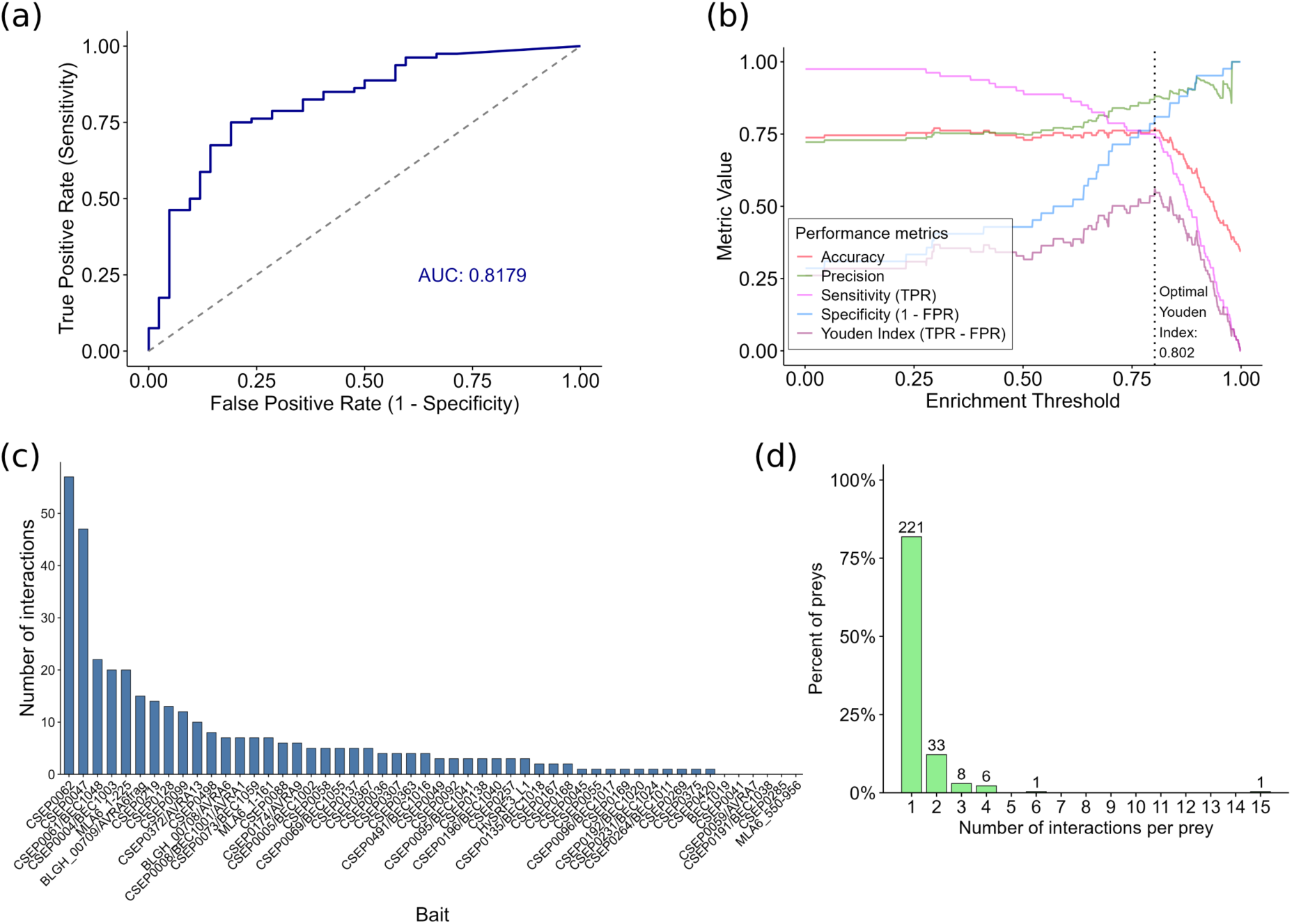
A high-confidence set of *Bh*-host interactions. (**a**) Receiver Operating Characteristic (ROC) curve of a binary classification model using enrichment score to predict an interaction. AUC, Area under the ROC curve. (**b**) Line plot of Accuracy, Precision, Sensitivity (TPR, True Positive Rate), Specificity (1 - FPR, False Positive Rate), and Youden Index (TPR - FPR) performance metrics as a function of enrichment score threshold. The vertical dotted line indicates the enrichment score at the optimal Youden Index. (**c**) Bar plot of the number of high-confidence interactions (enrichment score >= 0.80) for each bait. (**d**) Frequency bar plot depicting the percent of preys with the specified number of high-confidence interactions. The number of preys is above each bar.

A graphical representation of the resulting “Enrich80” interactome is presented in **Fig. 3, Supporting File S1**. We then integrated Enrich80 with the interolog-based interactome from *H. vulgare,* HvInt (Velásquez-Zapata *et al*., 2022), comprising 66,133 edges and 7,181 nodes. This two-step strategy enabled the construction of subnetworks that highlight both direct Y2H-supported interactions and their immediate predicted interaction environment within HvInt, facilitating the identification of effector-centered modules and potential signaling or defense-related pathways (**Fig. S3, File S2)**.

**Fig. 3.**
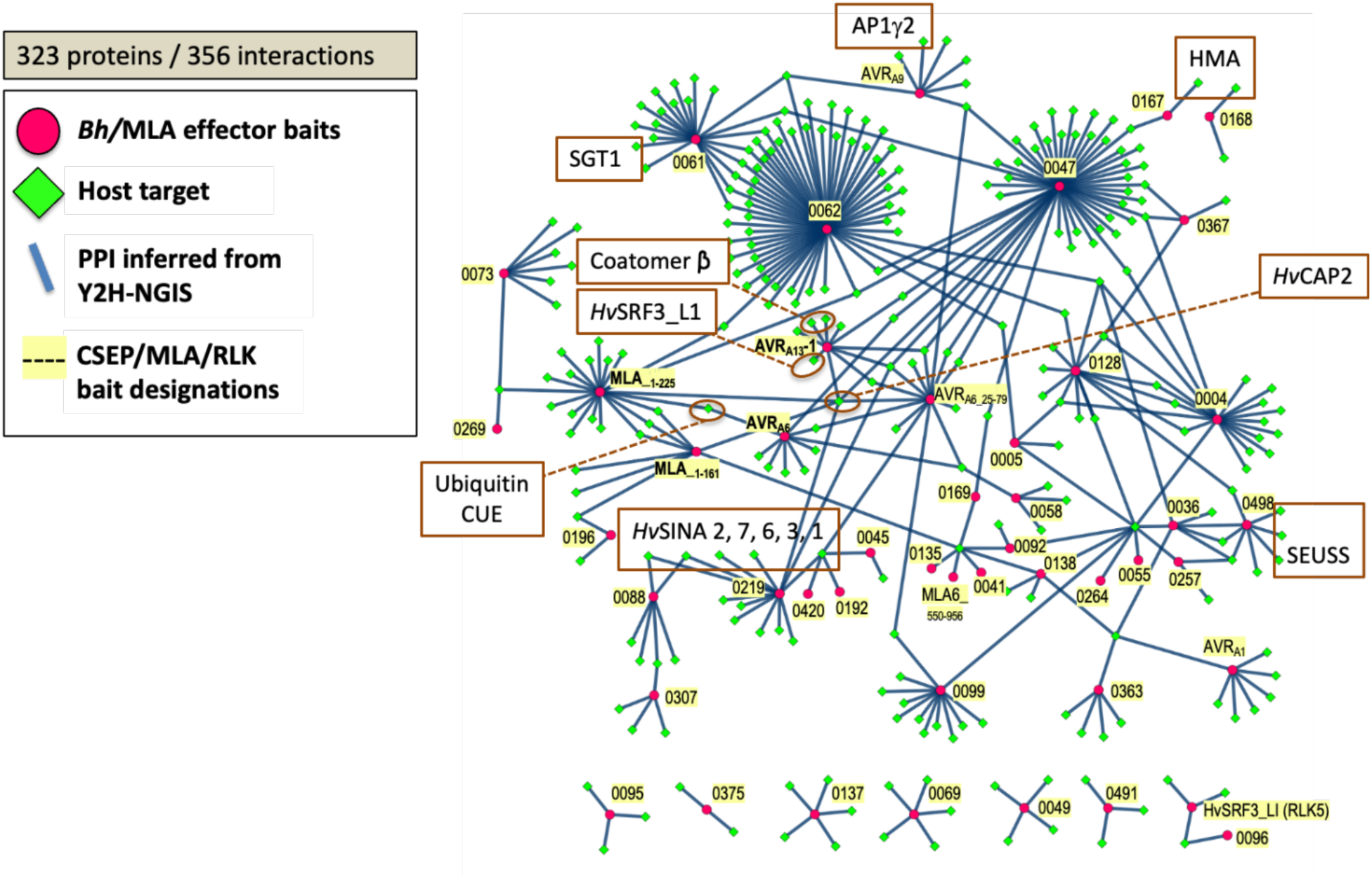
Pathogen-host Y2H interactions in the “Enrich80” network. Graphical representation of Y2H-NGIS interactions at an enrichment cutoff of 0.80. Red-filled circles designate *Hv*MLA or *Bh* effector baits (yellow highlight), green-filled diamonds signify host targets (the identity of selected proteins is indicated in brown boxes), blue solid lines represent PPI inferred from 52 Y2H-NGIS screens in the current report.

### Multimer predictions support the identified *Bh*-host interactions

To gauge whether the identified pairs are predicted to have compatible binding interfaces, we used AlphaFold2-Multimer to model PPI *in silico* (Evans *et al*., 2022; Yu *et al*., 2023). We used the detected in-frame prey fragments to model all interactions with Enrichment score greater than 0, those tested in binary assays, and a randomly selected subset of 400 bait-prey pairs with an Enrichment score of 0, totaling 2141 interactions (**Table S4**). From this set, we detected a small, but significant, positive Spearman correlation between Enrichment score and the AlphaFold2-derived interface predicted Template Modeling (ipTM) score (*r*_s_ = 0.0821, *p* = 1.44 x 10^-4^) (**Fig. S4a**) as well as a significant difference in ipTM score distributions between the Enrich80 set and lower-confidence interactions (Wilcoxon rank-sum test, *p* = 0.0016) (**Fig. S4b)**. As many of the multimer models contained at least one chain that was not confidently predicted, we enforced a mean predicted Local Distance Difference Test (pLDDT) score threshold above 0.65 for both bait and prey chains, resulting in 681 higher-confidence models. From this set, we detected greater differences in the distributions of ipTM and nearly all other interface confidence scores between the Enrich80 set and those below it (**Fig. S4c**). Together, these analyses provide information on potential binding interfaces and orthogonal support for our set of high-confidence interactions, as well as many lower-scoring PPI (**Fig. S4a, Table S4**).

### InterPro domain enrichment identifies commonly targeted host proteins

To functionally classify the candidate interactions, we performed InterPro domain enrichment for preys associated with each CSEP bait, as well as the non-redundant list of all CSEP-targeted preys in the high-confidence Enrich80 set (n = 231) (**Table S5**). Proteins with zinc finger domains were most frequent, with the RING/FYVE/PHD-type homologous superfamily (IPR013083) present in ∼10% (n = 24) of all identified preys (**Fig. 4**). The ubiquitin-like superfamily (IPR029071) was found in ten preys with diverse domain architectures. DNA-binding and transcriptional regulator domains also dominated the list, with SEUSS-like, Myc-type basic Helix Loop Helix (bHLH), MYB/SANT-like, Auxin response factor, Dof zinc finger, Aux/IAA, MYB, WRKY, and TEOSINTE BRANCHED1/CYCLOIDEA/PROLIFERATING CELL FACTOR (TCP) proteins targeted across at least eight CSEPs. Eight preys contain a protein kinase domain (IPR000719), although it was not significantly enriched due to the high background of protein kinases in the barley proteome. **Fig. S5** depicts the protein domains associated with all prey interactors for each bait (**Table S4**).

**Fig 4.**
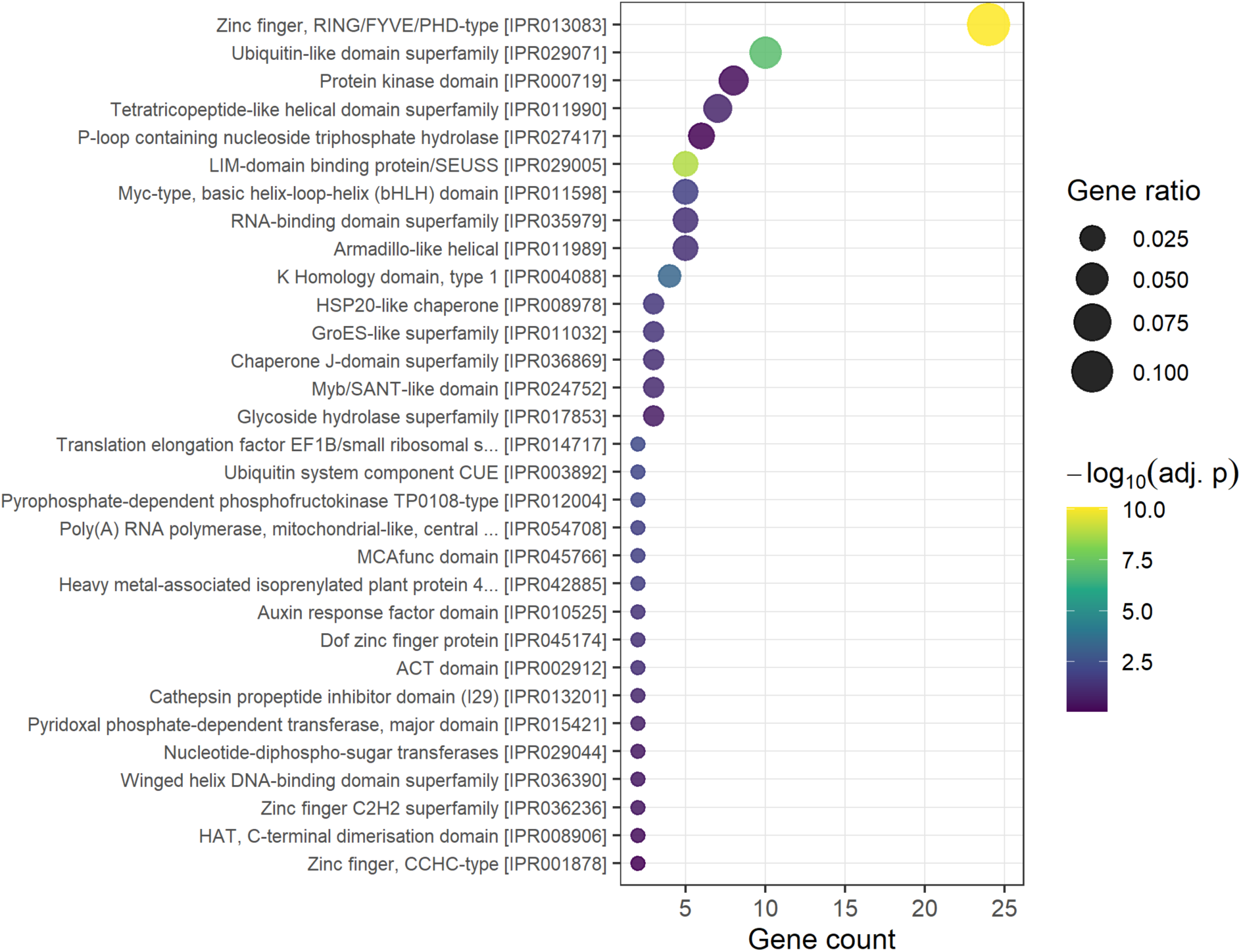
Common protein classes targeted by CSEPs. Bubble plot depicting Interpro terms associated with the non-redundant set of CSEP-targeted proteins (n = 231). Data points are colored according to the adjusted p-value (one-sided Fisher’s exact test, Benjamini-Hochberg correction) and sized relative to the number of genes for each term divided by the total number of genes. Terms represented by at least two proteins are shown.

### Ubiquitin-associated proteins are a major target of *Bh* CSEPs

Among proteins detected with Zinc finger RING/FYVE/PHD-type domains, there are distinct classes of E3 ubiquitin-protein ligases, including members of the SEVEN IN ABSENTIA (SINA), MEDIATOR25 -BINDING RING-H2 PROTEIN (MBR1/2-like), RED AND FAR-RED INSENSITIVE 2 (RFI2), RETINOBLASTOMA-BINDING PROTEIN 6 (RBBP6), MAHOGUNIN RING FINGER 1/ RING FINGER PROTEIN 157 (MGRN1/RNF157-like), RING FINGER PROTEIN 8 (RNF8), and RING FINGER PROTEIN 127 (RNF127) families.

Notably, CSEP0047 interacted with diverse E3 ligases, including RNF127-type RRS1-ASSOCIATED RING-TYPE E3 (RARE) homolog HORVU.MOREX.r3.4HG0374770 (Chen *et al*., 2025), RFI2-type AVRPIZ-T INTERACTING PROTEIN 6 (APIP6) ortholog HORVU.MOREX.r3.1HG0023230 (also targeted by CSEP0367) (Park *et al*., 2012; Shi *et al*., 2024), RNF8-type RPM1-INTERACTING PROTEIN 2 (RIN2) homolog HORVU.MOREX.r3.2HG0146750 (also targeted by CSEP0372/AVR_A13_) (Kawasaki *et al*., 2005), BOTRYTIS SUSCEPTIBLE1-INTERACTOR (BOI) homologs HORVU.MOREX.r3.4HG0389860 and HORVU.MOREX.r3.7HG0730970 (Luo *et al*., 2010), and MBR1/2-like MUTANT SNC1 ENHANCING 16 (MUSE16) homolog HORVU.MOREX.r3.3HG0235900 (also targeted by CSEP0061/BEC1048) (Huang *et al*., 2023), all of which have reported roles in regulating disease resistance in *Arabidopsis* and/or rice. In addition to the MUSE16 homolog, other MBR1/2-like E3 ligases were targeted by more effectors, suggesting that this E3 family is a key focal point of *Bh* pathogenesis. In total, four MBR1/2-like E3 ligases were targeted across six CSEPs. HORVU.MOREX.r3.1HG0088310 interacted with four effectors: CSEP0004/BEC1003/BEC5, CSEP0128, CSEP0196/BEC1037/BEC1040, and CSEP0367. CSEP0004/BEC1003/BEC5 also interacted with MBR1/2-like HORVU.MOREX.r3.2HG0204830 and HORVU.MOREX.r3.3HG0278640. All of these barley proteins belong to a family, including MUSE16, MBR1/2, TIE1-ASSOCIATED RING-TYPE E3 LIGASE1 (TEAR1), and SNIPER8, that has roles in the degradation of transcriptional activators and repressors, JA signaling, leaf development, and regulation of immune responses in *Arabidopsis* (Zhang *et al*., 2017; Huang *et al*., 2023; Wu *et al*., 2025; Liu *et al*., 2026), with homologs in rice that contribute to *Magnaporthe oryzae* resistance (Zhong *et al*., 2018; Liu *et al*., 2021).

A diverse set of additional ubiquitin ligases interacted with various CSEPs. Another RFI2-type E3 related to rice APIP6, HORVU.MOREX.r3.7HG0685620, interacted with CSEP0004/BEC1003/BEC5. CSEP0219 interacted with five distinct SINA E3 ligases (HORVU.MOREX.r3.7HG0711610, HORVU.MOREX.r3.1HG0034990, HORVU.MOREX.r3.3HG0249640, HORVU.MOREX.r3.2HG0116790, HORVU.MOREX.r3.6HG0553110), which are important regulators of multiple hormone signaling pathways, abiotic stress responses, and plant immunity (Miao *et al*., 2016; Ren *et al*., 2021, 2024; Deb *et al*., 2026). The related, non-RALPH effectors CSEP0088 and CSEP0307 interacted with E3 CONSTITUTIVE PHOTOMORPHOGENIC 1 (COP1) homolog HORVU.MOREX.r3.6HG0618700. COP1 proteins function in light and hormone signaling, development, and stress responses, and have also been implicated in maintaining NLR protein abundance (Lim *et al*., 2018; Nan & Hou, 2025). CSEP0073/BEC1059 interacted with HORVU.MOREX.r3.4HG0391130, a barley homolog of MGRN1/RNF157-like LOSS OF GLUTAMINE DUMPER 1 (LOG2), an E3 reported to regulate amino acid export at the plasma membrane (Pratelli *et al*., 2012). CSEP0062 and CSEP0128 interacted with HORVU.MOREX.r3.6HG0603690, a homolog of PARAQUAT TOLERANCE 3, an RBBP6-family E3 ligase that functions as a negative regulator of the oxidative stress response in *Arabidopsis* (Luo *et al*., 2016). CSEP0092 interacted with the barley BRIZ1 homolog HORVU.MOREX.r3.2HG0204960, an E3 associated with abscisic acid (ABA) signaling (Linden *et al*., 2021). CSEP0073/BEC1059 and CSEP269 interacted with HORVU.MOREX.r3.4HG0389410, a barley ortholog of the *Arabidopsis* SH3 domain proteins (SH3Ps). SH3Ps have been implicated in endocytosis and the polyubiquitination of immune regulators (Kong *et al*., 2021; Adamowski *et al*., 2024). Collectively, these candidate interactions suggest that one prevailing function of *Bh* CSEPs is to target host ubiquitination processes to modulate hormone, stress response, and immune signaling pathways (**Table S2**).

In addition to ubiquitin ligases, we identified multiple ubiquitin and ubiquitin-like proteins, as well as larger proteins with ubiquitin-like domains. Of note, related effectors interacted with ubiquitin (CSEP0062) and ubiquitin-like NEDD8 proteins (CSEP0061/BEC1048). Interestingly, homologs of B CELL LYMPHOMA 2-ASSOCIATED ATHANOGENE 6 (BAG6) with an N-terminal ubiquitin-like domain were associated with multiple CSEPs. As mentioned above, BAG6-like HORVU.MOREX.r3.3HG0231510 had high-confidence interactions with 15 CSEPs (**Fig. 2d**), and interacted with most, but not all, of the CSEPs tested in binary assays. Another BAG6-like protein, HORVU.MOREX.r3.6HG0541060, appeared to be more specific, with high-confidence interactions with only CSEP0004/BEC1003/BEC5 and CSEP0128. In *Arabidopsis* and potato, orthologs of BAG6-like proteins regulate proteasome activity, activation of autophagy, and disease resistance against fungal and oomycete pathogens (Li *et al*., 2016; Wang *et al*., 2023).

Some CSEPs target proteins associated with autophagic processes. CSEP0174/AVR_A9_ interacted with DOMINANT SUPPRESSOR OF KAR 2 (DSK2) ortholog HORVU.MOREX.r3.4HG0409910, a selective autophagy receptor that interfaces with brassinosteroid and gibberellin signaling, and has been implicated in the degradation of the chitin receptor LYK5 in wheat (Wu *et al*., 2026). CSEP0047 interacted HORVU.MOREX.r3.6HG0590780, an ortholog of NEIGHBOR OF BRCA1 (NBR1), which is a selective autophagy receptor that recognizes ubiquitinated cargo and directs it to ATG8-decorated autophagosomes for degradation (Li *et al*., 2026). CSEP0047 also interacted with PHAGOPHORE ASSEMBLY FACTOR 1 homolog HORVU.MOREX.r3.7HG0721230, with putative functions in autophagosome formation (Kojima *et al*., 2021). Additionally, CSEP0047 interacted with the E3 ufmylation ligase UFM1-protein ligase 1 homolog (UFL1) HORVU.MOREX.r3.1HG0007850 that is part of a conserved autophagy receptor complex at the endoplasmic reticulum (ER) that is activated by ribosome stalling and critical for ER stress responses (Stephani *et al*., 2020).

### CSEP interactions with Transcriptional regulators

A diversity of proteins involved in gene expression, including canonical transcription factors (TFs) and other transcriptional regulators, was targeted across eight CSEPs, totaling 32 interactions (**Table S2**). CSEP0062 and CSEP0047 accounted for over 60% of these PPI. We uncovered CSEP0047 and CSEP0498 interacting with different homologs of SEUSS, a transcriptional co-repressor involved in plant development and abiotic stress, with possible roles in immunity (Sridhar *et al*., 2004; Bui, 2009; Shrestha *et al*., 2014). Among the basic helix-loop-helix TFs, were an IAA-LEUCINE RESISTANT3 homolog (HORVU.MOREX.r3.2HG0138530) targeted by CSEP0128 and CSEP0498 that has been shown in *Arabidopsis* to regulate iron and immune responses (Herlihy *et al*., 2020). A bHLH FAMA homolog (HORVU.MOREX.r3.1HG0072670) was targeted by CSEP0047; *Arabidopsis* FAMA is involved in stomatal development, ABA signaling, and resistance to bacterial and fungal pathogens (Li *et al*., 2018; Zhang *et al*., 2025). A homolog of MYC2-TARGETED BHLH1 (HORVU.MOREX.r3.4HG0402030) interacted with CSEP0047 and CSEP0062, which is involved in terminating jasmonic acid-mediated transcriptional responses in *Arabidopsis* (Liu *et al*., 2019). Additional regulators related to hormone signaling include two Auxin-Response Factors (HORVU.MOREX.r3.2HG0211750 and HORVU.MOREX.r3.6HG0566320) targeted by CSEP0004/BEC1003/BEC5, an Aux/IAA protein (HORVU.MOREX.r3.7HG0706260) targeted by CSEP0047, and a TIFY/JASMONATE ZIM (JAZ)-domain protein (HORVU.MOREX.r3.5HG0479120) targeted by CSEP0047.

Three proteins (HORVU.MOREX.r3.1HG0029350, HORVU.MOREX.r3.2HG0196160, and HORVU.MOREX.r3.2HG0163020) with a Myb/SANT putative DNA-binding (IPR024752) domain were targeted by CSEP0062, and HORVU.MOREX.r3.2HG0163020 also interacted with the related CSEP0061/BEC1048. Additionally, CSEP0062 targeted HORVU.MOREX.r3.4HG0410020 and HORVU.MOREX.r3.5HG0503060, two homologs of NODULE INCEPTION (NIN)-LIKE transcription factors critical for nitrogen signaling which have been recently shown to suppress JA-mediated defense responses in Arabidopsis and act as negative regulators of immune responses and ferroptotic cell death in rice (Chen *et al*., 2022; Pu *et al*., 2025). CSEP0047 interacted with HORVU.MOREX.r3.2HG0152890, a class II TCP. TCP family members have numerous roles in regulating plant development, cell division, hormone signaling, and are targeted by multiple pathogen effectors (Lopez *et al*., 2015). CSEP0062 interacted with HORVU.MOREX.r3.5HG0484060, a WRKY-IIa subfamily member that is related to barley WRKY1/2 and WRKY76 in rice and wheat, all of which have been reported to be negative regulators of fungal disease resistance (Shen *et al*., 2007; Liu *et al*., 2016; Kan *et al*., 2021; Jin *et al*., 2022). Collectively, these results suggest that *Bh* targets a range of transcription factors and activators/repressors with varied activities to promote susceptibility (**Table S2**).

### CSEPs target protein kinase signaling

Among the protein kinases, we found transmembrane and cytoplasmic RLKs, as well as other types of protein kinases. CSEP0372/AVR_A13_ and CSEP0069 targeted STRUBBELIG-RECEPTOR FAMILY (SRF, RLK^LRR-V^) members *Hv*SRF3_L1 (HORVU.MOREX.r3.7HG0739520) and HvSRF6/7-L1 (HORVU.MOREX.r3.4HG0405780), respectively. HvSRF3_L2 (HORVU.MOREX.r3.6HG0570440) also interacted with CSEP0372/AVR_A13_ in binary assays, although it was below our high-confidence threshold (Enrich = 0.6623). Recently, CSEP0372/AVR_A13_ was independently found to target *Hv*SRF3_L1/L2 to disrupt iron homeostasis and immunity (Shi *et al*., 2026). Increasingly, SRF3-like and SRF6-like family members are reported to regulate immune responses across dicots and monocots (Alcázar *et al*., 2010; Wang *et al*., 2018; Hernandez *et al*., 2019; Platre *et al*., 2022; Duan *et al*., 2024; Bhasin *et al*., 2025; Shi *et al*., 2026).

Notably, the related effectors CSEP0061/BEC1048 and CSEP0062 targeted the mixed lineage kinase-like (MLKL) proteins HORVU.MOREX.r3.7HG0729760 and HORVU.MOREX.r3.7HG0729840. HORVU.MOREX.r3.7HG0729760 also interacted with CSEP0174/AVR_A9_ (Enrich = 0.6499) in binary assays. MLKLs exhibit a 4-helix bundle (4HB)-kinase domain architecture and are executioners of necroptic cell death in vertebrates (Murphy, 2020). In plants, MLKLs have a critical role in immune responses to bacterial, fungal, and oomycete pathogens, where they can oligomerize to form channels at the plasma membrane to mediate calcium influx and activate cell death downstream of NLR receptors (Mahdi *et al*., 2020; Li *et al*., 2024a; Shen *et al*., 2024; He *et al*., 2024; Bennett *et al*., 2026; Asuke *et al*., 2026). HORVU.MOREX.r3.7HG0729760 is a close homolog of wheat RWT3.6.8 ASSOCIATED KINASE (R3AK) that functions with the NLR RWT3.6.8 to confer wheat blast resistance (Bennett *et al*., 2026; Asuke *et al*., 2026). To our knowledge, this is the first report of pathogen effectors interacting directly with MLKL proteins.

Additionally, we uncovered interactions between CSEP0137 and HORVU.MOREX.r3.4HG0348910, an SPARK-RLK^RKF3^ domain protein, which is related to the ARBUSCULAR RECEPTOR-LIKE KINASE 1 that regulates mycorrhizal associations in rice (Montero *et al*., 2021). CSEP0047 targeted HORVU.MOREX.r3.2HG0193090, a Dual specificity protein kinase YAK1 homolog with activities in development and abiotic stress responses (Forzani *et al*., 2019), as well as HORVU.MOREX.r3.1HG0037210, an ortholog of the Arabidopsis ENHANCED DISEASE RESISTANCE 1 kinase, which regulates resistance to fungal pathogens, including powdery mildew (Frye *et al*., 2001; Hiruma *et al*., 2011). CSEP0307 interacted with the cyclin-dependent kinase C HORVU.MOREX.r3.3HG0323020, indicating that the cell cycle is also a *Bh*-targeted process. We also uncovered an interaction between CSEP0062 and the protein phosphatase 2C HORVU.MOREX.r3.1HG0017240. Together, these results indicate that *Bh* interferes with host kinase signaling through various mechanisms to promote susceptibility (**Table S2**).

### CSEPs interact with additional proteins with putative roles in immunity

We found related CSEPs that target HMA domain proteins: CSEP0168 interacted with HORVU.MOREX.r3.2HG0174410 and CSEP0167 with HORVU.MOREX.r3.2HG0174500, two small proteins consisting of a Heavy metal-associated isoprenylated plant protein (HIPP) 47/16 domain (IPR042885). The closest rice homolog of these barley proteins is HIPP20, a susceptibility factor to blast disease that is bound and stabilized by the *M. oryzae* effector AVR-PikD (Maidment *et al*., 2021; Oikawa *et al*., 2024). CSEP0061/BEC1048 targeted *Hv*SGT1 (HORVU.MOREX.r3.3HG0273490), which complexes with the molecular chaperones HSP90 and RAR1 to stabilize NLRs, including MLA (Azevedo *et al*., 2006; Chapman *et al*., 2021, 2022). CSEP005/BEC1002 interacted with HORVU.MOREX.r3.4HG0392460, a homolog of *Arabidopsis* ARMADILLO REPEAT ONLY 2, which has been reported to be a direct activator of plant CYCLIC NUCLEOTIDE-GATED CHANNELS that have critical roles in calcium and defense signaling. We also uncovered interactions with CSEP0138 and HORVU.MOREX.r3.7HG0636480, an LRR-containing protein with no additional domains that has similar architecture to SHOC2, a scaffolding protein that acts as a positive regulator of the RAS-MAPK/ERK signaling pathway in humans (Jang *et al*., 2021). Collectively, these results demonstrate that CSEPs target a multitude of host immune processes during infection (**Table S2**).

### AVR subnetworks within the barley interactome

AVR effectors have assorted targets in various cellular compartments, such as the plasma membrane or endoplasmic reticulum (ER) (Crean *et al*., 2023; Li *et al*., 2024b; Isaksson *et al*., 2025; Shi *et al*., 2026). As shown in **Table 1**, we screened several characterized *Bh* AVR_A_ effectors and nearly all of them had unique and diverse targets, with little overlap (**Fig. S6, File S3)**.

**Table 1.** AVR-interacting proteins confirmed via binary Y2H.

| Bait | Prey annotation<sup>1</sup> | Prey HORVU/BLGH I.D. | Y2H-SCORES<br>Enrichment | Y2H<sup>2</sup><br>binary<br>score |
| --- | --- | --- | --- | --- |
| AVR<sub>A1</sub><br>(CSEP0008,<br>BEC1001)<br>(4) | 17.9 kDa class I heat shock protein | HORVU.MOREX.r3.4HG0389430 | 0.856158 | 2 |
|  | 16.9 kDa class I heat shock protein | HORVU.MOREX.r3.3HG0226130 | 0.737163 | 2 |
|  | *HvERdj3B* | HORVU1Hr1G022990<sup>3</sup> (AK376215.1 and KAE8793706.1) | N/A | 4 |
|  | Ubiquitin family protein (BAG6 homolog) | HORVU.MOREX.r3.3HG0231510 | 0.422809 | 4 |
| AVR<sub>A6</sub><br>BLGH_00708<br>(5) | NARROW LEAF 1 peptidase | HORVU.MOREX.r3.5HG0419500 | 0.976616 | 5 |
|  | Chaperone protein dnaJ | HORVU.MOREX.r3.2HG0162940 | 0.692422 | 5 |
|  | Calcium-dependent protein kinase 1 adaptor protein 2 (*HvCAP2*) | HORVU.MOREX.r3.7HG0721710 | 0.283800 | 2 |
|  | SERRATE homolog | HORVU.MOREX.r3.7HG0682160 | 0.880664 | 2 |
|  | Ubiquitin CUE | HORVU.MOREX.r3.7HG0745710 | 0.843452 | 1 |
| AVR<sub>A7-1</sub><br>CSEP0059<br>(1) | Ubiquitin family protein (BAG6 homolog) | HORVU.MOREX.r3.3HG0231510 | 0.686082 | 4 |
| AVR<sub>A9</sub><br>CSEP0174<br>(6) | AP-1 complex subunit gamma-2 | HORVU.MOREX.r3.6HG0626940 | 0.988882 | 5 |
|  | Ubiquitin family protein (BAG6 homolog) | HORVU.MOREX.r3.3HG0231510 | 0.874607 | 5 |
|  | Receptor protein kinase-like (MLKL) | HORVU.MOREX.r3.7HG0729760 | 0.649909 | 3 |
|  | COP9 signalosome complex subunit 5 | BLGH_02776-mRNA-1 | 0.592902 | 2 |
|  | AP-1 complex subunit gamma-2 | HORVU.MOREX.r3.7HG0656090 | 0.277273 | 4 |
|  | Calcium-dependent protein kinase 1 adaptor protein 2 (*HvCAP2*) | HORVU.MOREX.r3.7HG0721710 | N/A | 2 |
| AVR<sub>A13-1</sub><br>CSEP0372<br>(9) | Coatomer subunit beta | HORVU.MOREX.r3.3HG0258150 | 0.938256 | 5 |
|  | Coatomer subunit beta | HORVU.MOREX.r3.3HG0258280 | 0.951990 | 5 |
|  | E3 ubiquitin protein ligase RIN2 | HORVU.MOREX.r3.2HG0146750 | 0.959171 | 5 |
|  | Receptor-like protein kinase *HvSRF3_L1* | HORVU.MOREX.r3.7HG0739520 | 0.975643 | 5 |
|  | Protein THYLAKOID FORMATION1 chloroplastic | HORVU.MOREX.r3.2HG0135590 | 0.943858 | 5 |
|  | COP9 signalosome complex subunit 5 | BLGH_02776-mRNA-1 | 0.867123 | 5 |
|  | Calcium-dependent protein kinase 1 adaptor protein 2 (*HvCAP2*) | HORVU.MOREX.r3.7HG0721710 | 0.918156 | 5 |
|  | DWNN domain a CCHC-type zinc finger | HORVU.MOREX.r3.5HG0460350 | 0.813071 | 4 |
|  | Receptor-like protein kinase *HvSRF3_L2* | HORVU.MOREX.r3.6HG0570440 | 0.662262 | 3 |

<sup>1</sup> Order from top to bottom based on Y2H sum scores and enrichment.
<sup>2</sup> Interaction score, where 0 = no colonies present on any selection plates, 1 = colonies present on SC-LWH, 2 = colonies present on SC-LWH + 0.1mM 3AT, 3 = colonies present on SC-LWH + 0.5mM 3AT, 4 = colonies present on SC-LWH + 1.0mM 3AT, and 5 = >15 colonies present on SC-LWH + 1.0mM 3AT.
<sup>3</sup> Morex V3 gene model is mis-annotated; the Morex V1 and NCBI identifiers are listed. Published in (Li *et al.*, 2024b).

### Shared interactors of AVR_A_ and MLA: *Hv*CAP2 and CUE domain protein subnetworks

We hypothesized that some AVR_A_ targets might also interact with MLA immune receptors. These could represent cellular targets of the effector that facilitate direct interactions between the AVR and receptor, or signaling components that are engaged before or after receptor activation (Manser *et al*., 2024). One example is CALCIUM-DEPENDENT PROTEIN KINASE 1 ADAPTOR PROTEIN 2 (*Hv*CAP2, HORVU.MOREX.r3.7HG0721710), a member of the syntaxin superfamily with a coiled-coil spectrin repeat domain (Kwon *et al*., 2018). In the Enrich80 *Hv-Bh* interactome, *Hv*CAP2 interacts specifically with the barley MLA N-terminal CC domain (MLA_1-161_), CC+NB domains (MLA_1-225_), as well as AVR_A13_-1, CSEP0219, CSEP0047, and CSEP0062 (**Fig. 5, Fig. S6, File S3**). It also appears to have a weaker interaction with AVR_A6__BLGH_00708 and AVR_A9_, but not AVR_A1_ or AVR_A7_ (**Fig. 5, Table S3**).

**Fig. 5.**
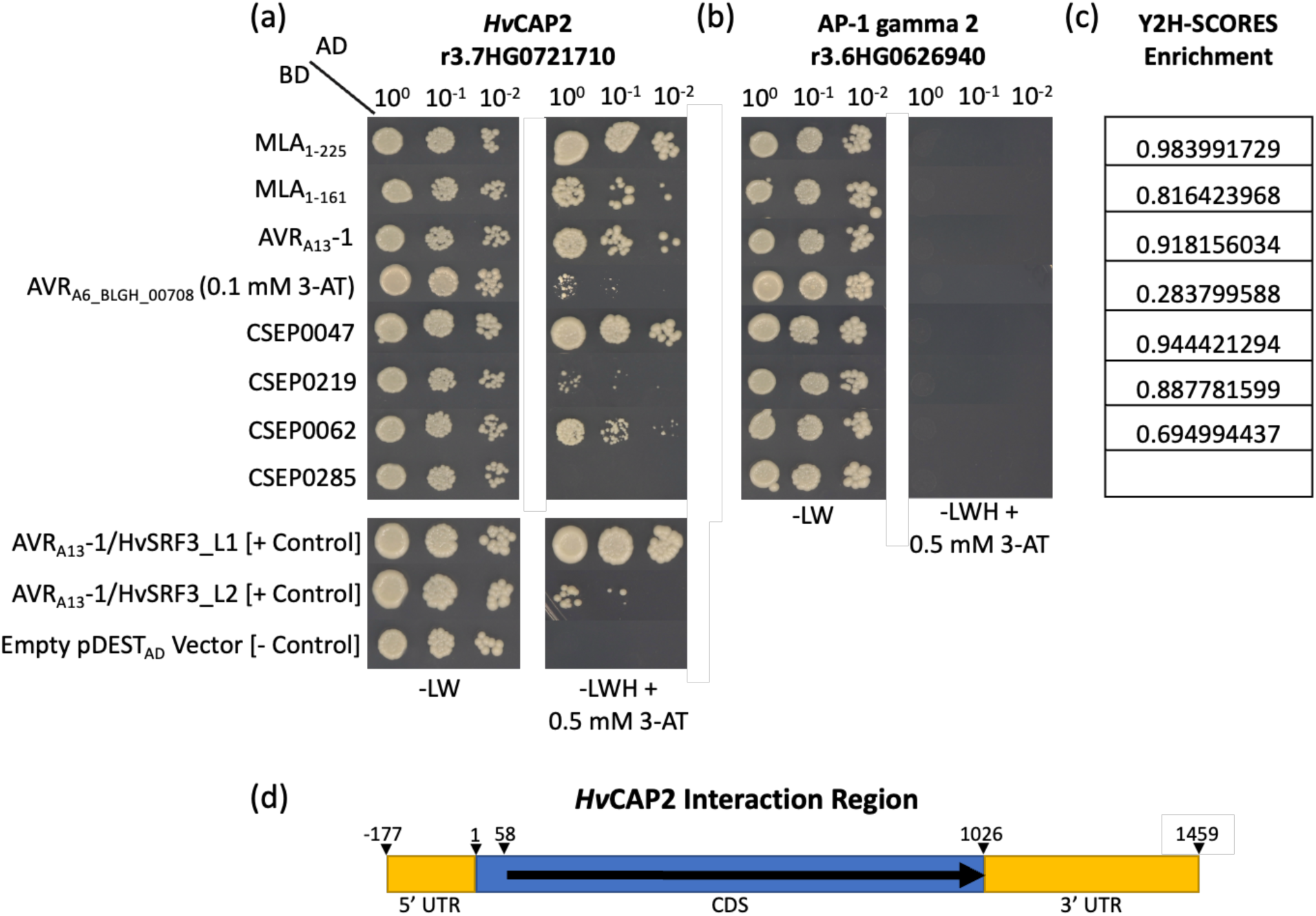
*Hv*CAP2 interacts with seven MLA, AVR and CSEP baits. (**a**) *Hv*CAP2 interactions with MLA, AVR, and CSEP baits. CSEP0285 (negative CSEP control) did not predict a Y2H interaction region in these Y2H-NGIS interactions. Shown are yeast colonies on SC-LWH + 0.5 mM 3-AT 4 days after plating, except for AVR_A6__BLGH_00708, which is shown at four days after plating with 0.1 mM 3-AT. (**b**) Prey negative specificity control HORVU.MOREX.r3.6HG0626940 is an AP-1 gamma-2 subunit known to interact with AVR_A9_ (CSEP0174) (Deb *et al*., 2026). (**c**) Y2H-SCORES enrichment score. (**d**) Diagram of *Hv*CAP2 coding sequence with interaction region predicted by NGPINT indicated by black arrow.

We showed previously that *Hv*CAP2 (formerly annotated as polyamine 1-binding protein) interacts with the highly conserved MLA_1-225_ region representing MLA1, MLA3, MLA6, MLA7, MLA8, MLA9, MLA10, MLA12, and MLA13; in addition to MLA_1-225_ orthologs *Ta*Sr33 and *Ta*MLA from wheat (*Triticum aestivum*), *Sc*Sr50 from rye (*Secale cereal*), and NLR outgroups *Hv*PBR1 and *At*ZAR1, but not *Ta*Sr22 or *Ta*Pm3a (Velásquez-Zapata *et al*., 2022). *Mc*CAP2 from the ice plant, *Mesembryanthemum crystallinum,* serves as an adaptor protein for trafficking of calcium-dependent protein kinase 1 (*Mc*CPK1) in vesicular structures and the endoplasmic reticulum (Chehab *et al*., 2007). AlphaFold2/3 models of *Hv*CAP2 indicate that it is a long coil structure, and because of their conserved nature, most MLA_1-225_ alleles and orthologs interact with it, yet only about one half of AVRs tested appear to bind (**Table S3**).

In addition to *Hv*CAP2, the only other example we identified that connects a powdery mildew AVR effector and its cognate MLA NLR receptor is a CUE domain containing protein (Coupling of Ubiquitin conjugation to ER degradation; HORVU.MOREX.r3.7HG0745710) (**Fig. 3, Fig. S6**). This CUE is quite specific and only interacts with MLA6__1-225_ and AVR_A6_ (BLGH_00708). CUE domains are ∼50 amino acid ubiquitin binding motifs found in protein trafficking, DNA repair, and targeted degradation (Shih *et al*., 2003; Kang *et al*., 2003; Bagola *et al*., 2013). Proteins containing CUE domains can function as adaptors that recruit ubiquitin-conjugating enzymes (E2s) or help target misfolded proteins (Prag *et al*., 2003). CUE domains mostly recognize ubiquitin chains, although exceptions have been reported (Lim *et al*., 2019). Like *Hv*CAP2, we have also observed that ubiquitin CUE interacts with the N-terminal MLA_1-225_ region of MLA1, MLA3, MLA6, MLA7, MLA8, MLA9, MLA10, MLA12, and MLA13; in addition to MLA_1-225_ orthologs for *Ta*Sr33 and *Ta*MLA, *Sc*Sr50, as well as *Hv*PBR1, but not *Ta*Sr22, *Ta*Pm3a, or *At*ZAR1 (Velásquez-Zapata *et al*., 2022). This current observation is the primary report of ubiquitin CUE as a link between AVR_A6_ and MLA6, however these may be interacting at different times and diverse cellular compartments. We integrated the *Hv*CAP2-MLA-AVR_A_ subnetwork with the interolog based interactome from *Hordeum vulgare,* HvInt (Velásquez-Zapata *et al*., 2022), to better visualize predicted interactions within the global barley interactome (**Fig. S7, File S4**).

### Loss-of-cell death mutations in AVR_A13_-1 delineate essential sites for interaction with coatomer β, *Hv*SRF3_L1, and *Hv*CAP2

To probe further into MLA-AVR_A_-subnetworks, we leveraged the recently reported cryo-EM structure of barley MLA13 in complex with AVR_A13_-1 (Lawson *et al*., 2025). AVR_A13_-1 associates with MLA13 C-terminal LRRs and central winged helix domain (WHD) via its basal loops, and mutations in the AVR_A13_ basal loop residues that contact MLA13 result in loss of recognition (Lawson *et al*., 2025). We synthesized alanine substitution mutants to test AVR_A13_-1 candidate interactors with the hypothesis that additional MLA-AVR_A_ interactors may associate via AVR_A13_-1 basal loops.

As illustrated in **Fig. 6**, the Y52A, G60A, and F65A loss-of cell death mutations, in addition to double and triple mutants thereof, all abolish interactions between AVR_A13_-1 and both copies of coatomer β, while the Y81A and N109A mutants still bind. These results are in direct alignment with the activation of cell death in AVR_A13_-1/MLA13 co-expression assays in *N. benthamiana* by Lawson and colleagues (Lawson *et al*., 2025) (**Fig. 6a**). The overlap in loss-of-function phenotypes suggests that these mutations are critical contact points for both a cellular target and cognate immune receptor, linking potential AVR virulence function directly to immune recognition. Further, alanine substitutions at positions Y52 and F65, as well as the Y52A/F65A double mutant, disrupt the Y2H interaction between AVR_A13_-1 and four additional barley targets; *Hv*SRF3_L1, *Hv*CAP2, DWNN CCHC-type zinc finger protein, and E3 ubiquitin protein ligase RIN2. The Y2H interaction of two others (*Bh* COP9 subunit 5 and THYLAKOID FORMATION 1 (*Hv*THF1) are partially inhibited and display varying reductions in colony growth among the Y52A and F65A single mutants, with a greater reduction in the Y52A/F65A double mutant (**Fig. 6b**). Together, these results indicate that AVR_A13_ might bind different proteins via distinct amino acid residues, and some interfaces are also important for recognition by MLA (**Fig. 6c**).

**Fig. 6.**
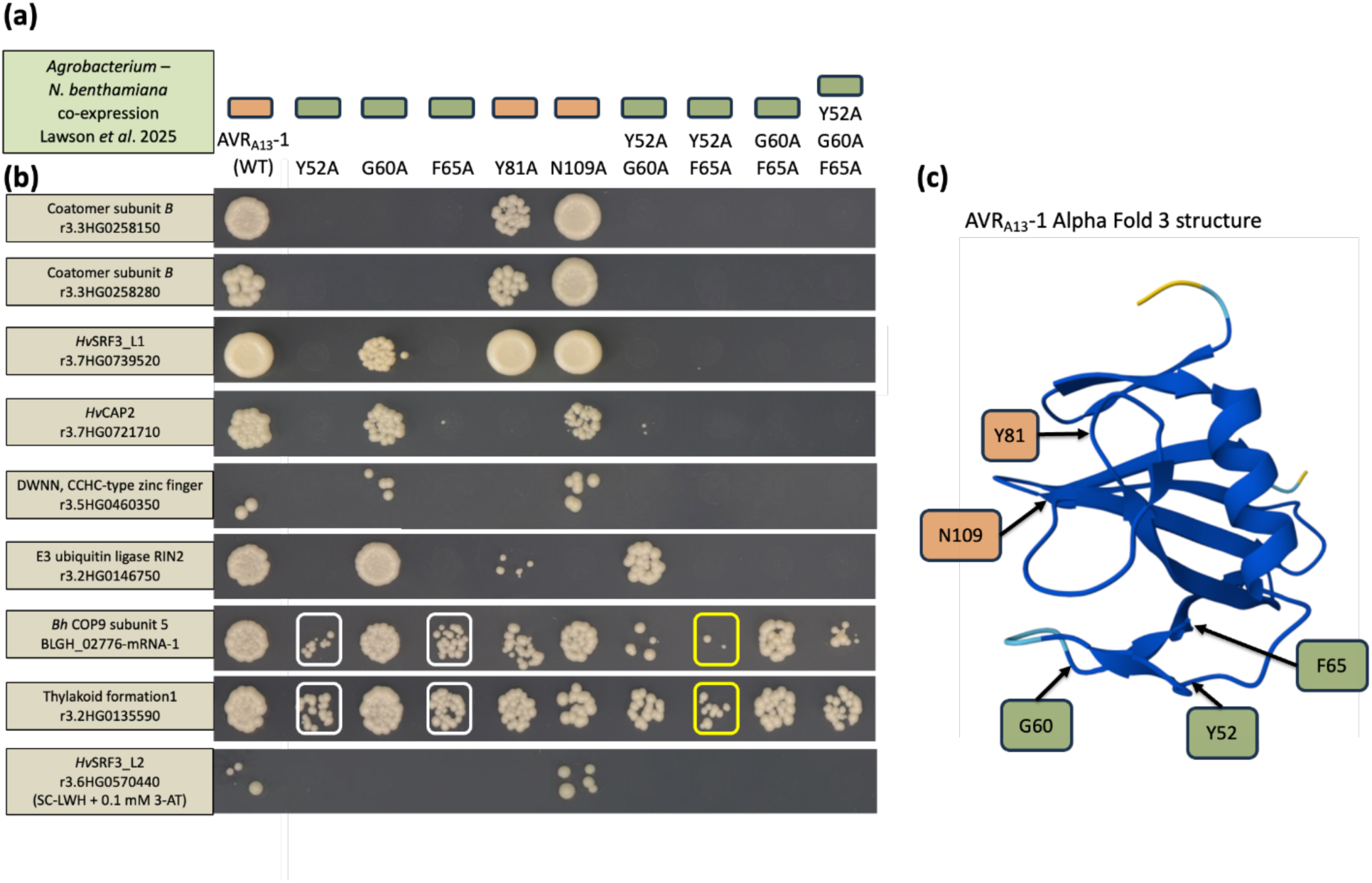
AVR_A13_-1 (CSEP0372) mutations reveal specificity of Y2H interactions. (**a**) Activation of cell death as expressed in AVR_A13_-1/MLA13 *N. benthamiana* co-expression assays. Cell death phenotypes from (Lawson *et al*., 2025) are indicated as colored boxes with tan signifying cell death (MLA13 recognition) and green signifying no cell death (loss of MLA13 recognition). (**b**) Nine verified interactors tested as preys against AVR_A13_-1 wildtype and alanine substitution mutants. Shown are yeast colonies on SC-LWH + 0.5 mM 3-AT or SC-LWH + 0.1 mM 3-AT at 4 days after plating. (**c**) AlphaFold3 prediction of AVR_A13_-1 with amino acids associated with loss (highlighted in green), or presence (highlighted in tan) of cell death in AVR_A13_-1/MLA13 *N. benthamiana* co-expression assays from (Lawson *et al*., 2025).

### Alanine mutations uncover specificity of AVR_A6_ interactions with NARROW LEAF 1 peptidase, DNAj, *Hv*CAP2, SERRATE, and ubiquitin CUE domain protein

We performed a similar substitution analysis for AVR_A6_ (BLGH_00708) and tested its candidate interactors (**Fig. 7a**). Previous work showed through CSEP0333 - AVR_A6_ domain swaps that the N-terminus comprising the β1 and α-helix, as well as the C-terminus that includes β-strands β3–β6, had no effect on recognition of MLA6. However, swapping the central segment of AVR_A6_ into CSEP0333 was sufficient to activate MLA6-mediated cell death in barley protoplasts, *i.e.*, there was a gain in effector recognition (Cao *et al*., 2023). Here, we took advantage of the AVR_A6_ amino acid residues previously identified that converted CSEP0333 into an MLA6-recognized AVR, and synthesized alanine substitutions to test for loss of interaction in comparison to native AVR_A6_ (Cao *et al*., 2023) (**Fig. 7b**). The L50A, R54A, E56A, R59A, G63A, and F70A substitutions had different effects on each of the AVR_A6_ interactors. For NARROW LEAF 1 peptidase (HORVU.MOREX.r3.5HG0419500), the R54A, L50A, E56A, and G63A substitutions mimic wild-type AVR_A6_. However, interaction was abolished or reduced dramatically for F70A and R59A, as well as the R54+F70 double mutant, and the six-alanine substitution, indicating the importance of the F70 residue and its requirement for interaction with this peptidase.

**Fig. 7.**
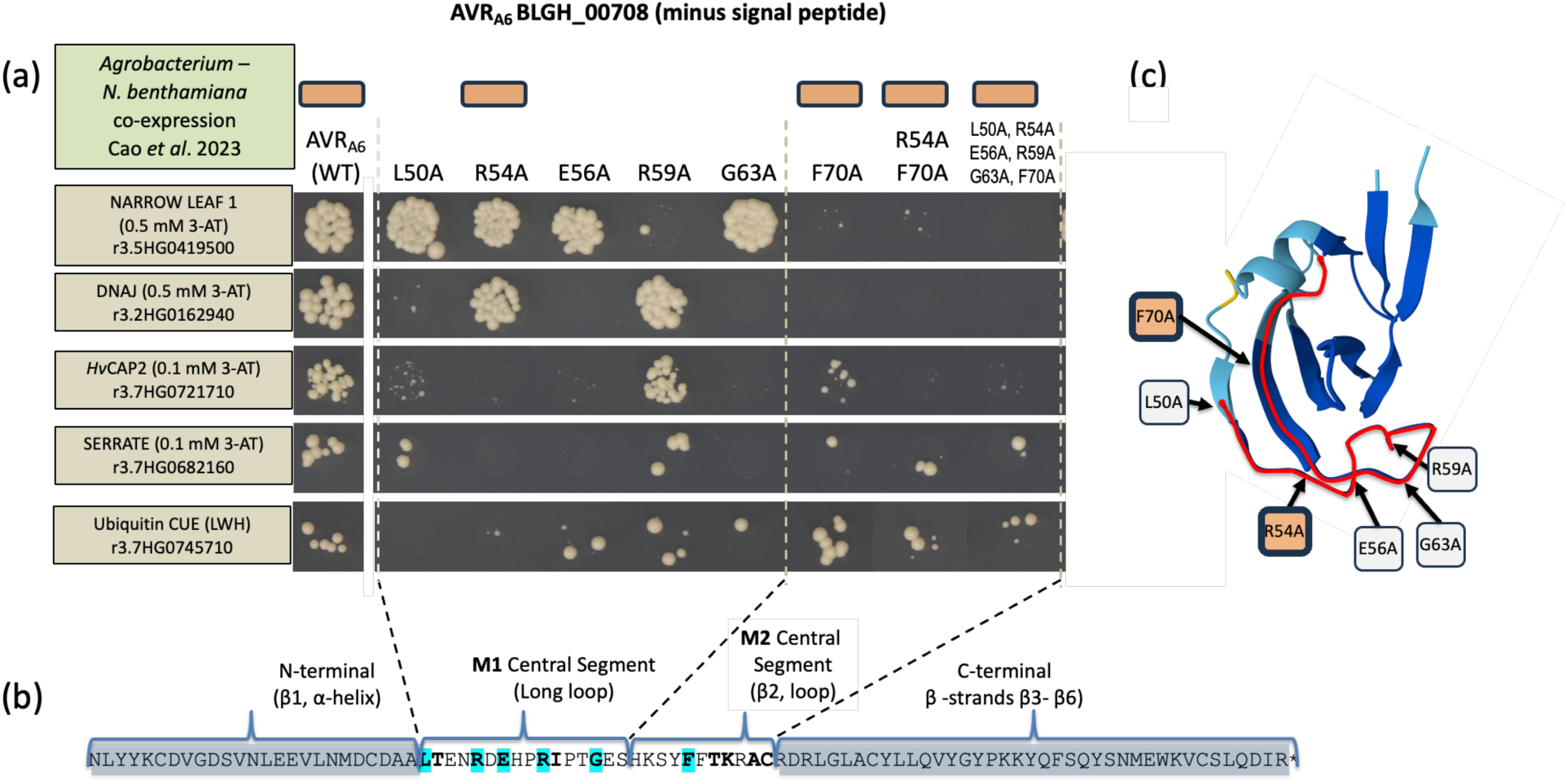
Yeast two-hybrid interactions with an AVR_A6_ (BLGH_00708) mutant panel. (a) Five verified AVR_A6_ interactors tested as preys against wild-type and alanine substitution mutants of AVR_A6_ (BLGH_00708). Shown are yeast colonies 4 days after plating on the indicated Y2H selection plates. (b) Amino acid sequence of AVR_A6_ (BLGH_00708) showing the N-terminal region, M1 and M2 central sections, and C-terminal region. Magenta-highlighted letters indicate positions of the alanine substitutions. **c)** AlphaFold3 prediction of AVR_A6_ with amino acids associated with presence (highlighted in tan) of cell death in AVR_A6_/MLA6 *N. benthamiana* co-expression assays [based on (Cao *et al*., 2023)]. The M1 and M2 central sections are highlighted by red tracing.

Likewise, the AVR_A6_ R54A and R59A mutations retained interaction with chaperone protein DNAj (HORVU.MOREX.r3.2HG0162940), whereas L50A, E56A, G63A, F70A, R54A+F70A and the six-amino acid alanine substitution was dramatically reduced or eliminated -- again indicating the importance of the F70 residue in its interaction with DNAj. Interactions with *Hv*CAP2 (HORVU.MOREX.r3.7HG0721710) were knocked down or eliminated except for R59A and less so for F70A. This may indicate that F70 is not as critical for interaction with *Hv*CAP2, although it is overcome by the R54A mutation in the R54A+F70A double mutant when interacting with *Hv*CAP2. Interestingly, the CUE domain containing protein (HORVU.MOREX.r3.7HG0745710), that interacts with both MLA6__1-225_ and AVR_A6_ (BLGH_00708), grew to similar level as compared to AVR_A6_ wildtype in the F70A substitution but was uniquely knocked out in the L50A and R54A mutants, and decreased in the R54A+F70A double mutant and the six-amino acid alanine substitution. Thus, in the case of ubiquitin CUE, R54A appears to be the more important mutation in the R54+F70 pair, analogous to *Hv*CAP2 (**Fig. 7c**). In summary, we identified AVR_A6_ mutations that differentially impact binding to host targets, and some of these correlate with MLA6 recognition (**Fig. 7**).

## Discussion

### The tip of the iceberg: networks and subnetworks in barley-powdery mildew interactions

Disease susceptibility and resistance is the culmination of a myriad of molecular interactions between pathogens and potential hosts. To better understand cellular components underlying *Blumeria* pathogenesis and barley MLA function, we generated an effector-host protein-protein interactome for nearly 10% of the annotated *Bh* CSEPs sampled across different structural subclades (**Fig. S1**). Our results suggest that manipulating host ubiquitin-associated processes, including E3 ligases, (selective) autophagy, and associated cellular trafficking, is a focal point of *Bh* effectors to neutralize plant defense (Hamada *et al*., 2026; Yuen & Bozkurt, 2026; Deb *et al*., 2026). Additionally, CSEPs are directed at transcription factors and other transcriptional regulators, likely to stabilize or antagonize their functions to shift host transcriptomes towards susceptibility. As the activity and/or abundance of many of these transcriptional regulators is known to be modulated via ubiquitination (Song *et al*., 2025), we speculate that some CSEPs might coordinate interactions between E3 ligases and host transcriptional machinery to modulate gene expression during infection. While ubiquitin- and transcription-related proteins comprise ∼30% of the PPI, we found many others that are likely to have other essential roles in powdery mildew disease outcomes, such as those involved in hormone, immune, kinase, and oxidative stress signaling, as well as metabolism, membrane transport, and vesicle trafficking (**Table S2**).

Although rigorous retesting was performed to identify a high-confidence set of interactions, a limitation of this study is that not all candidates were confirmed in binary assays, and a positive Y2H result is only one piece of evidence of a genuine PPI inside plant cells. Nonetheless, Y2H-based host-pathogen interactomes have served as a foundation for dissecting pathogenic strategies and host immune responses (Mukhtar *et al*., 2011; Weßling *et al*., 2014). It should be noted that we found many positive interactions below our enrichment score threshold, as well as many lower-scoring interactions that were well-supported by AlphaFold2-multimer analyses, indicating that there is more to discover within the larger dataset.

While it is likely that closely related CSEPs have overlapping host targets, we did not find widespread evidence for this apart from a few examples. One example is CSEP00167/CSEP0168 interactions with two HMA domain proteins, with another lower confidence CSEP0167-HMA interaction (HORVU.MOREX.r3.2HG0208690, Enrich = 0.4998). Interestingly, recent work demonstrated that the RALPH AvrWTK4 from *Blumeria graminis* f. sp. *tritici* binds to and is recognized by the integrated HMA-like domain of the wheat tandem kinase R protein WTK4 (Bernasconi *et al*., 2026). AvrWTK4 is sequence-related to *Bh* CSEP0177 and CSEP0179, which are part of a subclade of ∼20 effectors that includes CSEP0167 and CSEP0168 (**Fig. S1**), suggesting that targeting HMA domain proteins is a feature of this RALPH subfamily. Other examples include the related non-RALPHs CSEP0088/CSEP0307 targeting CONSTITUTIVE PHOTOMORPHOGENESIS PROTEIN 1 E3 ligase homolog, and CSEP0058/AVR_A6__BLGH_00708 interacting with the NARROW LEAF 1 peptidase homolog. Additionally, CSEP0061/CSEP0062 target MLKL proteins, a Myb/SANT domain protein, and a transmembrane emp24 domain-containing protein. However, as previous work has reported (Ahmed *et al*., 2015; Li *et al*., 2024b), we identified structurally divergent CSEPs that appear to converge on the same or similar host targets, with no clear pattern. The most notable example of this is the BAG6 homolog that interacted with most, but not all CSEPs, in binary tests. Given its role in autophagy and immunity, it is tempting to speculate that BAG6 might be a fundamental and redundant target of many CSEPs, although this awaits functional characterization. To address this outstanding question of host target overlap and provide clarity on how RALPH structural diversification has impacted virulence functions, this study has identified hundreds of interactions that can be utilized in all-by-all testing as suggested in (Bilstein-Schloemer *et al*., 2025).

### AVR-host target interactions linked to MLA recognition

We uncovered *Bh* AVR interactions with a diverse set of host proteins (**Table 1**). Mutational analysis of newly discovered AVR_A13_-1 interactors revealed some of these may bind through AVR_A13_-1 basal loops, similar to the AVR_A13_-MLA13 interaction region (Lawson *et al*., 2025), and what has been reported for AVR_A13_-*Hv*SRF3_L1 (Shi *et al*., 2026). Thus, recognition of AVR_A13_ by MLA13 could be directly linked to the virulence activity/host targets of the effector, and MLA13 has evolved to detect essential contact surfaces required for AVR_A13_ function. Meanwhile, some interactors of AVR_A13_ and AVR_A6_ do not appear to require the region that is recognized by the cognate MLA, suggesting multiple interfaces for different targets (**Fig. 6**, **Fig. 7**). Understanding how AVRs interact with and influence the native activities of these host proteins at a mechanistic and structural level could elucidate general principles of MLA immune receptor function.

Two paralogous coatomer β subunits associated with AVR_A13_ via its Y52, G60, and F65 residues in the RALPH basal loop region (**Fig. 6**). Intriguingly, both copies of coatomer β display significantly reduced transcript accumulation in *mla6*-m18982 susceptible mutant plants during fungal penetration at 20 HAI (**Fig. S2-22)**, as well as a significant *trans* eQTL association with the *Mla* locus at 32 HAI (q value = 7.35^E-07^, **Table S2),** suggesting a role for coatomer in powdery mildew defense (Surana *et al*., 2017). Coatomer subunit β is part of the highly conserved coat protein complex I (COPI) that assembles the coat of vesicles involved in retrograde transport from the cis-Golgi to the ER. Disruption of COPI subunits impacts Golgi structure, enhances sensitivity to abiotic stressors, and results in the accumulation of autolysosome-like structures and cell death in plants (Ahn *et al*., 2015; Sánchez-Simarro *et al*., 2020). Experiments to understand the role(s) of the COPI complex during powdery mildew infection and the consequences of AVR_A13_/β-COP interactions in plant cells could uncover fundamental aspects of cereal immune responses.

### A PPI resource for future investigations into *Blumeria*-cereal interactions

The CSEP host targets reported here unlock investigations into the specific and cooperative strategies that *Blumeria* fungi employ during infection (Li *et al*., 2024b; Deb *et al*., 2026), as well as previously unknown aspects of cereal immunity. Future work validating host targets and their role(s) in susceptibility and resistance could lead to the identification of host genes that can be engineered or targeted in breeding programs to develop crops with enhanced disease resistance.

## Supporting information

Elmore et al. Hv-Bh Interactome 2026 Supplemental

## Supporting Information

**Supporting Fig. S1** contains the All CSEP structural dendrogram.

**Supporting Fig. S2** contains time-course expression graphs of all 52 CSEP/MLA/*Hv*SRF3 baits and AVR and *Hv*CAP2 subnetwork targets.

**Supporting Fig. S3** contains an overview of HvInt integration of the Enrich80 network.

**Supporting Fig. S4** contains AlphaFold2-multimer analysis of candidate interactions.

**Supporting Fig. S5** contains InterPro domains associated with interactors for each CSEP.

**Supporting Fig. S6** contains an annotated graph of the HvCAP2-MLA-AVR subnetwork.

**Supporting Fig. S7** contains an overview of HvInt integration of the HvCAP2-MLA-AVR Y2H subnetwork.

**Supporting Table S1** contains bait and prey sequences used for Y2H binary validation among *Mla* alleles, orthologs, and outgroup NLRs.

**Supporting Table S2** contains Y2H-NGIS results with combined HORVU assembly genome ids, Y2H-SCORES, protein functional classifications, and eQTL data.

**Supporting Table S3** contains the total summary of binary tests among baits and preys, AVR mutant analyses.

**Supporting Table S4** contains the results of Alphafold2-multimer analyses.

**Supporting Table S5** contains the results of InterPro domain enrichment.

**Supporting Cytoscape File S1** contains the “Enrich80” Y2H interactome between barley and *Blumeria hordei*.

**Supporting Cytoscape File S2** contains the “Enrich80” Y2H interactome between barley and *Blumeria hordei* integrated with HvInt and literature interactors.

**Supporting Cytoscape File S3** contains the “HvCAP2-MLA-AVR” Y2H subnetwork between barley and *Blumeria hordei*.

**Supporting Cytoscape File S4** contains “HvCAP2-MLA-AVR” Y2H subnetwork between barley and *Blumeria hordei* integrated with HvInt and literature interactors.

## Data availability

All relevant data are within the manuscript and its Supporting files. R code and the ReadMe files for the NGPINT and Y2H-SCORES software are provided at the GitHub page https://github.com/Wiselab2/.

Zenodo links to NGPINT and Y2H-SCORES can be found at https://doi.org/10.5281/zenodo.15256036 and https://zenodo.org/records/18284172, respectively. Additionally, NGPINT is accessible via containers in Sylabs https://cloud.sylabs.io/library/schuyler/ngpint/ngpint and Dockerhub https://hub.docker.com/r/schuylerds/ngpint. Y2H-SCORES can be accessed via containers in Sylabs https://cloud.sylabs.io/library/schuyler/y2h-scores/y2h_scores and Dockerhub https://hub.docker.com/repository/docker/schuylerds/y2h_scores.

**RNA-Seq:** Infection-time-course RNA-Seq datasets are available in NCBI-GEO under the accession number GSE101304 (https://www.ncbi.nlm.nih.gov/gds/?term=GSE101304).

**Y2H-NGIS**: Raw Y2H-NGIS sequencing reads have been deposited at NCBI-GEO under accession numbers in **Table 2**.

**Table 2.**
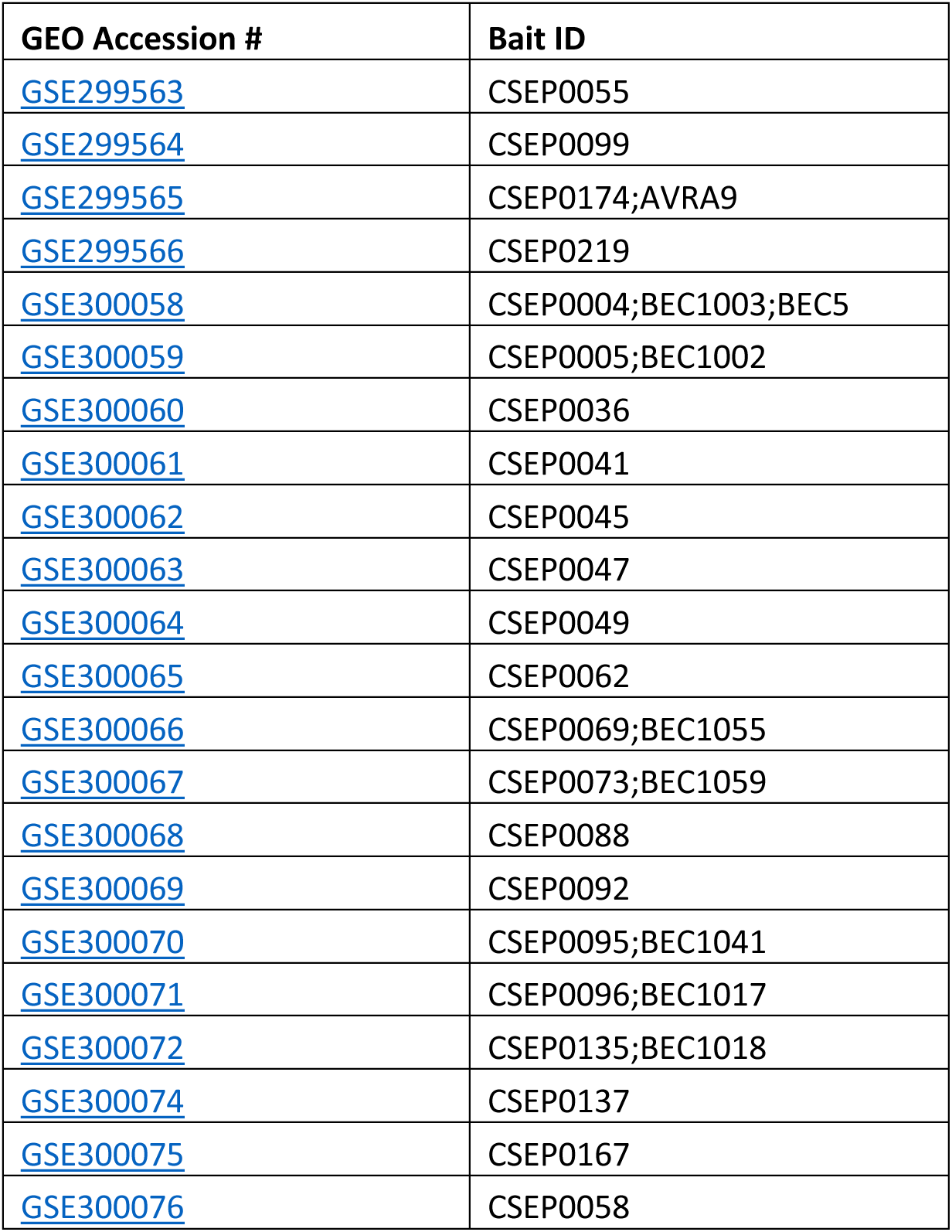
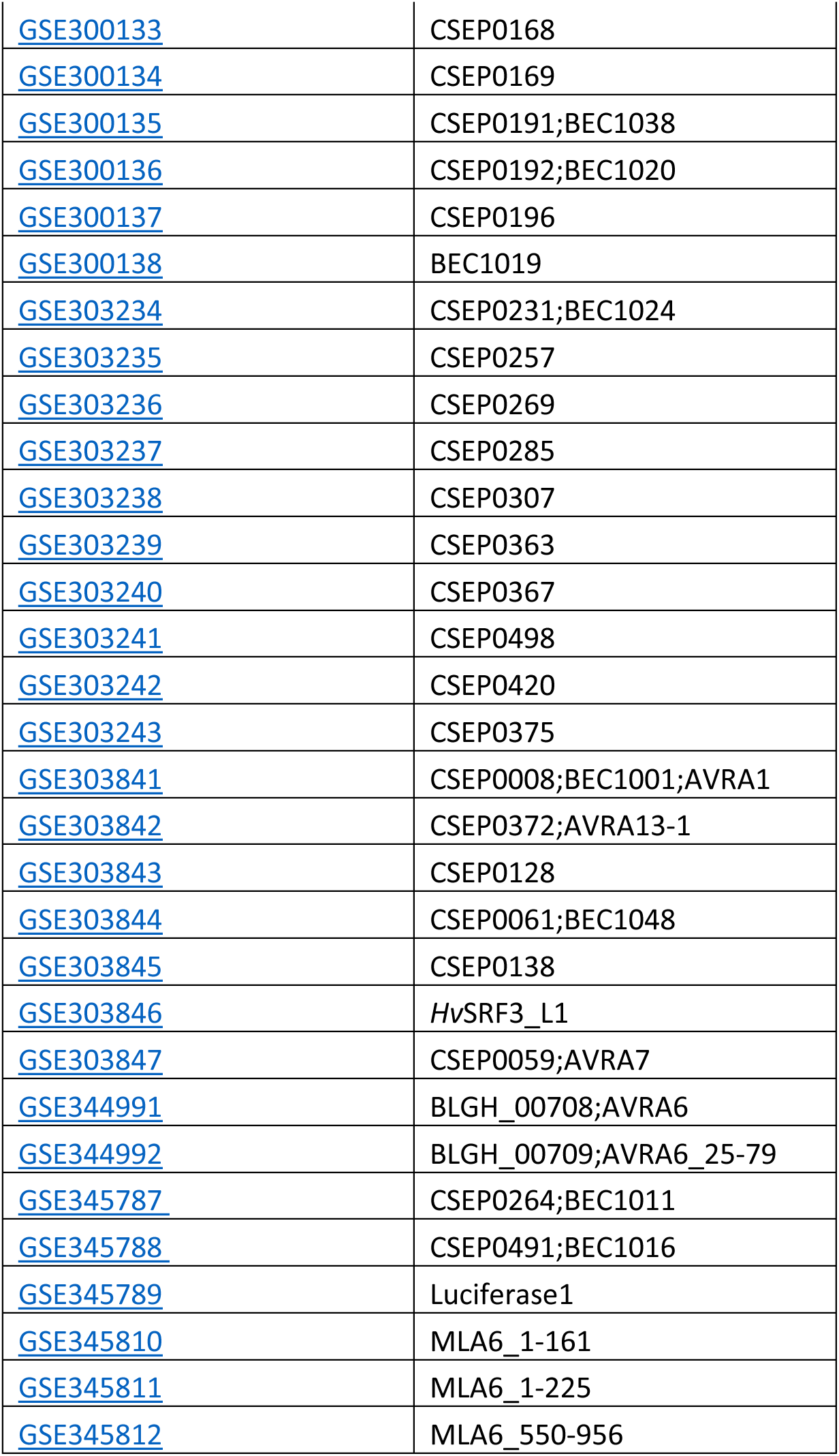
NCBI-GEO accessions for baits used in Y2H-NGIS.

## Acknowledgements

The authors thank Drs. Pietro Spanu (Imperial College, U.K.) and Ralph Panstruga (RWTH Aachen, Germany) for sharing a copy of the *Bh* CSEP clone library [under NSF-PGRP 13-39348 / ERA-CAPS ‘DURES*Trit*’], Priyanka Surana, Sagnik Banergee and Shivansh Patel for early informatics analysis, and Ana Mia Corujo, Morgan Bixby, Jessica Faust and John Higgins for invaluable lab assistance during initiation of the project. AI tools were used in the development of some of the code used to analyze data and code for figure generation, and all results were checked for accuracy.

Research supported in part by USDA-ARS Postdoctoral Research Associateship and USDA-NIFA-ELI Postdoctoral Fellowship 2017-67012-26086 to JME, Fulbright – Minciencias 2015 & Schlumberger Faculty for the Future fellowships to VVZ, Oak Ridge Institute for Science and Education (ORISE) under U.S. Department of Energy agreement number 60-5030-2-004 to SDS, and National Science Foundation – Plant Genome Research Program grant 13-39348, USDA-National Institute of Food and Agriculture grant 2020-67013-31184, USDA-Agricultural Research Service projects 3625-21000-067-00D and 5030-21220-068-000-D to RPW. Research also supported by USDA-Agricultural Research Service projects 5062-21220-024-000D to JME and 5062-21220-025-000D to MJM. This research used resources provided by the SCINet project and/or the AI Center of Excellence of the USDA-Agricultural Research Service projects 0201-88888-003-000D and 0201-88888-002-000D. The funders had no role in study design, data collection and analysis, decision to publish, or preparation of the manuscript. Mention of trade names or commercial products in this publication is solely for the purpose of providing specific information and does not imply recommendation or endorsement by the USDA, NIFA, ARS, DOE, ORAU/ORISE, or NSF. USDA is an equal opportunity provider, employer and lender.

This contribution is dedicated to the memory of our friend and colleague, Dr. Patrick Schweizer (IPK, Gatersleben, Germany), who inspired us to look beyond.

## Author Contributions

JME, VVZ, GF and RPW designed the research; JME, VVZ, GF, MM, ZG, AB and MJM performed experiments; VVZ, SS, JME and MJM contributed new analytic/computational tools. JME, VVZ, GF, MJM and RPW analyzed data. JME, VVZ, and RPW and wrote the manuscript; JME, VVZ, GF, MJM and RPW edited the manuscript.

