## Supplementary material for "Effector-host networks in the barley powdery mildew pathosystem reveal significant targeting of ubiquitin-associated proteins and transcriptional regulators": Elmore et al. Hv-Bh Interactome 2026 Supplemental

**Fig. S1** All CSEP structural dendrogram

**Fig. S2** Expression graphs rw7f: Time-course expression graphs of all 52 CSEP/MLA/*Hv*SRF3 baits and AVR and *Hv*CAP2 subnetwork targets [**appended to the end**]

**Fig. S3** Cytoscape S2_overview_Enrich80 HvInt: Overview of HvInt integration of the Enrich80 network

**Fig. S4** AlphaFold2-multimer analysis of candidate interactions

**Fig. S5** InterPro domains associated with interactors for each CSEP

**Fig. S6** HvCAP2-MLA-AVR subnetwork

**Fig. S7** Cytoscape S4_overview_HvCAP2 HvInt: Overview of HvInt integration of the HvCAP2-MLA-AVR Y2H subnetwork.

**Fig. S1** All CSEP structural dendrogram

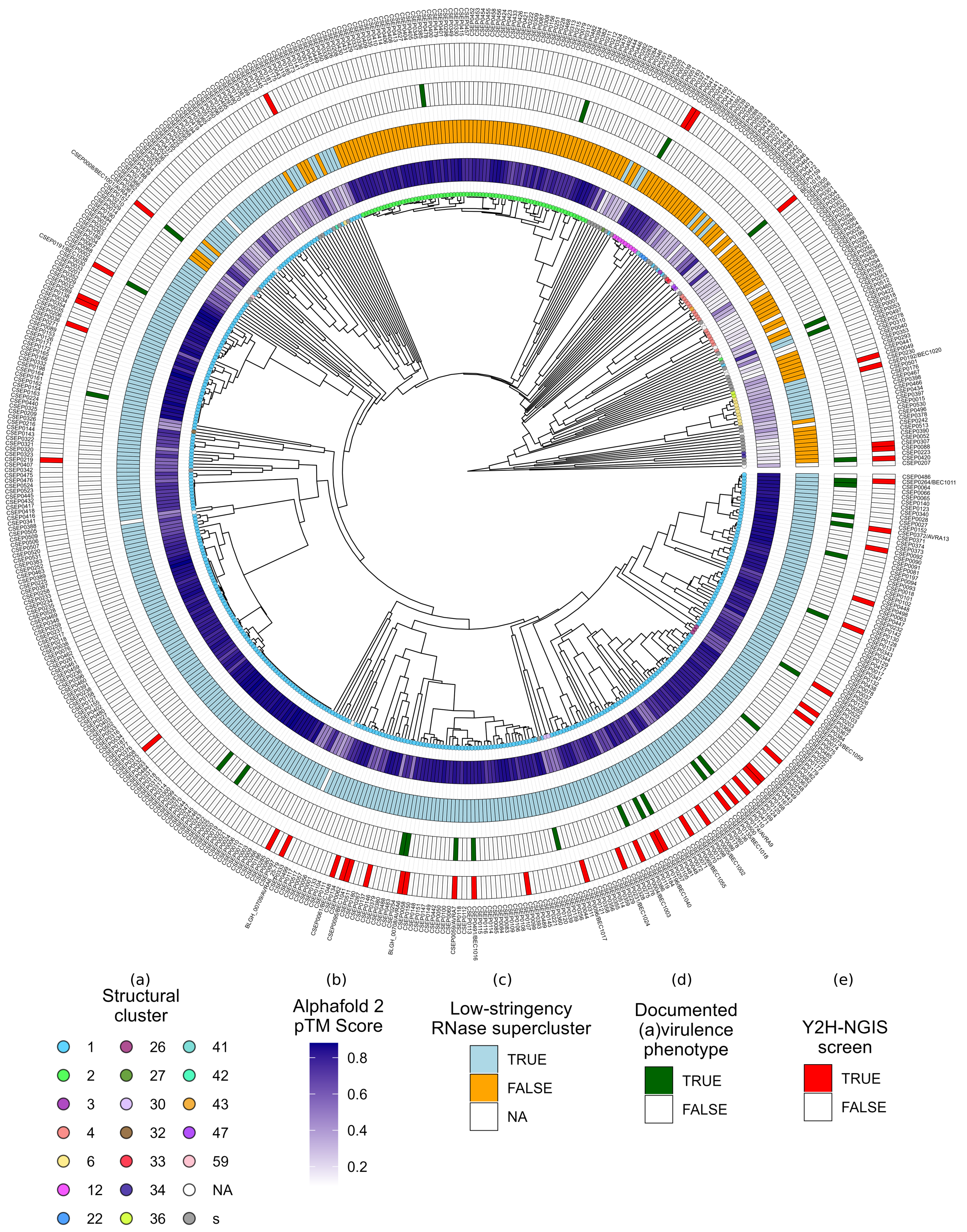

**Fig S1. Structural dendrogram of 536 *Bh* CSEPs with associated metadata**.
(a) Dendrogram based on pairwise alignments of AlphaFold2-predicted structures. Tip labels correspond to the structural family classification of (Seong & Krasileva, 2023). (b) AlphaFold2 predicted Template Modeling (pTM) score. (c) RNase-like supercluster designation from (Seong & Krasileva, 2023). (d) Documented avirulence or virulence phenotypes (Bilstein-Schloemer *et al.*, 2025). (e) Effectors screened in this study.

**Fig. S2** Time-course expression graphs of all 52 CSEP/MLA/*Hv*SRF3 baits and AVR and *Hv*CAP2 subnetwork targets

**(appended after Fig. S7)**

Time-course expression of 48 *Blumeria hordei (Bh)* effector candidates, barley MLA, receptor-like kinase *Hv*SRF3_L1, and various host targets from the barley-*Bh* interactome. RNA-Sequencing data were extracted from an infection time course of barley CI 16151 (***Mla6*; black solid line**) and the derived fast-neutron susceptible mutant (***mla6*-m18982; red dashed line**), at 0, 16, 20, 24, 32 and 48 h after inoculation with *Bh* isolate 5874 ($AVR_{a1}$, $AVR_{a3}$, $AVR_{a6}$, $AVR_{a12}$; NCBI-GEO GSE101304) and analyzed as described in (Chapman *et al*., 2021; Velásquez-Zapata *et al*., 2022). Read counts were normalized using the taxon-specific method (Klingenberg and Meinicke, 2017). Genes differentially expressed at an adjusted *p* of <0.001 for barley and <0.003 for *Bh* were considered significant (Velásquez-Zapata *et al*., 2024). Order of CSEP baits matches **Figure 1** dendogram.

**Fig. S3** Cytoscape S2_overview_Enrich80 HvInt: Overview of HvInt integration of the Enrich80 network

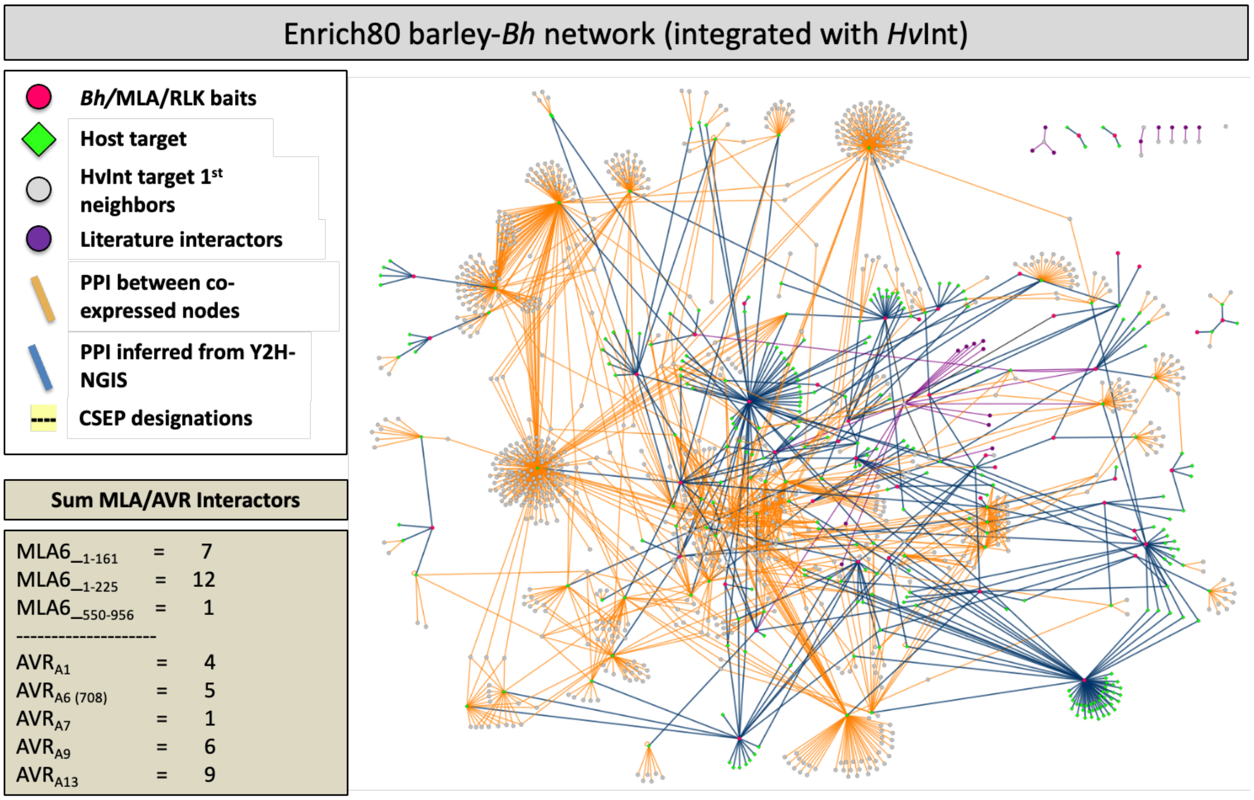


**Fig. S3. Positioning AVR/CSEP effector interactions within the barley interactome network.** Priority MLA, AVR, and CSEP baits were graphed to better visualize predicted interactions within the *Blumeria*-barley interactome. Nodes and edges were visualized in Cytoscape 3.10.4 with Y2H-SCORES enrichment score cutoffs of 0.80, 0.85, and 0.90 (**Cytoscape File S2**). Shown is a graph representation of an enrichment score cutoff of 0.80; this equates to an 88% retest rate for preys above the threshold of interaction, and 56% below. Red-filled circles designate *Hv*MLA or *Bh* effector baits, green-filled diamonds signify host targets, grey-filled circles are 1$^{st}$ neighbor target interactors, blue solid lines represent PPI inferred from 53 Y2H-NGIS screens in the current report and tan solid lines represent PPI between co-expressed nodes in the interolog-based barley interactome, *Hv*Int, comprising 66,133 edges and 7,181 nodes (Velásquez-Zapata et al. 2022). Purple-filled circles with black connecting lines are key *Bh* effector-host target interactions added into the interactome from published literature.

**Fig. S4** AlphaFold2-multimer analysis of candidate interactions

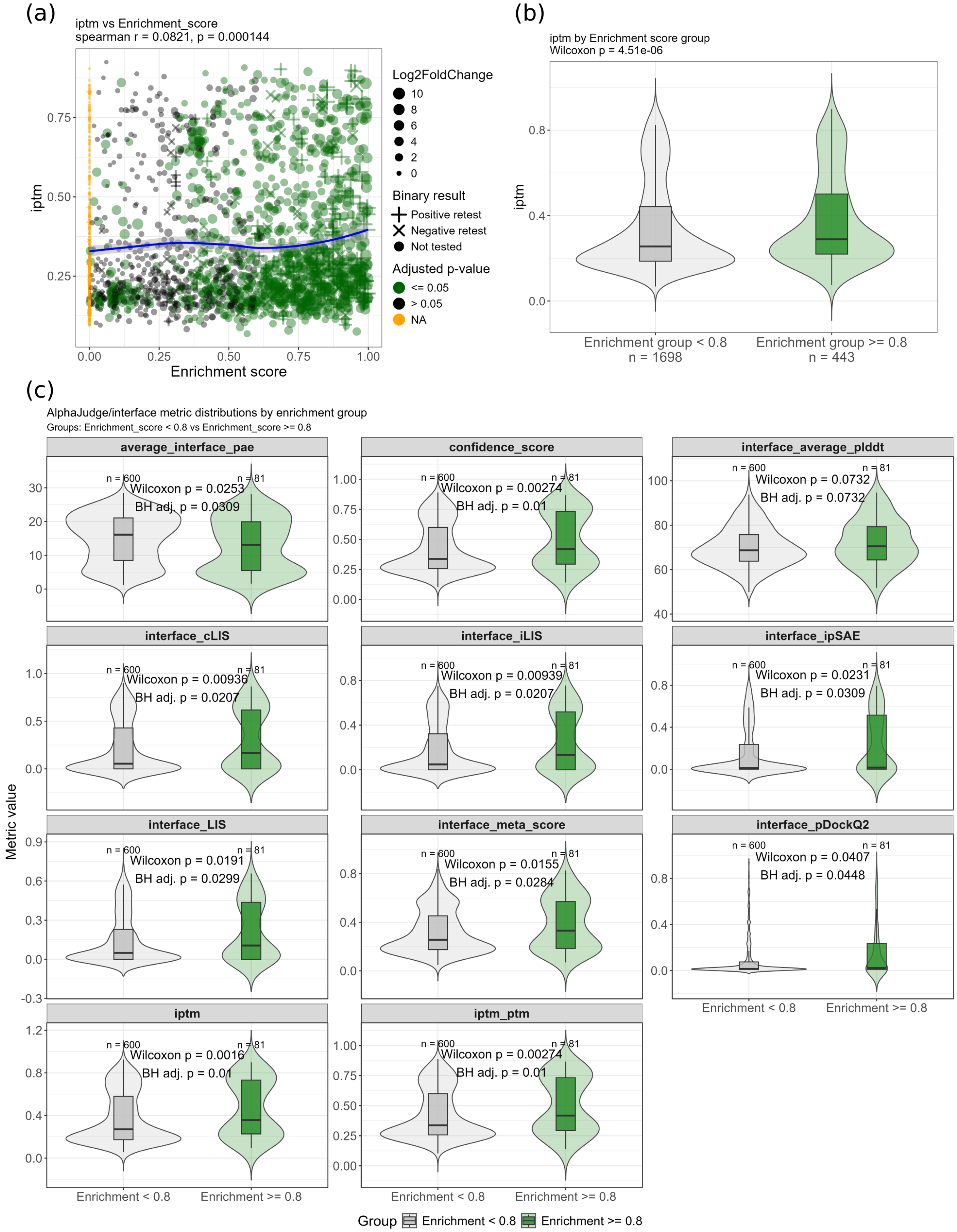

**Fig. S4. Structural modeling of candidate interactions.**

(a) Correlation of enrichment score with AlphaFold2-multimer interface predicted Template modeling (ipTM) score for all modeled interactions (n = 2141). Interactions with an enrichment of 0 were randomly selected to serve as a negative control. Point size represents the log2 fold change and color indicates the adj. p-value for statistical enrichment relative to the Y2H library controls. Point shape indicates the result from binary retesting for each interaction: "+", positive retest; "X", negative retest; "o", not tested. (b) Violin plot of the distribution of ipTM scores in the Enrich80 and Enrichment < 0.80 sets for all modeled interactions (n = 2141). Differences in the distributions between the two sets were assessed with a Wilcoxon rank-sum test. (c). Violin plots of the distributions of ipTM and additional interface confidence metrics from AlphaJudge (see Materials and Methods) for interactions where both chains were predicted with a mean predicted Local Distance Difference Test (pLDDT) score >= 0.65 (n = 681). Differences in the distributions between the two sets were assessed with a Wilcoxon rank-sum test with Benjamini-Hochberg multiple test correction.

**Fig. S5** InterPro domains associated with interactors for each CSEP

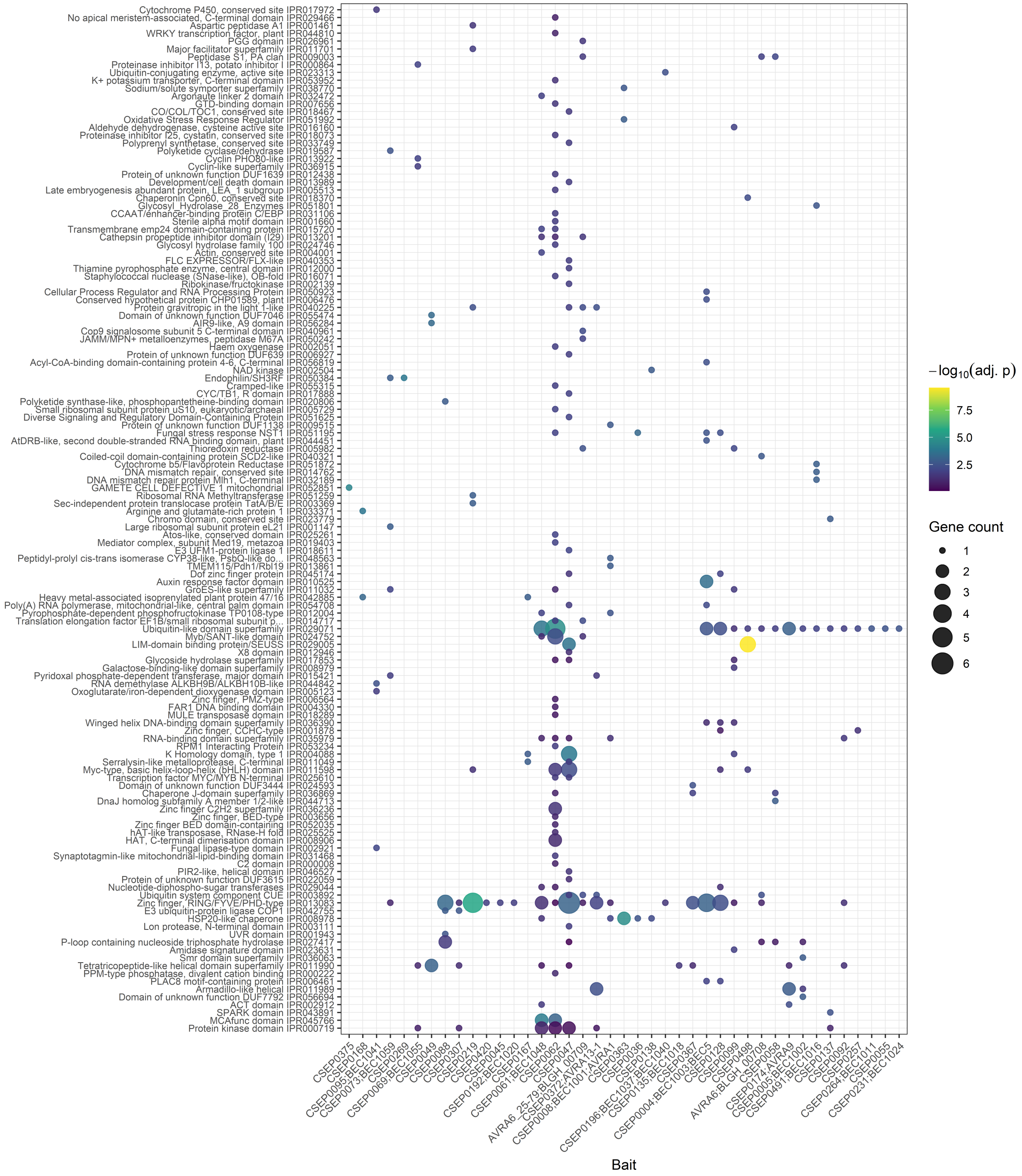


**Fig. S5. CSEPs target proteins with a diverse set of domains.**

Bubble plot depicting the enrichment of InterPro protein domains associated with proteins targeted by each CSEP bait. Data points are colored according to the adj. p-value (one-sided Fisher's exact test, Benjamini-Hochberg correction) and sized relative to the number of genes associated with each term. Interpro terms were reduced based on the proteome-wide overlap in genes associated with each domain. Terms on the y-axis are clustered based on the similarity in protein sets associated with each domain. CSEPs on the x-axis were clustered based on the overlap in their protein targets.

**Fig. S6** Annotated graph illustration of the *Hv*CAP2-MLA-AVR subnetwork

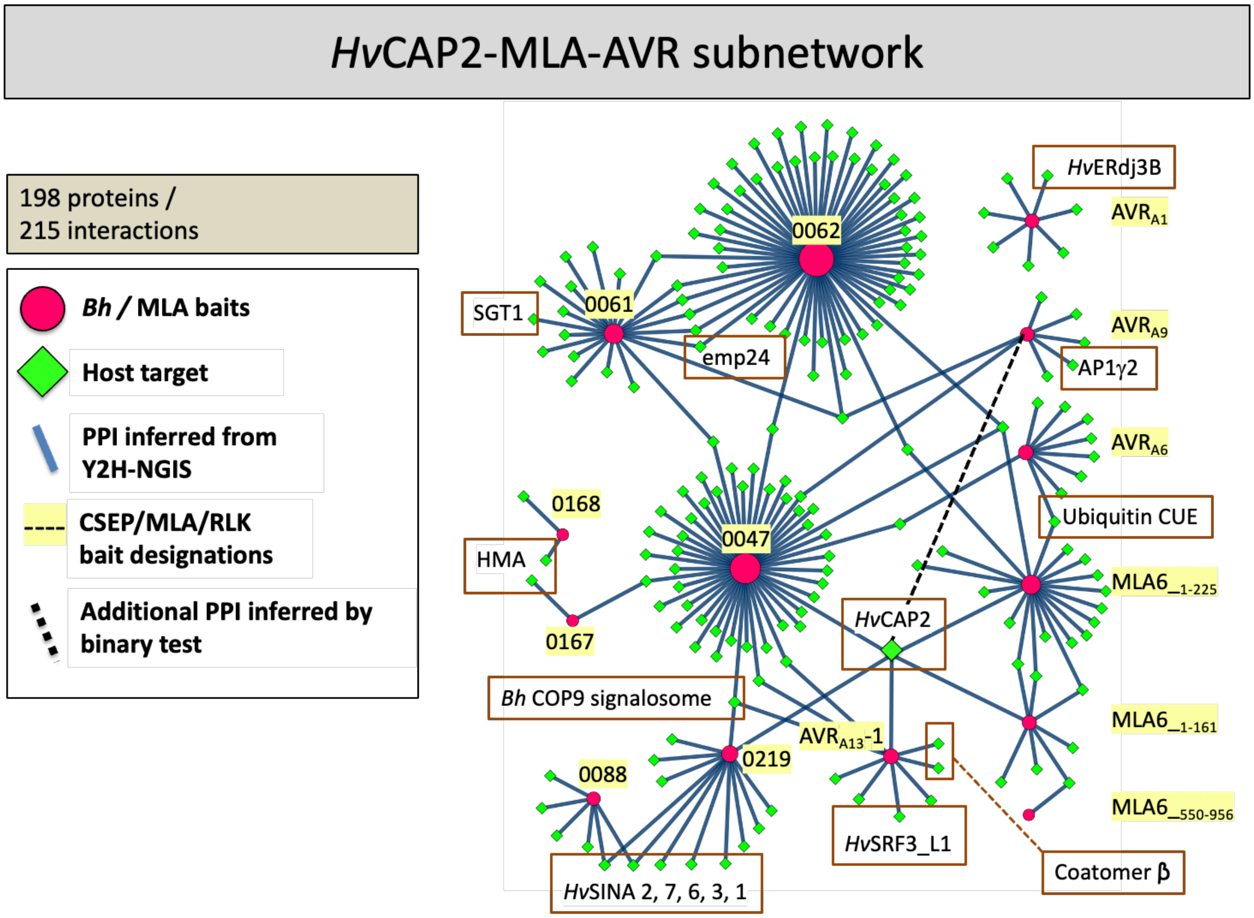


**Fig. S6. Positioning AVR effector interactions within the *Hv*CAP2 subnetwork.**
Priority MLA, AVR, and CSEP baits were subset to better visualize predicted interactions within the *Blumeria*-barley interactome. Nodes and edges were visualized in Cytoscape 3.10.4 with an Y2H-SCORES enrichment score cutoff of 0.80 (**Cytoscape File S3**). Red-filled circles designate *Hv*MLA or *Bh* effector baits (yellow highlight), green-filled diamonds signify host targets (target interactors with annotations are boxed with brown borders), blue solid lines represent PPI inferred from 52 Y2H-NGIS screens in the current report. Barley host annotations derived from Morex V3 (Mascher et al. 2022). Although *Hv*CAP2 - $AVR_{A9}$ interactions were not uncovered through Y2H-SCORES analyses, they did interact when we tested all AVRs in a binary test.

**Fig. S7** Cytoscape S4_overview_HvCAP2 HvInt: Overview of HvInt integration of the HvCAP2-MLA-AVR Y2H subnetwork.

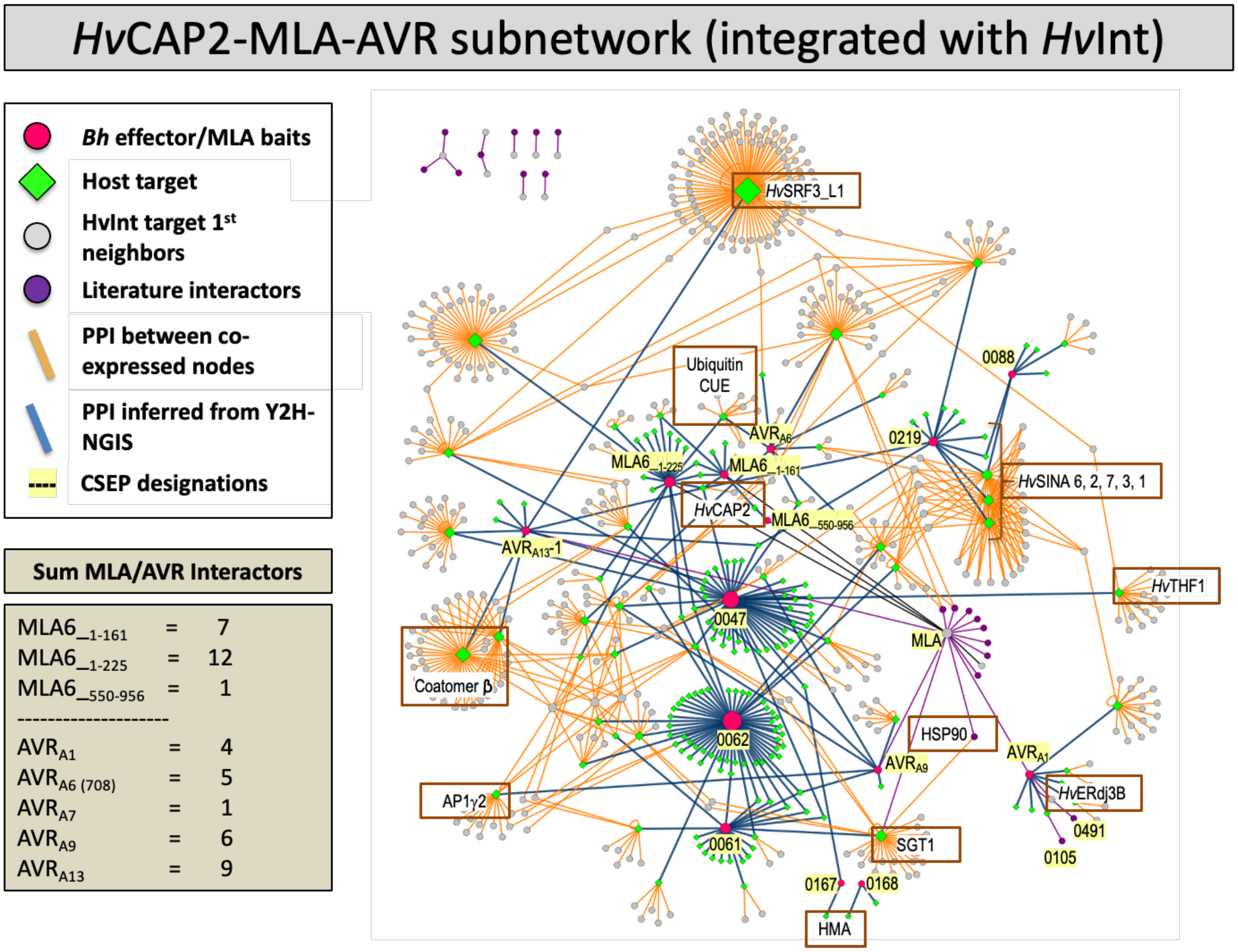


**Fig. S7. Positioning AVR effector interactions within the barley interactome network.** Priority MLA, AVR, and CSEP baits were subset to better visualize predicted immune interactions within the *Blumeria*-barley interactome. 674 nodes and 868 edges were visualized in Cytoscape 3.10.4 with a 'Y2H-Scores' enrichment cutoff of 0.8 (**Cytoscape File S4**). Red-filled circles designate *Hv*MLA or *Bh* effector baits, green-filled diamonds signify host targets, grey-filled circles are 1$^{st}$ neighbor target interactors, blue solid lines represent PPI inferred from 52 Y2H-NGIS screens in the current report and tan solid lines represent PPI between co-expressed nodes in HvInt, comprising 66,133 edges and 7,181 nodes. Purple-filled circles with black connecting lines are key *Bh* effector-host target interactions added into the interactome from published literature.

### Elmore et al. Supporting Information Fig. S2

Time-course expression of 48 *Blumeria hordei (Bh)* effector candidates, barley MLA, receptor-like kinase *Hv*SRF3_L1 (RLK5), and various host targets from the barley-*Bh* interactome. RNA-Sequencing data were extracted from an infection time course of barley CI 16151 (***Mla6*; black solid line**) and the derived fast-neutron susceptible mutant (***mla6*-m18982; red dashed line**), at 0, 16, 20, 24, 32 and 48 h after inoculation with *Bh* isolate 5874 ($AVR_{a1}$, $AVR_{a3}$, $AVR_{a6}$, $AVR_{a12}$; NCBI-GEO GSE101304) and analyzed as described in (Chapman *et al*., 2021; Velásquez-Zapata *et al*., 2022). Read counts were normalized using the taxon-specific method (Klingenberg and Meinicke, 2017). Genes differentially expressed at an adjusted $p$ of <0.001 for barley and <0.003 for *Bh* were considered significant (Velásquez-Zapata *et al*., 2024). Order of CSEP baits matches Figure 1 dendogram.

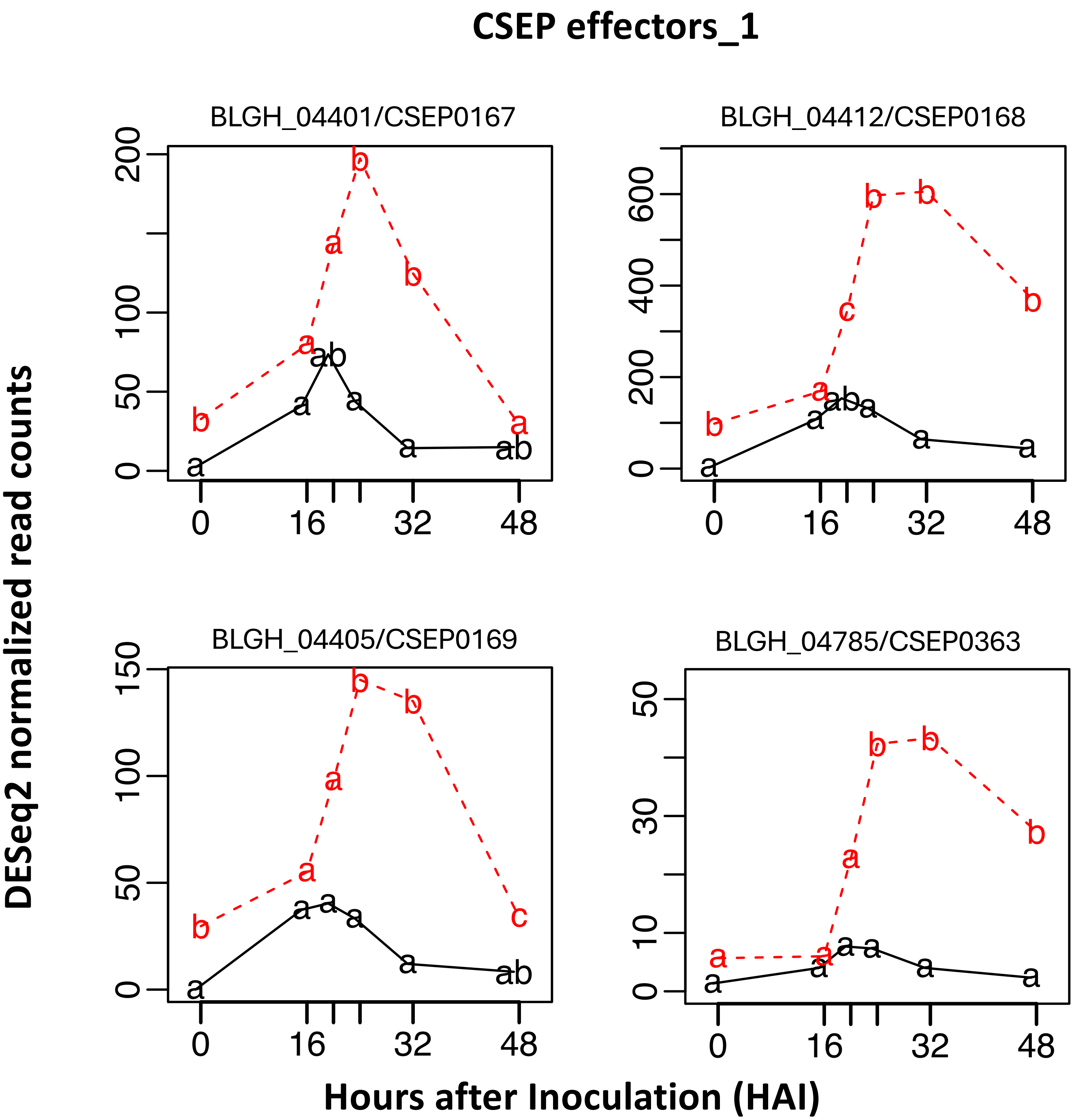


**Supporting Fig. S2-1.** Time-course expression of 48 *Blumeria hordei (Bh)* effector candidates, barley MLA, receptor-like kinase *Hv*SRF3_L1 (RLK5), and various host targets from the barley-*Bh* interactome. RNA-Sequencing data were extracted from an infection time course of barley CI 16151 (***Mla6*; black solid line**) and the derived fast-neutron susceptible mutant (***mla6*-m18982; red dashed line**), at 0, 16, 20, 24, 32 and 48 h after inoculation with *Bh* isolate 5874 ($AVR_{a1}$, $AVR_{a3}$, $AVR_{a6}$, $AVR_{a12}$; NCBI-GEO GSE101304) and analyzed as described in (Chapman *et al.*, 2021; Velásquez-Zapata *et al.*, 2022). Read counts were normalized using the taxon-specific method (Klingenberg and Meinicke, 2017). Genes differentially expressed at an adjusted *p* of <0.001 for barley and <0.003 for *Bh* were considered significant (Velásquez-Zapata *et al.*, 2024). Order of CSEP baits matches Figure 1 dendogram.

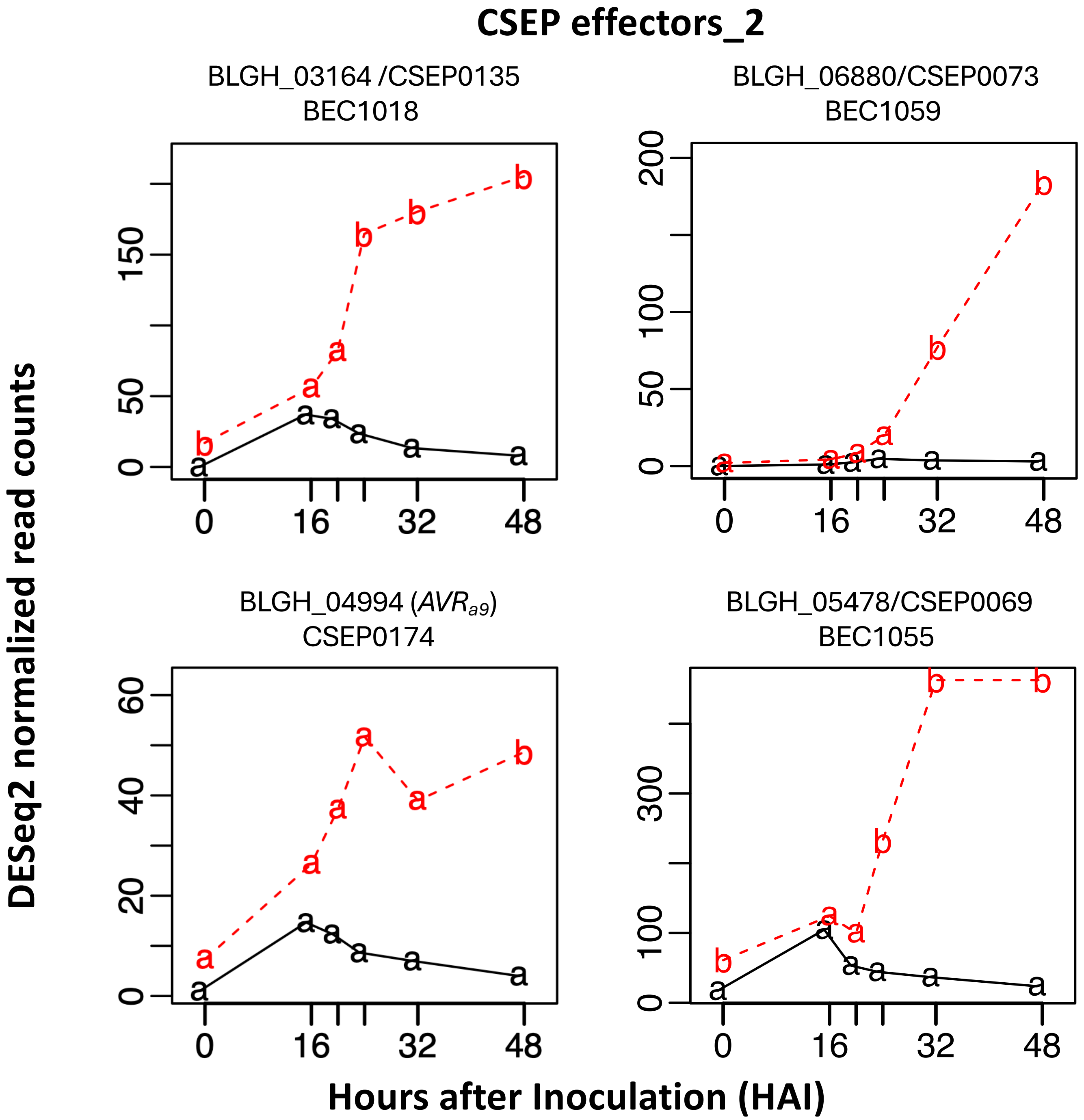


**Supporting Fig. S2-2.** Time-course expression of 48 *Blumeria hordei (Bh)* effector candidates, barley MLA, receptor-like kinase *Hv*SRF3_L1 (RLK5), and various host targets from the barley-*Bh* interactome. RNA-Sequencing data were extracted from an infection time course of barley CI 16151 (***Mla6*; black solid line**) and the derived fast-neutron susceptible mutant (***mla6*-m18982; red dashed line**), at 0, 16, 20, 24, 32 and 48 h after inoculation with *Bh* isolate 5874 ($AVR_{a1}$, $AVR_{a3}$, $AVR_{a6}$, $AVR_{a12}$; NCBI-GEO GSE101304) and analyzed as described in (Chapman *et al*., 2021; Velásquez-Zapata *et al*., 2022). Read counts were normalized using the taxon-specific method (Klingenberg and Meinicke, 2017). Genes differentially expressed at an adjusted *p* of <0.001 for barley and <0.003 for *Bh* were considered significant (Velásquez-Zapata *et al*., 2024). Order of CSEP baits matches Figure 1 dendogram.

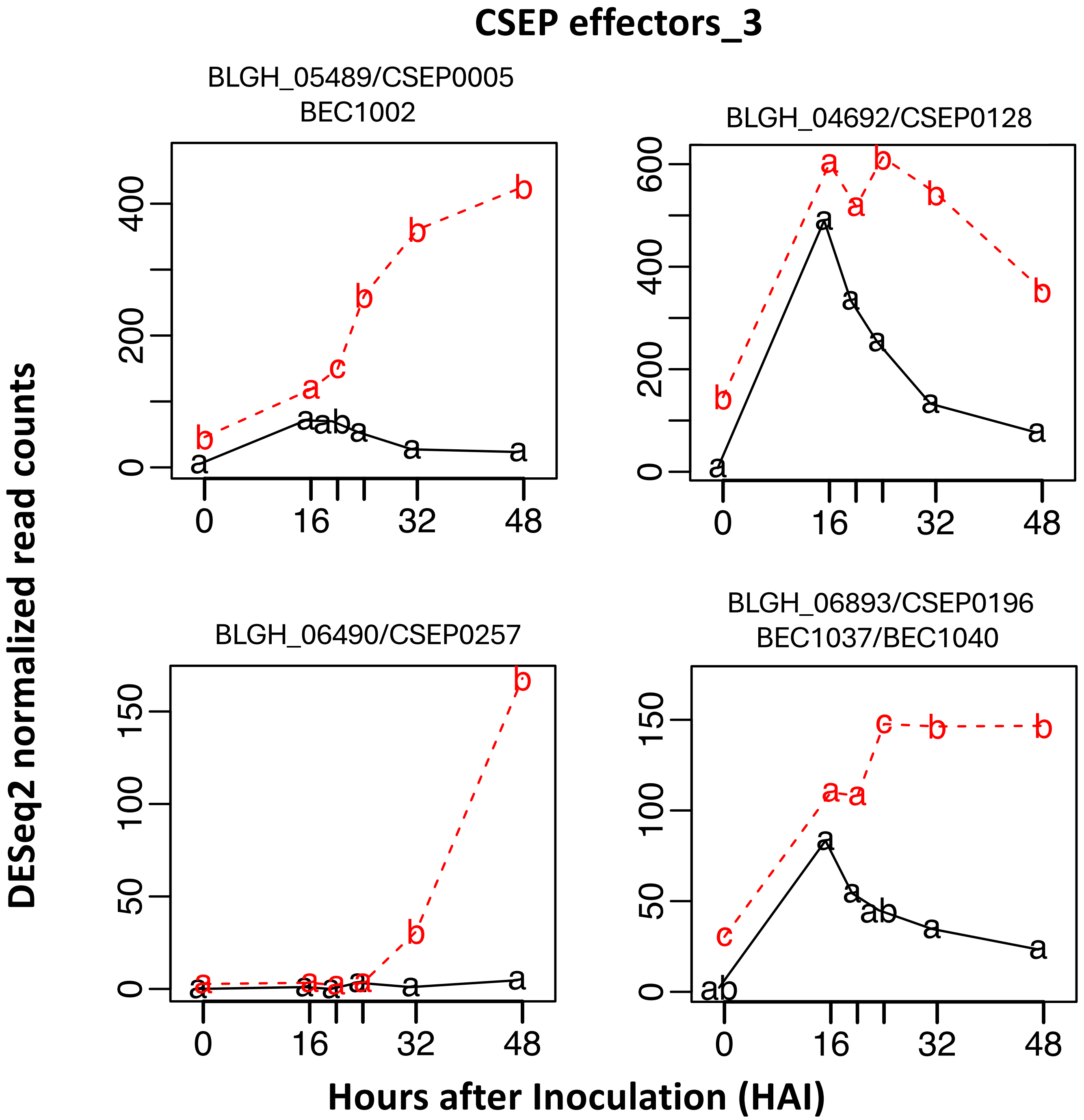


**Supporting Fig. S2-3.** Time-course expression of 48 *Blumeria hordei (Bh)* effector candidates, barley MLA, receptor-like kinase *Hv*SRF3_L1 (RLK5), and various host targets from the barley-*Bh* interactome. RNA-Sequencing data were extracted from an infection time course of barley CI 16151 (***Mla6*; black solid line**) and the derived fast-neutron susceptible mutant (***mla6*-m18982; red dashed line**), at 0, 16, 20, 24, 32 and 48 h after inoculation with *Bh* isolate 5874 ($AVR_{a1}$, $AVR_{a3}$, $AVR_{a6}$, $AVR_{a12}$; NCBI-GEO GSE101304) and analyzed as described in (Chapman *et al.*, 2021; Velásquez-Zapata *et al.*, 2022). Read counts were normalized using the taxon-specific method (Klingenberg and Meinicke, 2017). Genes differentially expressed at an adjusted *p* of <0.001 for barley and <0.003 for *Bh* were considered significant (Velásquez-Zapata *et al.*, 2024). Order of CSEP baits matches Figure 1 dendogram.

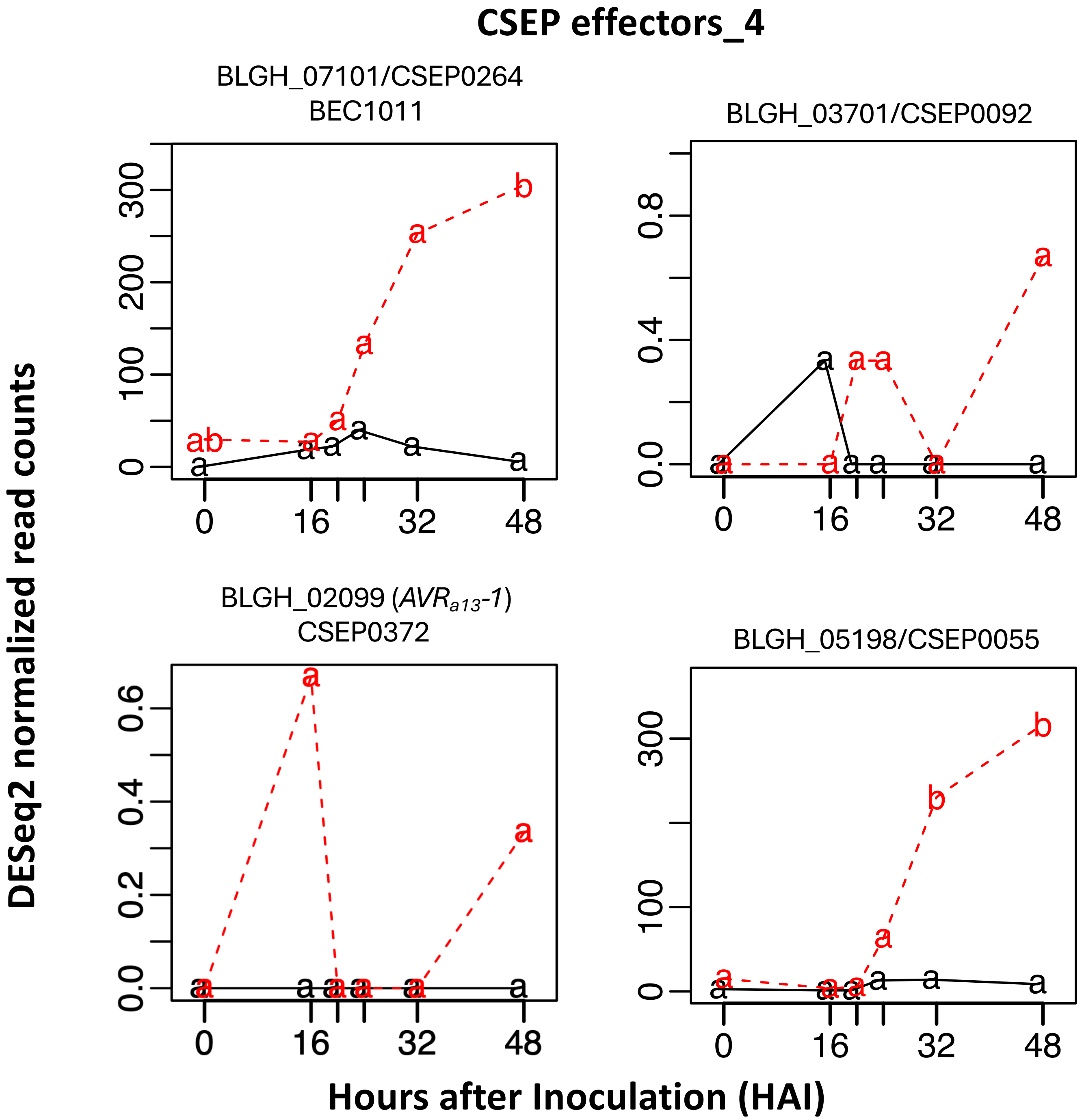


**Supporting Fig. S2-4.** Time-course expression of 48 *Blumeria hordei (Bh)* effector candidates, barley MLA, receptor-like kinase *Hv*SRF3_L1 (RLK5), and various host targets from the barley-*Bh* interactome. RNA-Sequencing data were extracted from an infection time course of barley CI 16151 (***Mla6*; black solid line**) and the derived fast-neutron susceptible mutant (***mla6*-m18982; red dashed line**), at 0, 16, 20, 24, 32 and 48 h after inoculation with *Bh* isolate 5874 (*AVR$_{a1}$, AVR$_{a3}$, AVR$_{a6}$, AVR$_{a12}$*; NCBI-GEO GSE101304) and analyzed as described in (Chapman *et al.*, 2021; Velásquez-Zapata *et al.*, 2022). Read counts were normalized using the taxon-specific method (Klingenberg and Meinicke, 2017). Genes differentially expressed at an adjusted *p* of <0.001 for barley and <0.003 for *Bh* were considered significant (Velásquez-Zapata *et al.*, 2024). Order of CSEP baits matches Figure 1 dendogram.

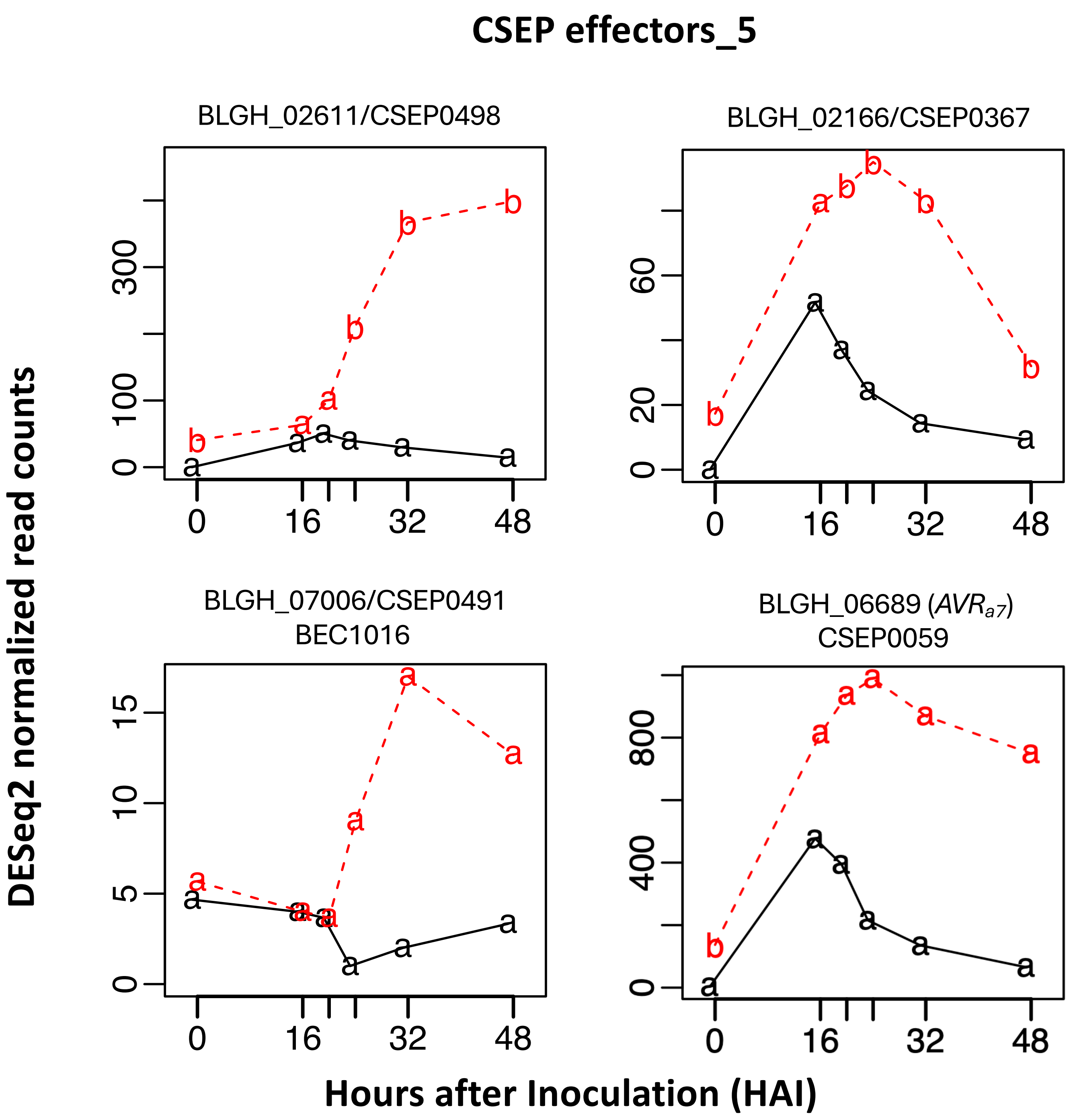


**Supporting Fig. S2-5.** Time-course expression of 48 *Blumeria hordei (Bh)* effector candidates, barley MLA, receptor-like kinase *Hv*SRF3_L1 (RLK5), and various host targets from the barley-*Bh* interactome. RNA-Sequencing data were extracted from an infection time course of barley CI 16151 (***Mla6*; black solid line**) and the derived fast-neutron susceptible mutant (***mla6*-m18982; red dashed line**), at 0, 16, 20, 24, 32 and 48 h after inoculation with *Bh* isolate 5874 ($AVR_{a1}$, $AVR_{a3}$, $AVR_{a6}$, $AVR_{a12}$; NCBI-GEO GSE101304) and analyzed as described in (Chapman *et al.*, 2021; Velásquez-Zapata *et al.*, 2022). Read counts were normalized using the taxon-specific method (Klingenberg and Meinicke, 2017). Genes differentially expressed at an adjusted *p* of <0.001 for barley and <0.003 for *Bh* were considered significant (Velásquez-Zapata *et al.*, 2024). Order of CSEP baits matches Figure 1 dendogram.

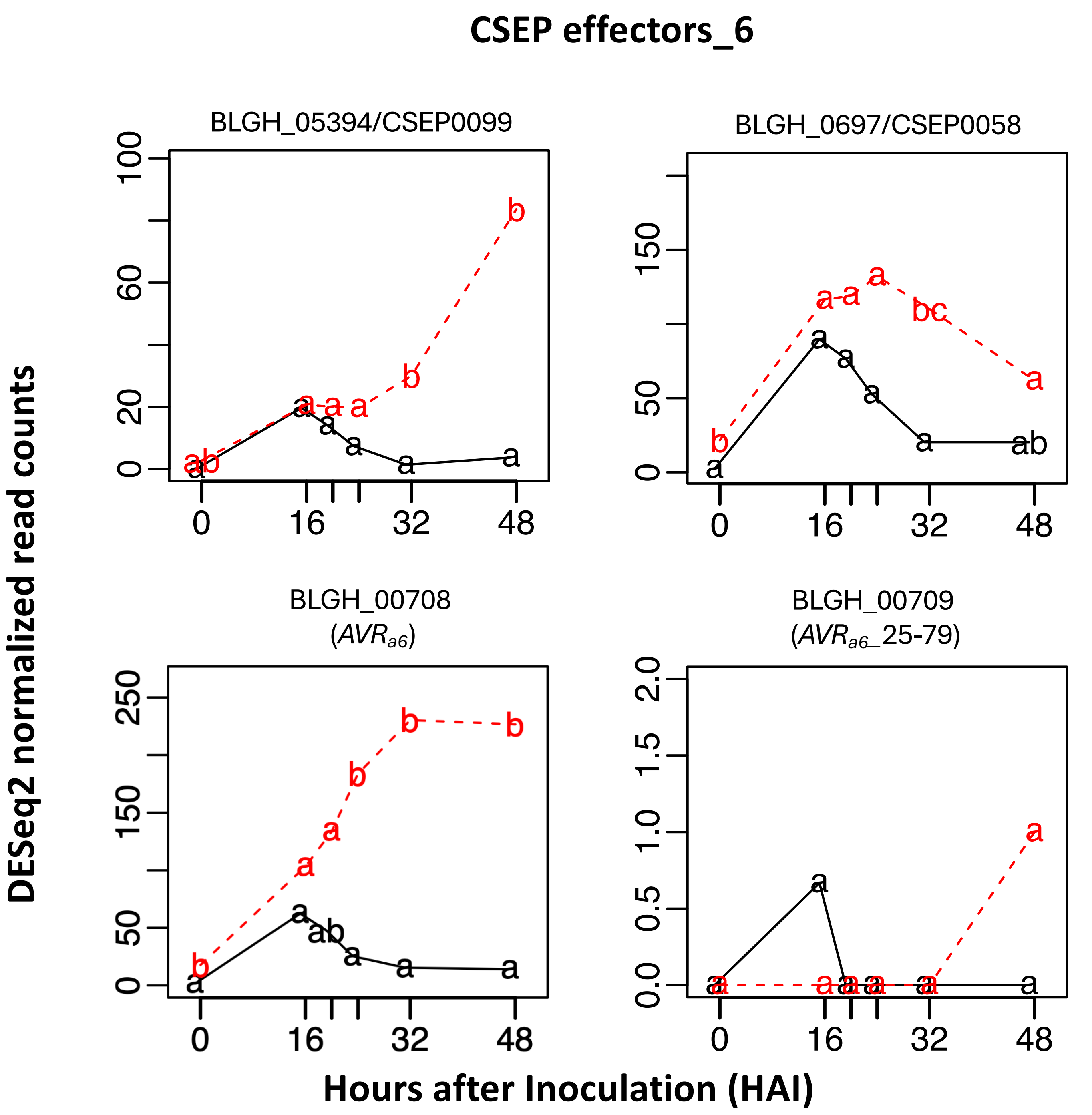


**Supporting Fig. S2-6.** Time-course expression of 48 *Blumeria hordei (Bh)* effector candidates, barley MLA, receptor-like kinase *Hv*SRF3_L1 (RLK5), and various host targets from the barley-*Bh* interactome. RNA-Sequencing data were extracted from an infection time course of barley CI 16151 (***Mla6*; black solid line**) and the derived fast-neutron susceptible mutant (***mla6*-m18982; red dashed line**), at 0, 16, 20, 24, 32 and 48 h after inoculation with *Bh* isolate 5874 ($AVR_{a1}$, $AVR_{a3}$, $AVR_{a6}$, $AVR_{a12}$; NCBI-GEO GSE101304) and analyzed as described in (Chapman *et al.*, 2021; Velásquez-Zapata *et al.*, 2022). Read counts were normalized using the taxon-specific method (Klingenberg and Meinicke, 2017). Genes differentially expressed at an adjusted *p* of <0.001 for barley and <0.003 for *Bh* were considered significant (Velásquez-Zapata *et al.*, 2024). Order of CSEP baits matches Figure 1 dendogram.

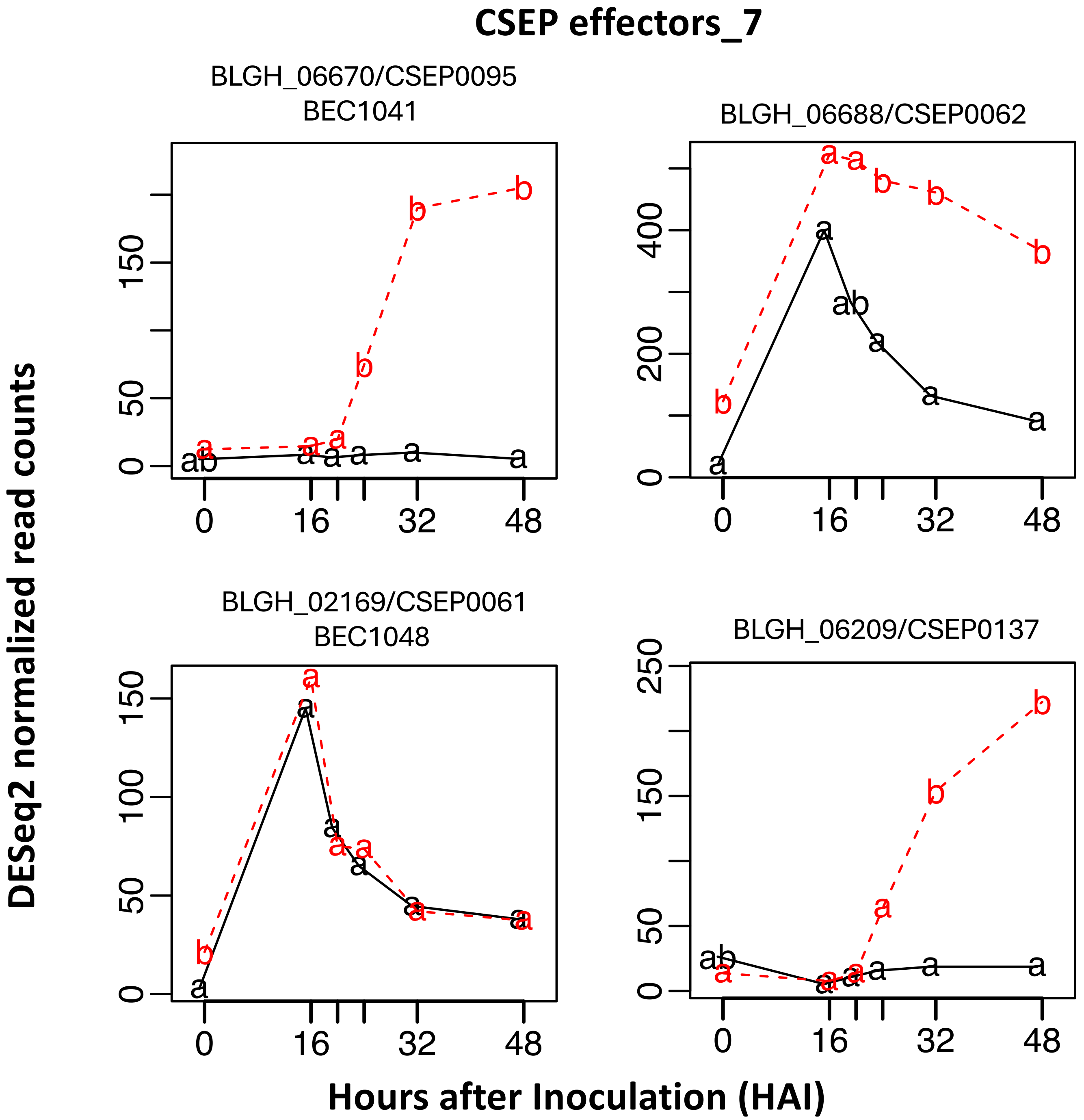


**Supporting Fig. S2-7.** Time-course expression of 48 *Blumeria hordei (Bh)* effector candidates, barley MLA, receptor-like kinase *Hv*SRF3_L1 (RLK5), and various host targets from the barley-*Bh* interactome. RNA-Sequencing data were extracted from an infection time course of barley CI 16151 (***Mla6*****; black solid line**) and the derived fast-neutron susceptible mutant (***mla6*****-m18982; red dashed line**), at 0, 16, 20, 24, 32 and 48 h after inoculation with *Bh* isolate 5874 ($AVR_{a1}$, $AVR_{a3}$, $AVR_{a6}$, $AVR_{a12}$; NCBI-GEO GSE101304) and analyzed as described in (Chapman *et al*., 2021; Velásquez-Zapata *et al*., 2022). Read counts were normalized using the taxon-specific method (Klingenberg and Meinicke, 2017). Genes differentially expressed at an adjusted *p* of <0.001 for barley and <0.003 for *Bh* were considered significant (Velásquez-Zapata *et al*., 2024). Order of CSEP baits matches Figure 1 dendogram.

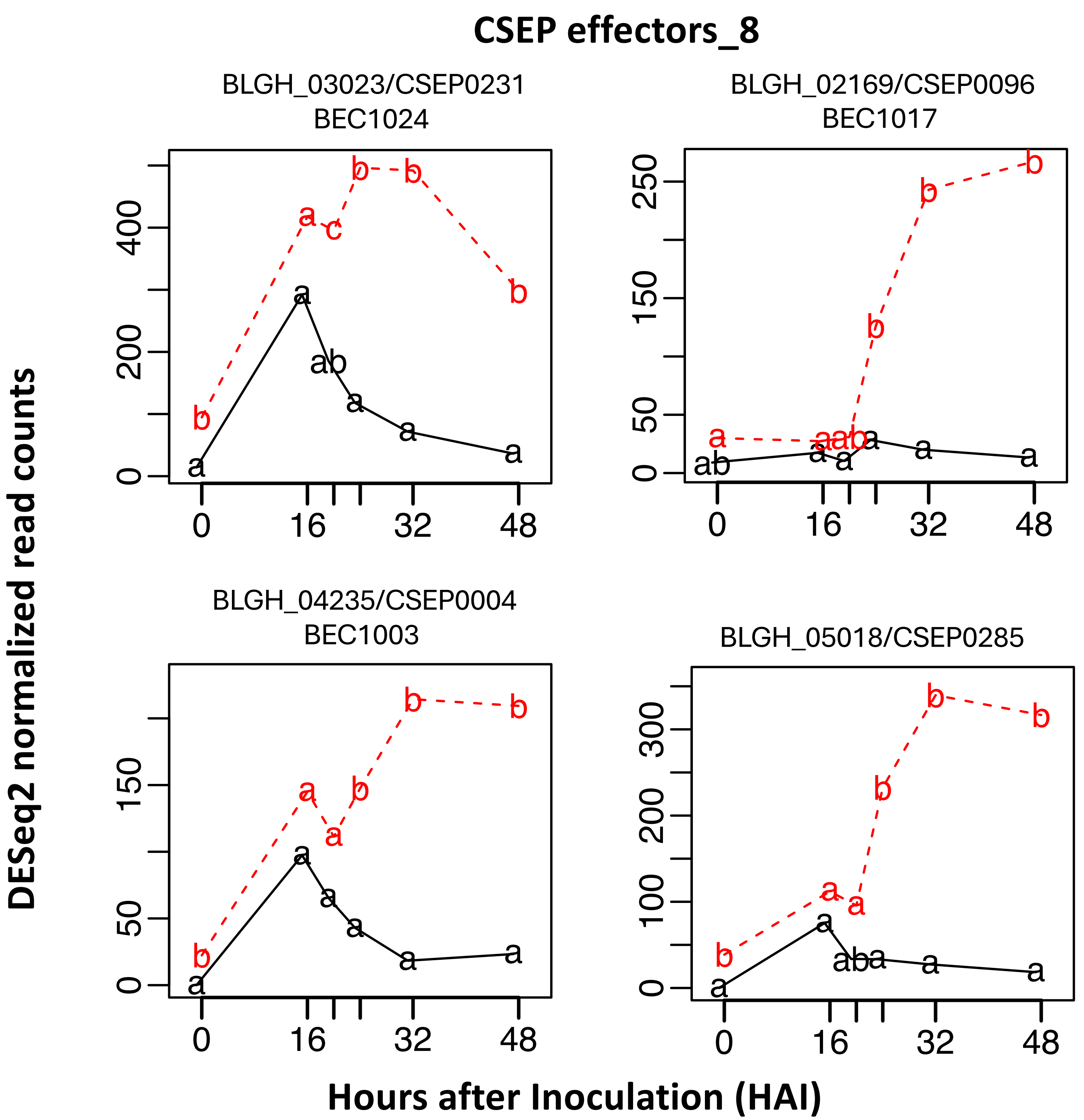


**Supporting Fig. S2-8.** Time-course expression of 48 *Blumeria hordei (Bh)* effector candidates, barley MLA, receptor-like kinase *Hv*SRF3_L1 (RLK5), and various host targets from the barley-*Bh* interactome. RNA-Sequencing data were extracted from an infection time course of barley CI 16151 (***Mla6*; black solid line**) and the derived fast-neutron susceptible mutant (***mla6*-m18982; red dashed line**), at 0, 16, 20, 24, 32 and 48 h after inoculation with *Bh* isolate 5874 ($AVR_{a1}$, $AVR_{a3}$, $AVR_{a6}$, $AVR_{a12}$; NCBI-GEO GSE101304) and analyzed as described in (Chapman *et al.*, 2021; Velásquez-Zapata *et al.*, 2022). Read counts were normalized using the taxon-specific method (Klingenberg and Meinicke, 2017). Genes differentially expressed at an adjusted *p* of <0.001 for barley and <0.003 for *Bh* were considered significant (Velásquez-Zapata *et al.*, 2024). Order of CSEP baits matches Figure 1 dendogram.

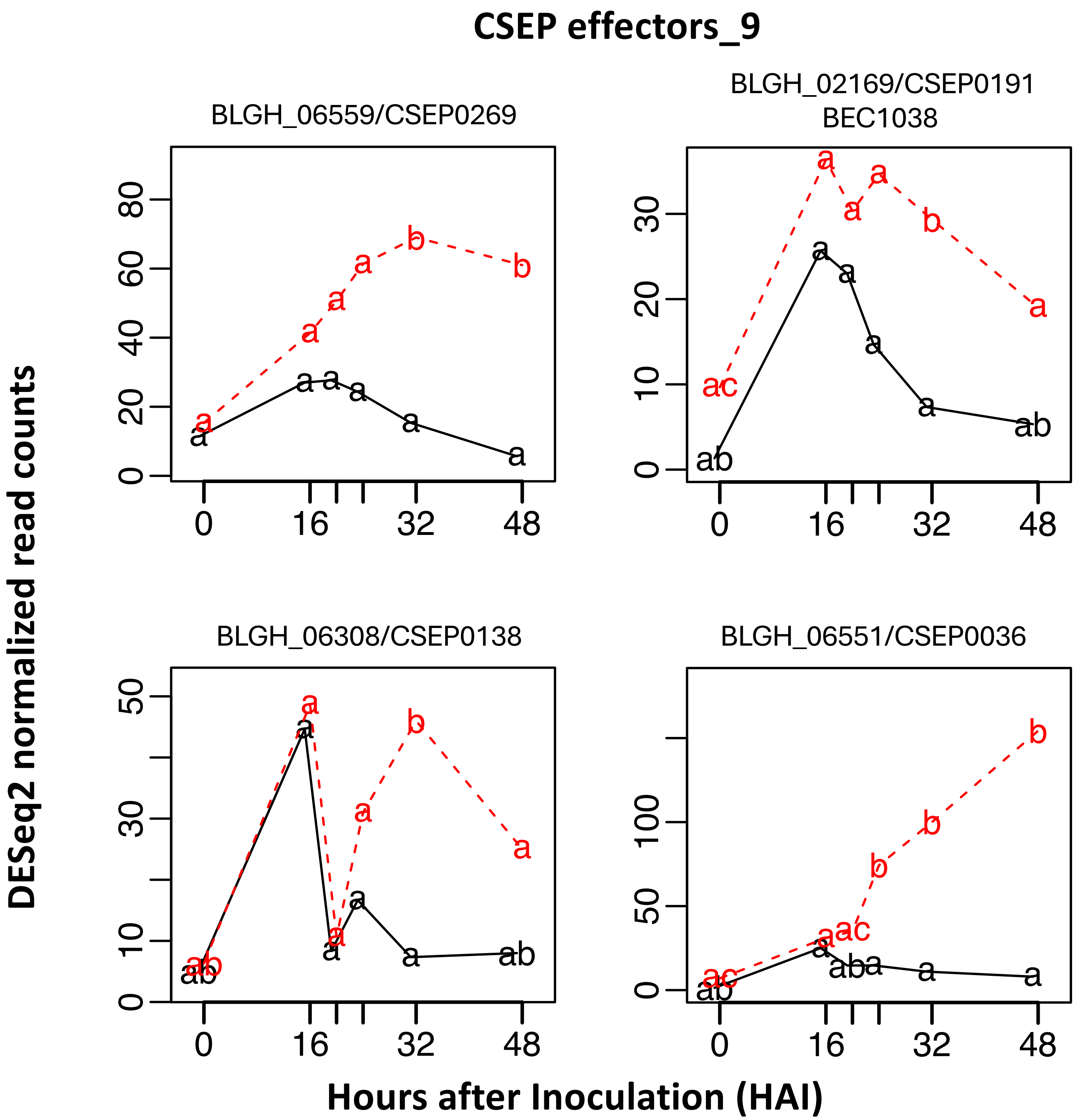


**Supporting Fig. S2-9.** Time-course expression of 48 *Blumeria hordei (Bh)* effector candidates, barley MLA, receptor-like kinase *Hv*SRF3_L1 (RLK5), and various host targets from the barley-*Bh* interactome. RNA-Sequencing data were extracted from an infection time course of barley CI 16151 (***Mla6*; black solid line**) and the derived fast-neutron susceptible mutant (***mla6*-m18982; red dashed line**), at 0, 16, 20, 24, 32 and 48 h after inoculation with *Bh* isolate 5874 ($AVR_{a1}$, $AVR_{a3}$, $AVR_{a6}$, $AVR_{a12}$; NCBI-GEO GSE101304) and analyzed as described in (Chapman *et al.*, 2021; Velásquez-Zapata *et al.*, 2022). Read counts were normalized using the taxon-specific method (Klingenberg and Meinicke, 2017). Genes differentially expressed at an adjusted *p* of <0.001 for barley and <0.003 for *Bh* were considered significant (Velásquez-Zapata *et al.*, 2024). Order of CSEP baits matches Figure 1 dendogram.

#### CSEP effectors_10

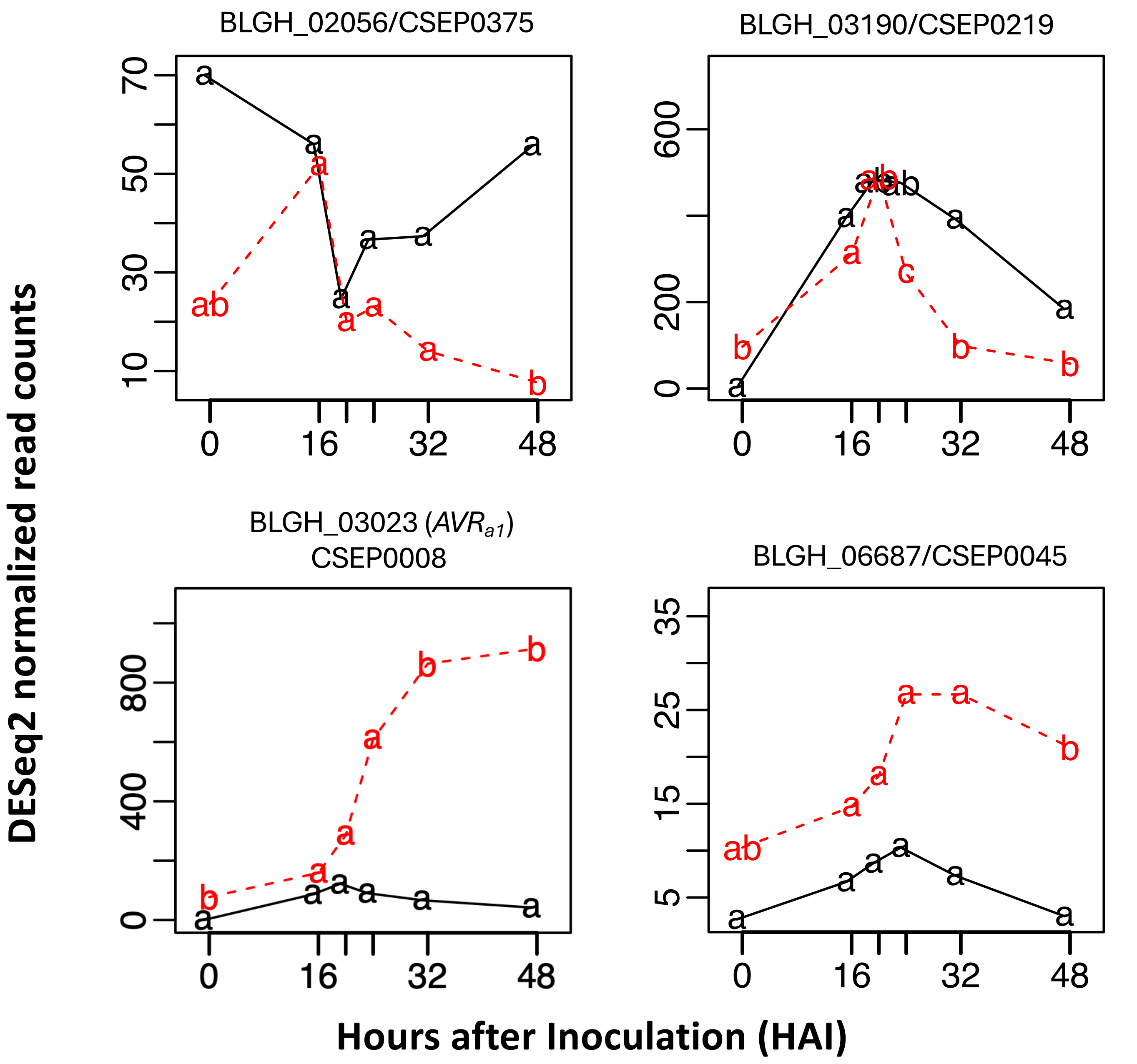


**Supporting Fig. S2-10.** Time-course expression of 48 *Blumeria hordei (Bh)* effector candidates, barley MLA, receptor-like kinase *Hv*SRF3_L1 (RLK5), and various host targets from the barley-*Bh* interactome. RNA-Sequencing data were extracted from an infection time course of barley CI 16151 (***Mla6*; black solid line**) and the derived fast-neutron susceptible mutant (***mla6*-m18982; red dashed line**), at 0, 16, 20, 24, 32 and 48 h after inoculation with *Bh* isolate 5874 ($AVR_{a1}$, $AVR_{a3}$, $AVR_{a6}$, $AVR_{a12}$; NCBI-GEO GSE101304) and analyzed as described in (Chapman *et al.*, 2021; Velásquez-Zapata *et al.*, 2022). Read counts were normalized using the taxon-specific method (Klingenberg and Meinicke, 2017). Genes differentially expressed at an adjusted *p* of <0.001 for barley and <0.003 for *Bh* were considered significant (Velásquez-Zapata *et al.*, 2024). Order of CSEP baits matches Figure 1 dendogram.

#### CSEP effectors_11

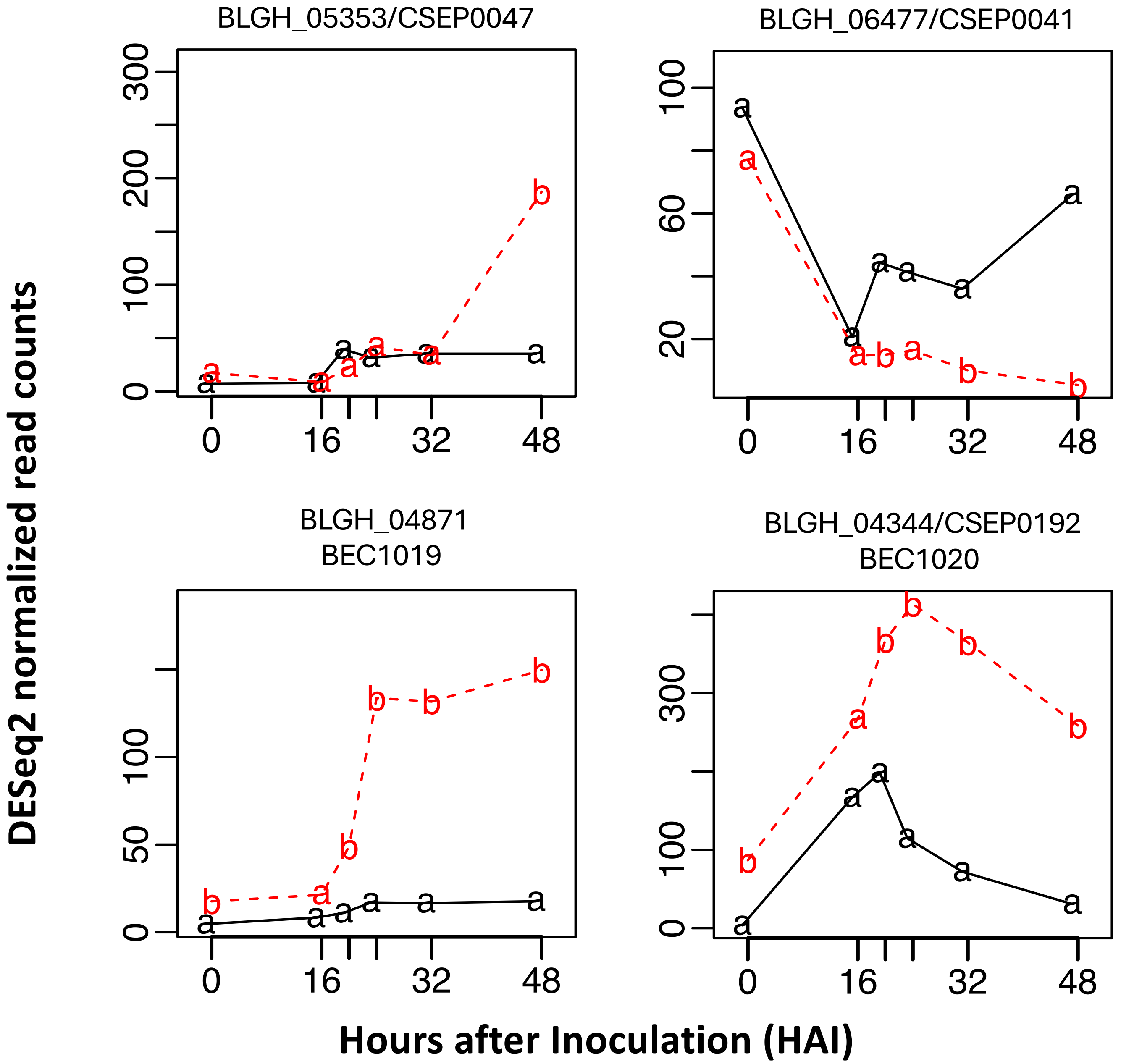


**Supporting Fig. S2-11.** Time-course expression of 48 *Blumeria hordei (Bh)* effector candidates, barley MLA, receptor-like kinase *Hv*SRF3_L1 (RLK5), and various host targets from the barley-*Bh* interactome. RNA-Sequencing data were extracted from an infection time course of barley CI 16151 (***Mla6*; black solid line**) and the derived fast-neutron susceptible mutant (***mla6*-m18982; red dashed line**), at 0, 16, 20, 24, 32 and 48 h after inoculation with *Bh* isolate 5874 ($AVR_{a1}$, $AVR_{a3}$, $AVR_{a6}$, $AVR_{a12}$; NCBI-GEO GSE101304) and analyzed as described in (Chapman *et al.*, 2021; Velásquez-Zapata *et al.*, 2022). Read counts were normalized using the taxon-specific method (Klingenberg and Meinicke, 2017). Genes differentially expressed at an adjusted *p* of <0.001 for barley and <0.003 for *Bh* were considered significant (Velásquez-Zapata *et al.*, 2024). Order of CSEP baits matches Figure 1 dendogram.

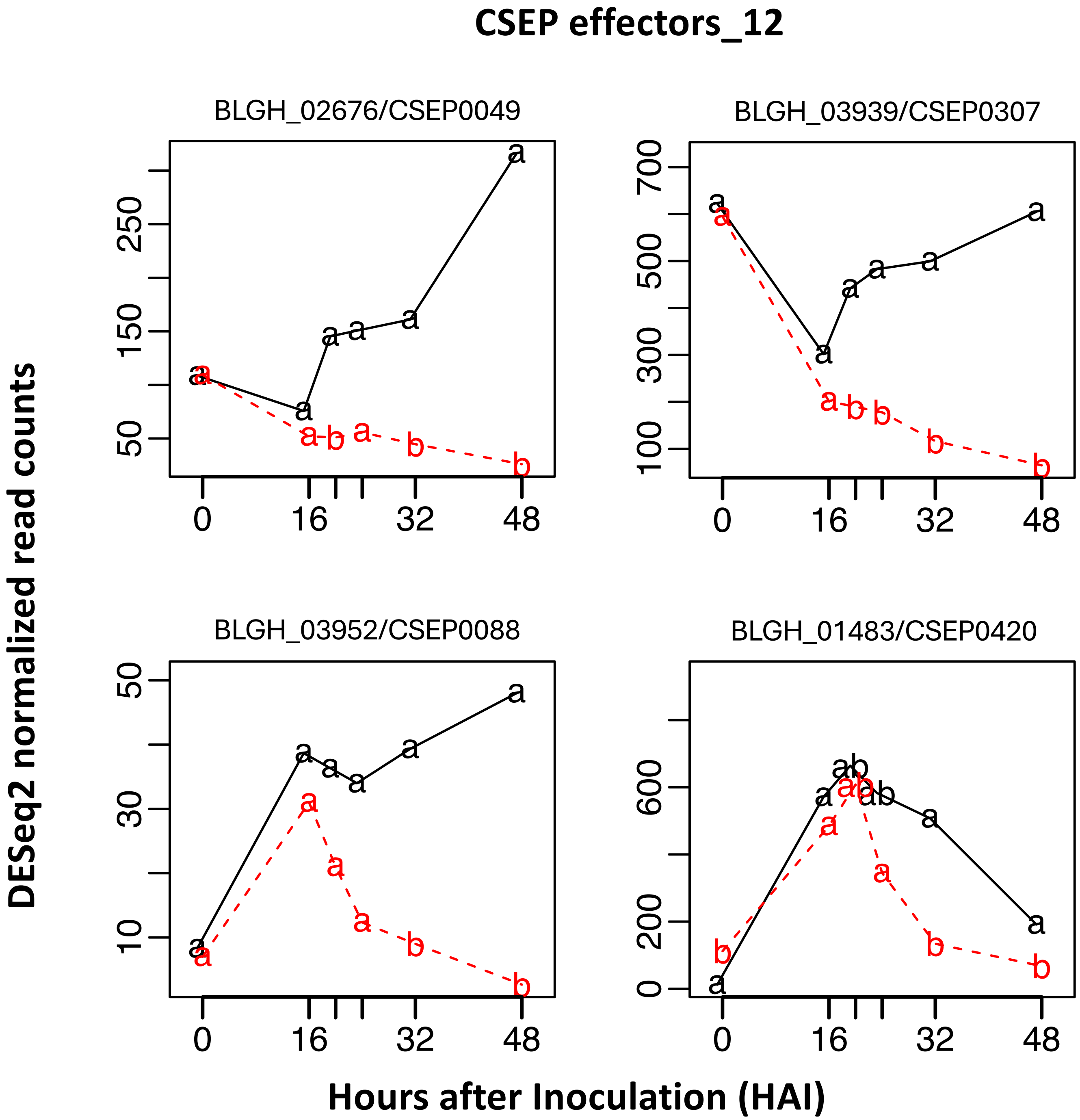


**Supporting Fig. S2-12.** Time-course expression of 48 *Blumeria hordei (Bh)* effector candidates, barley MLA, receptor-like kinase *Hv*SRF3_L1 (RLK5), and various host targets from the barley-*Bh* interactome. RNA-Sequencing data were extracted from an infection time course of barley CI 16151 (***Mla6*; black solid line**) and the derived fast-neutron susceptible mutant (***mla6*-m18982; red dashed line**), at 0, 16, 20, 24, 32 and 48 h after inoculation with *Bh* isolate 5874 ($AVR_{a1}$, $AVR_{a3}$, $AVR_{a6}$, $AVR_{a12}$; NCBI-GEO GSE101304) and analyzed as described in (Chapman *et al*., 2021; Velásquez-Zapata *et al*., 2022). Read counts were normalized using the taxon-specific method (Klingenberg and Meinicke, 2017). Genes differentially expressed at an adjusted *p* of <0.001 for barley and <0.003 for *Bh* were considered significant (Velásquez-Zapata *et al*., 2024). Order of CSEP baits matches Figure 1 dendogram.

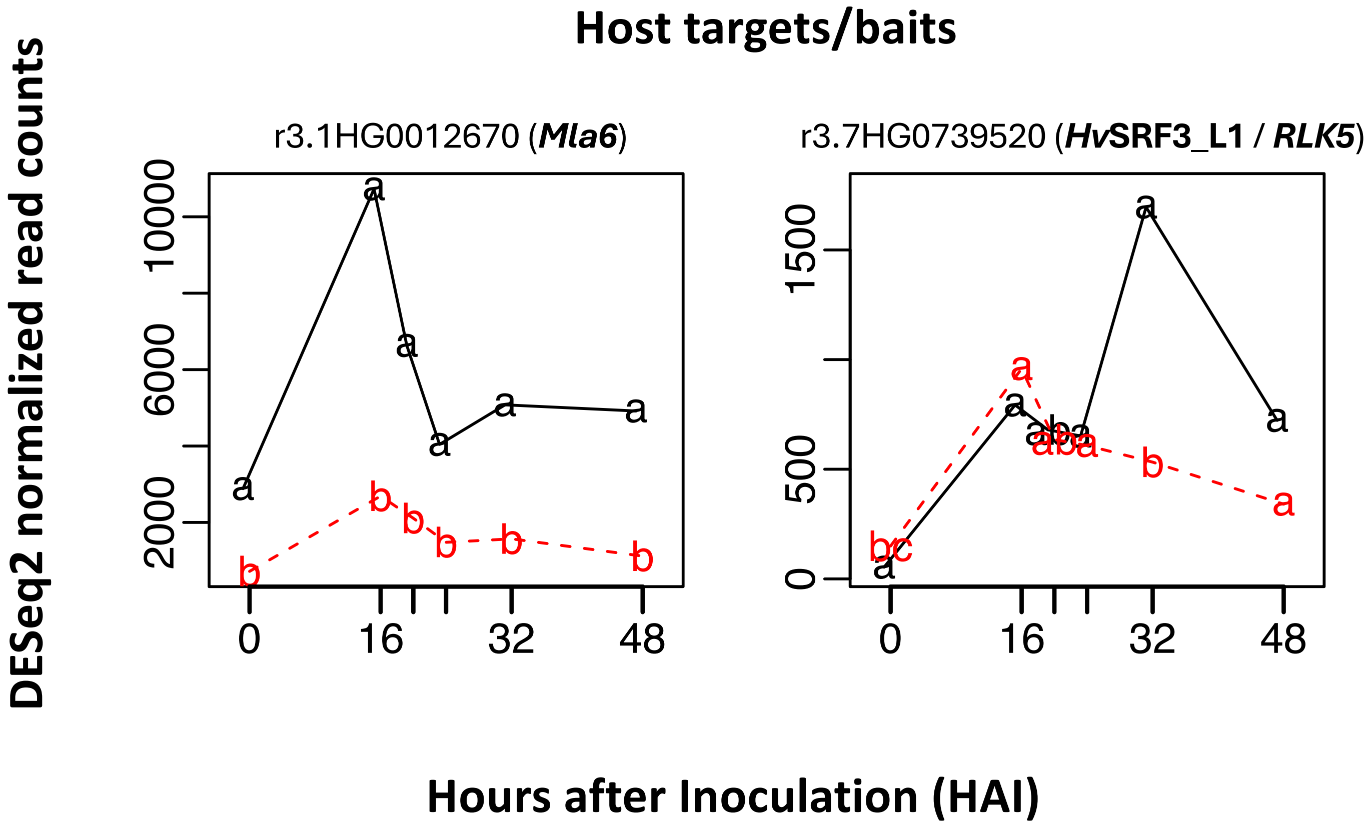


**Supporting Fig. S2-13.** Time-course expression of 48 *Blumeria hordei (Bh)* effector candidates, barley MLA, receptor-like kinase *Hv*SRF3_L1 (RLK5), and various host targets from the barley-*Bh* interactome. RNA-Sequencing data were extracted from an infection time course of barley CI 16151 (***Mla6*; black solid line**) and the derived fast-neutron susceptible mutant (***mla6*-m18982; red dashed line**), at 0, 16, 20, 24, 32 and 48 h after inoculation with *Bh* isolate 5874 ($AVR_{a1}$, $AVR_{a3}$, $AVR_{a6}$, $AVR_{a12}$; NCBI-GEO GSE101304) and analyzed as described in (Chapman *et al.*, 2021; Velásquez-Zapata *et al.*, 2022). Read counts were normalized using the taxon-specific method (Klingenberg and Meinicke, 2017). Genes differentially expressed at an adjusted *p* of <0.001 for barley and <0.003 for *Bh* were considered significant (Velásquez-Zapata *et al.*, 2024). Order of CSEP baits matches Figure 1 dendogram.

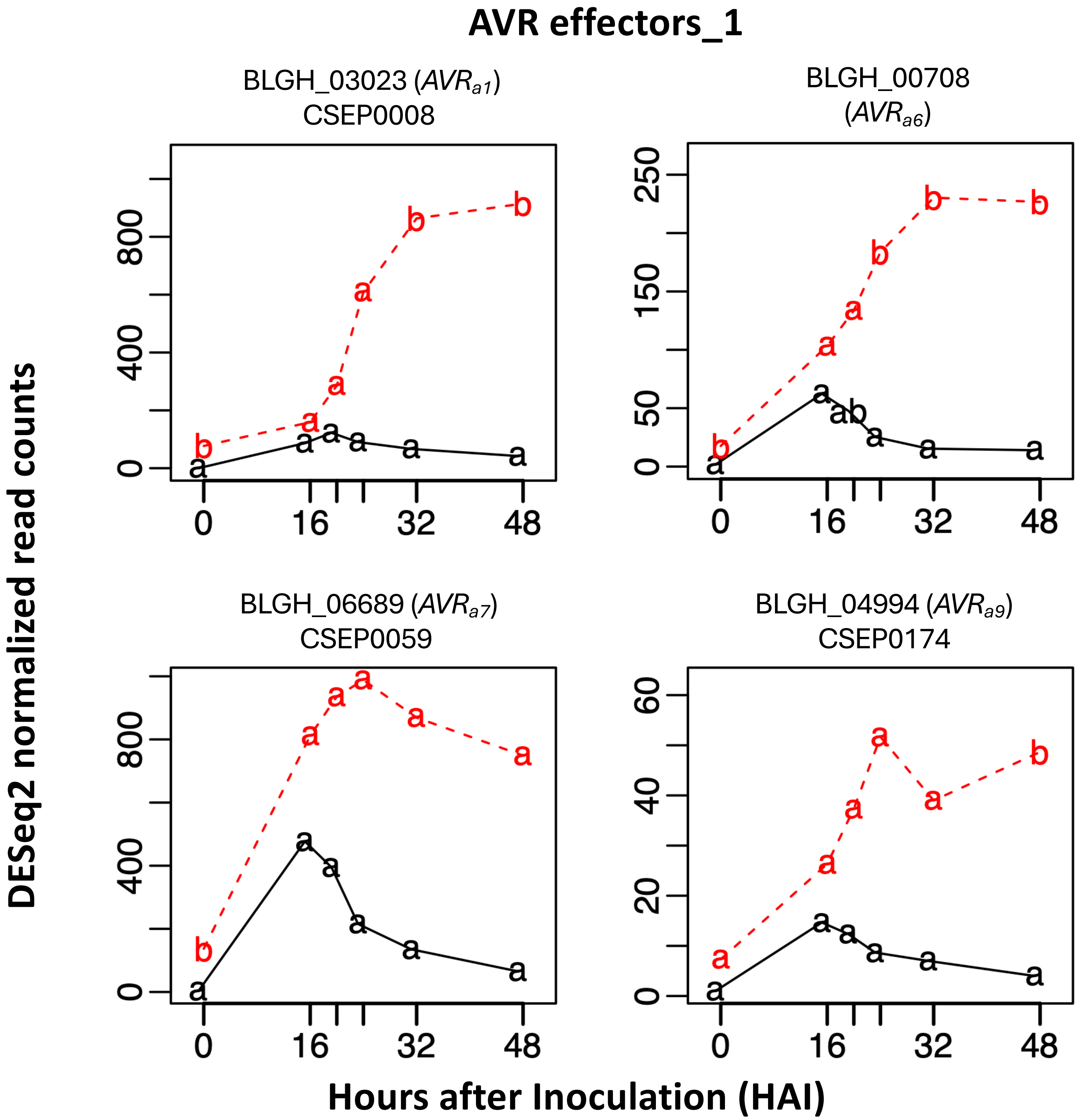


**Supporting Fig. S2-14.** Time-course expression of 48 *Blumeria hordei (Bh)* effector candidates, barley MLA, receptor-like kinase *Hv*SRF3_L1 (RLK5), and various host targets from the barley-*Bh* interactome. RNA-Sequencing data were extracted from an infection time course of barley CI 16151 (***Mla6*; black solid line**) and the derived fast-neutron susceptible mutant (***mla6*-m18982; red dashed line**), at 0, 16, 20, 24, 32 and 48 h after inoculation with *Bh* isolate 5874 ($AVR_{a1}$, $AVR_{a3}$, $AVR_{a6}$, $AVR_{a12}$; NCBI-GEO GSE101304) and analyzed as described in (Chapman *et al*., 2021; Velásquez-Zapata *et al*., 2022). Read counts were normalized using the taxon-specific method (Klingenberg and Meinicke, 2017). Genes differentially expressed at an adjusted *p* of <0.001 for barley and <0.003 for *Bh* were considered significant (Velásquez-Zapata *et al*., 2024). Order of CSEP baits matches Figure 1 dendogram.

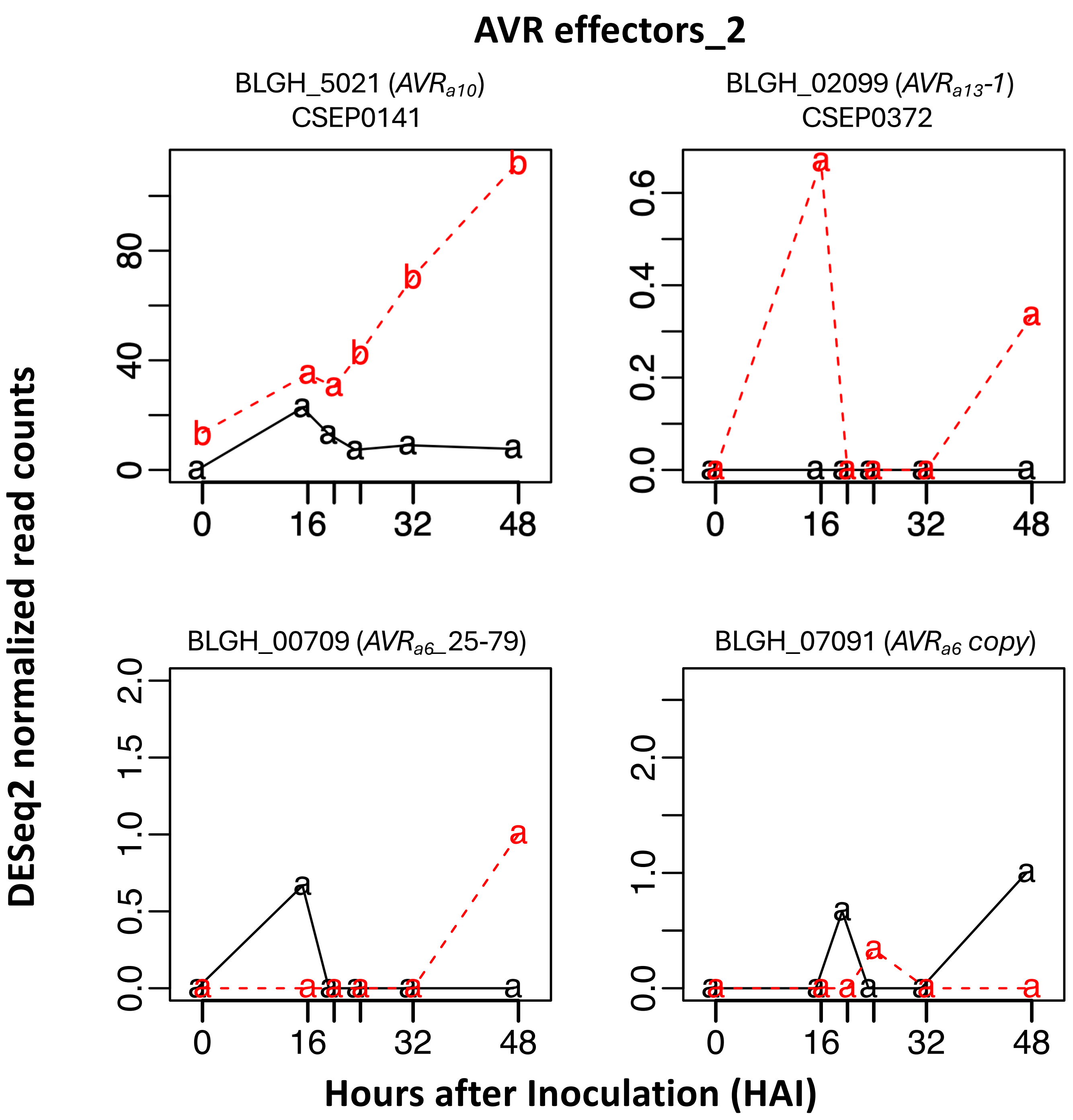


**Supporting Fig. S2-15.** Time-course expression of 48 *Blumeria hordei (Bh)* effector candidates, barley MLA, receptor-like kinase *Hv*SRF3_L1 (RLK5), and various host targets from the barley-*Bh* interactome. RNA-Sequencing data were extracted from an infection time course of barley CI 16151 (***Mla6*; black solid line**) and the derived fast-neutron susceptible mutant (***mla6*-m18982; red dashed line**), at 0, 16, 20, 24, 32 and 48 h after inoculation with *Bh* isolate 5874 (*AVR$_{a1}$, AVR$_{a3}$, AVR$_{a6}$, AVR$_{a12}$*; NCBI-GEO GSE101304) and analyzed as described in (Chapman *et al.*, 2021; Velásquez-Zapata *et al.*, 2022). Read counts were normalized using the taxon-specific method (Klingenberg and Meinicke, 2017). Genes differentially expressed at an adjusted *p* of <0.001 for barley and <0.003 for *Bh* were considered significant (Velásquez-Zapata *et al.*, 2024). Order of CSEP baits matches Figure 1 dendogram.

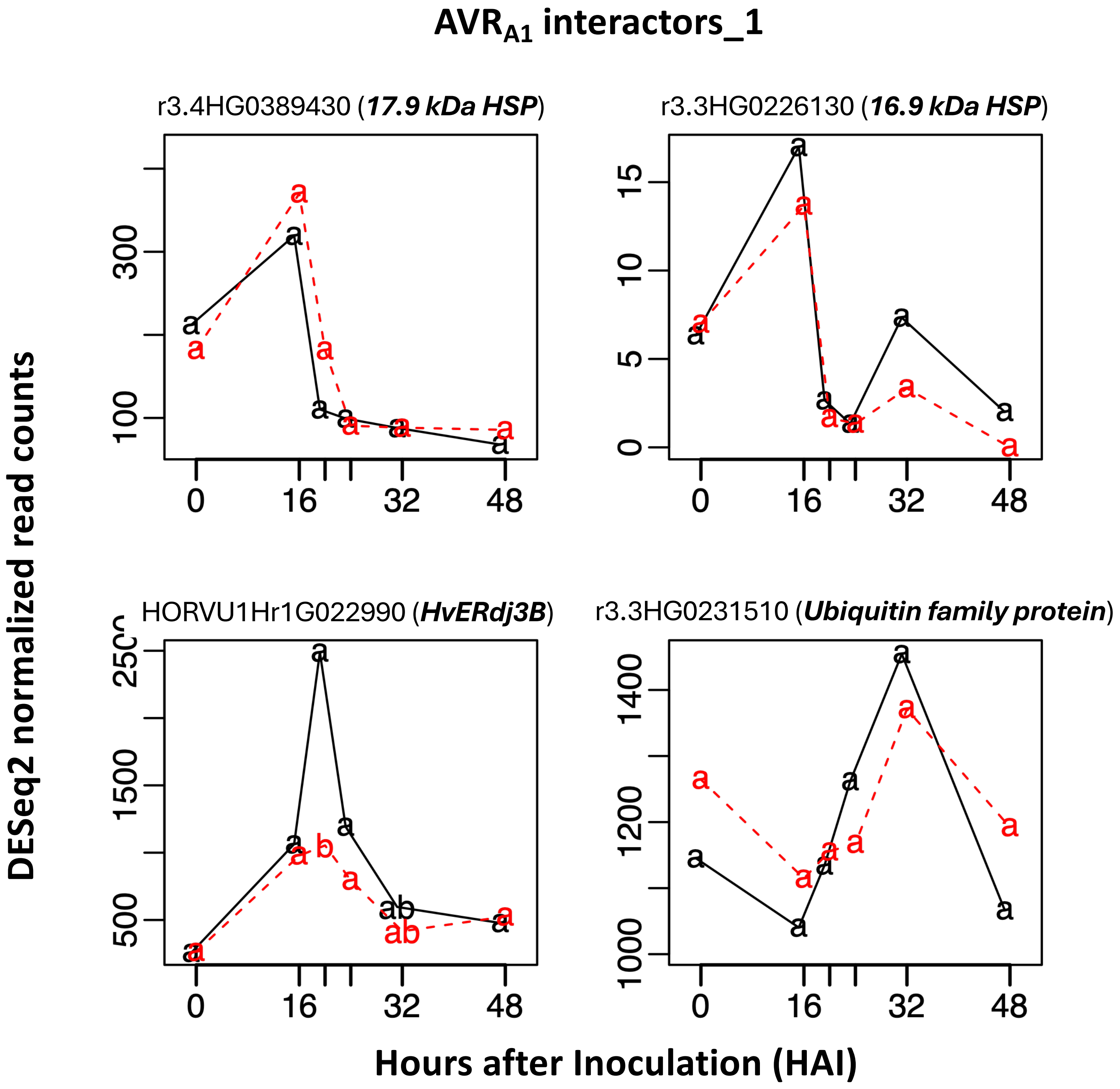


**Supporting Fig. S2-16.** Time-course expression of 48 *Blumeria hordei (Bh)* effector candidates, barley MLA, receptor-like kinase *Hv*SRF3_L1 (RLK5), and various host targets from the barley-*Bh* interactome. RNA-Sequencing data were extracted from an infection time course of barley CI 16151 (***Mla6*; black solid line**) and the derived fast-neutron susceptible mutant (***mla6*-m18982; red dashed line**), at 0, 16, 20, 24, 32 and 48 h after inoculation with *Bh* isolate 5874 ($AVR_{a1}$, $AVR_{a3}$, $AVR_{a6}$, $AVR_{a12}$; NCBI-GEO GSE101304) and analyzed as described in (Chapman *et al.*, 2021; Velásquez-Zapata *et al.*, 2022). Read counts were normalized using the taxon-specific method (Klingenberg and Meinicke, 2017). Genes differentially expressed at an adjusted *p* of <0.001 for barley and <0.003 for *Bh* were considered significant (Velásquez-Zapata *et al.*, 2024). Order of CSEP baits matches Figure 1 dendogram.

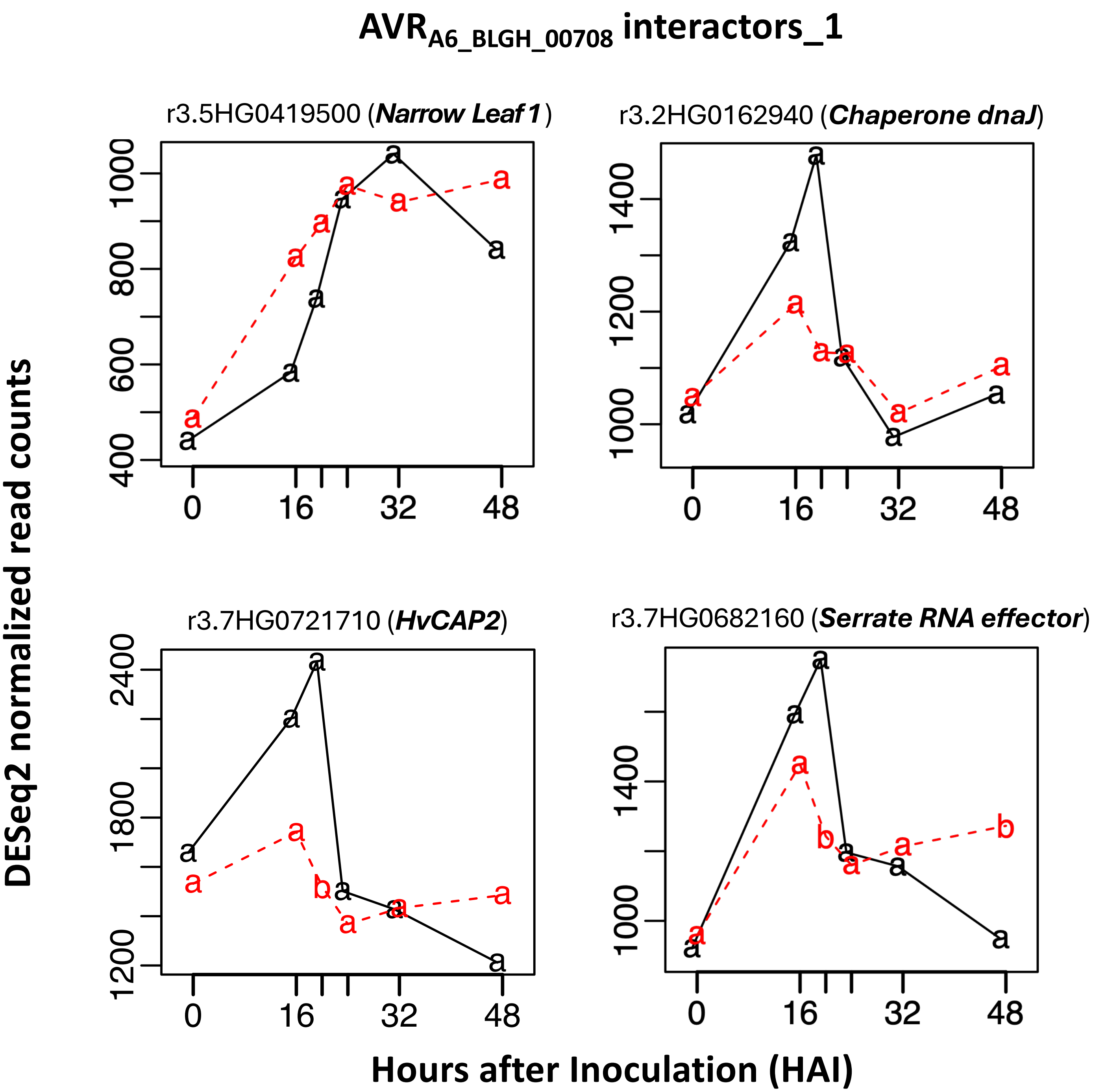


**Supporting Fig. S2-17.** Time-course expression of 48 *Blumeria hordei (Bh)* effector candidates, barley MLA, receptor-like kinase *Hv*SRF3_L1 (RLK5), and various host targets from the barley-*Bh* interactome. RNA-Sequencing data were extracted from an infection time course of barley CI 16151 (***Mla6*; black solid line**) and the derived fast-neutron susceptible mutant (***mla6*-m18982; red dashed line**), at 0, 16, 20, 24, 32 and 48 h after inoculation with *Bh* isolate 5874 ($AVR_{a1}$, $AVR_{a3}$, $AVR_{a6}$, $AVR_{a12}$; NCBI-GEO GSE101304) and analyzed as described in (Chapman *et al.*, 2021; Velásquez-Zapata *et al.*, 2022). Read counts were normalized using the taxon-specific method (Klingenberg and Meinicke, 2017). Genes differentially expressed at an adjusted *p* of <0.001 for barley and <0.003 for *Bh* were considered significant (Velásquez-Zapata *et al.*, 2024). Order of CSEP baits matches Figure 1 dendogram.

#### $AVR_{A6\_BLGH\_00708}$ interactors_2

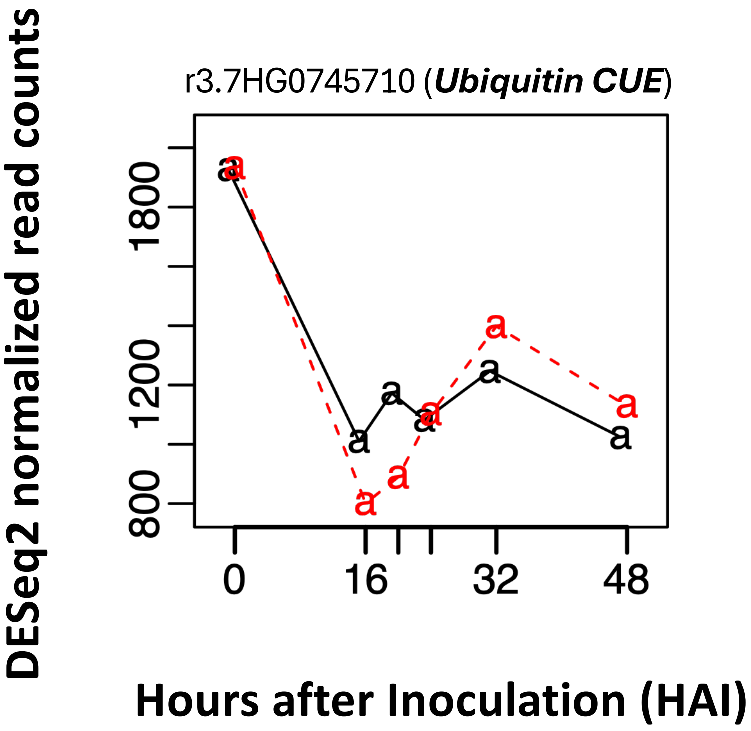


**Supporting Fig. S2-18.** Time-course expression of 48 *Blumeria hordei (Bh)* effector candidates, barley MLA, receptor-like kinase *Hv*SRF3_L1 (RLK5), and various host targets from the barley-*Bh* interactome. RNA-Sequencing data were extracted from an infection time course of barley CI 16151 (***Mla6*; black solid line**) and the derived fast-neutron susceptible mutant (***mla6*-m18982; red dashed line**), at 0, 16, 20, 24, 32 and 48 h after inoculation with *Bh* isolate 5874 ($AVR_{a1}$, $AVR_{a3}$, $AVR_{a6}$, $AVR_{a12}$; NCBI-GEO GSE101304) and analyzed as described in (Chapman *et al.*, 2021; Velásquez-Zapata *et al.*, 2022). Read counts were normalized using the taxon-specific method (Klingenberg and Meinicke, 2017). Genes differentially expressed at an adjusted *p* of <0.001 for barley and <0.003 for *Bh* were considered significant (Velásquez-Zapata *et al.*, 2024). Order of CSEP baits matches Figure 1 dendogram.

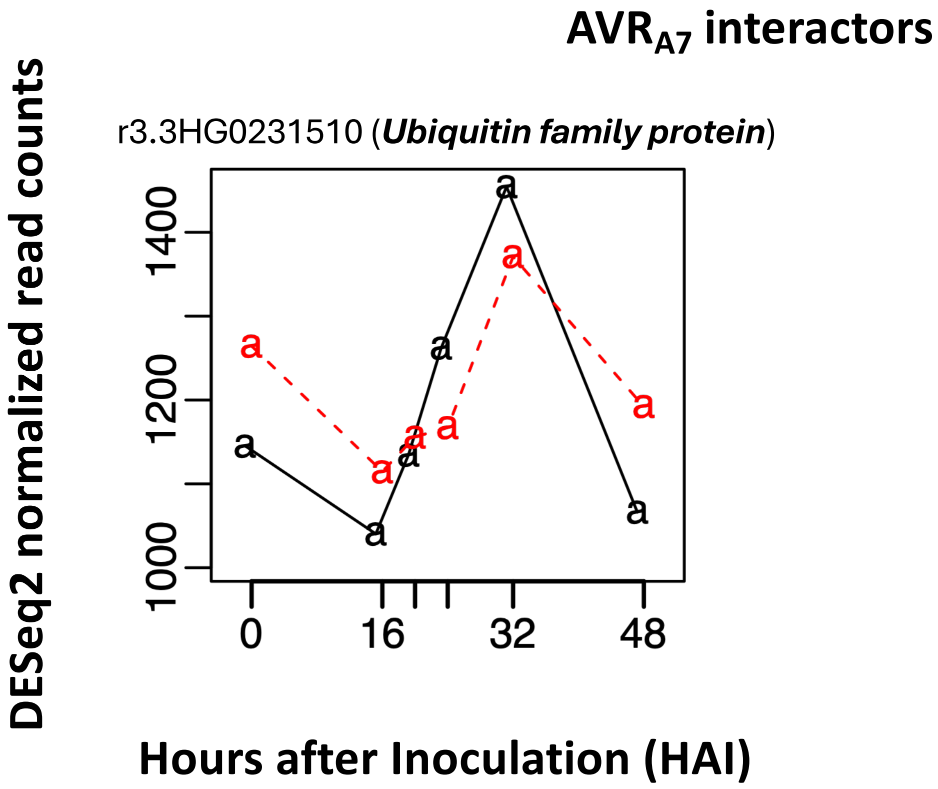


**Supporting Fig. S2-19.** Time-course expression of 48 *Blumeria hordei (Bh)* effector candidates, barley MLA, receptor-like kinase *Hv*SRF3_L1 (RLK5), and various host targets from the barley-*Bh* interactome. RNA-Sequencing data were extracted from an infection time course of barley CI 16151 (***Mla6*; black solid line**) and the derived fast-neutron susceptible mutant (***mla6*-m18982; red dashed line**), at 0, 16, 20, 24, 32 and 48 h after inoculation with *Bh* isolate 5874 (*AVR$_{a1}$, AVR$_{a3}$, AVR$_{a6}$, AVR$_{a12}$*; NCBI-GEO GSE101304) and analyzed as described in (Chapman *et al*., 2021; Velásquez-Zapata *et al*., 2022). Read counts were normalized using the taxon-specific method (Klingenberg and Meinicke, 2017). Genes differentially expressed at an adjusted *p* of <0.001 for barley and <0.003 for *Bh* were considered significant (Velásquez-Zapata *et al*., 2024). Order of CSEP baits matches Figure 1 dendogram.

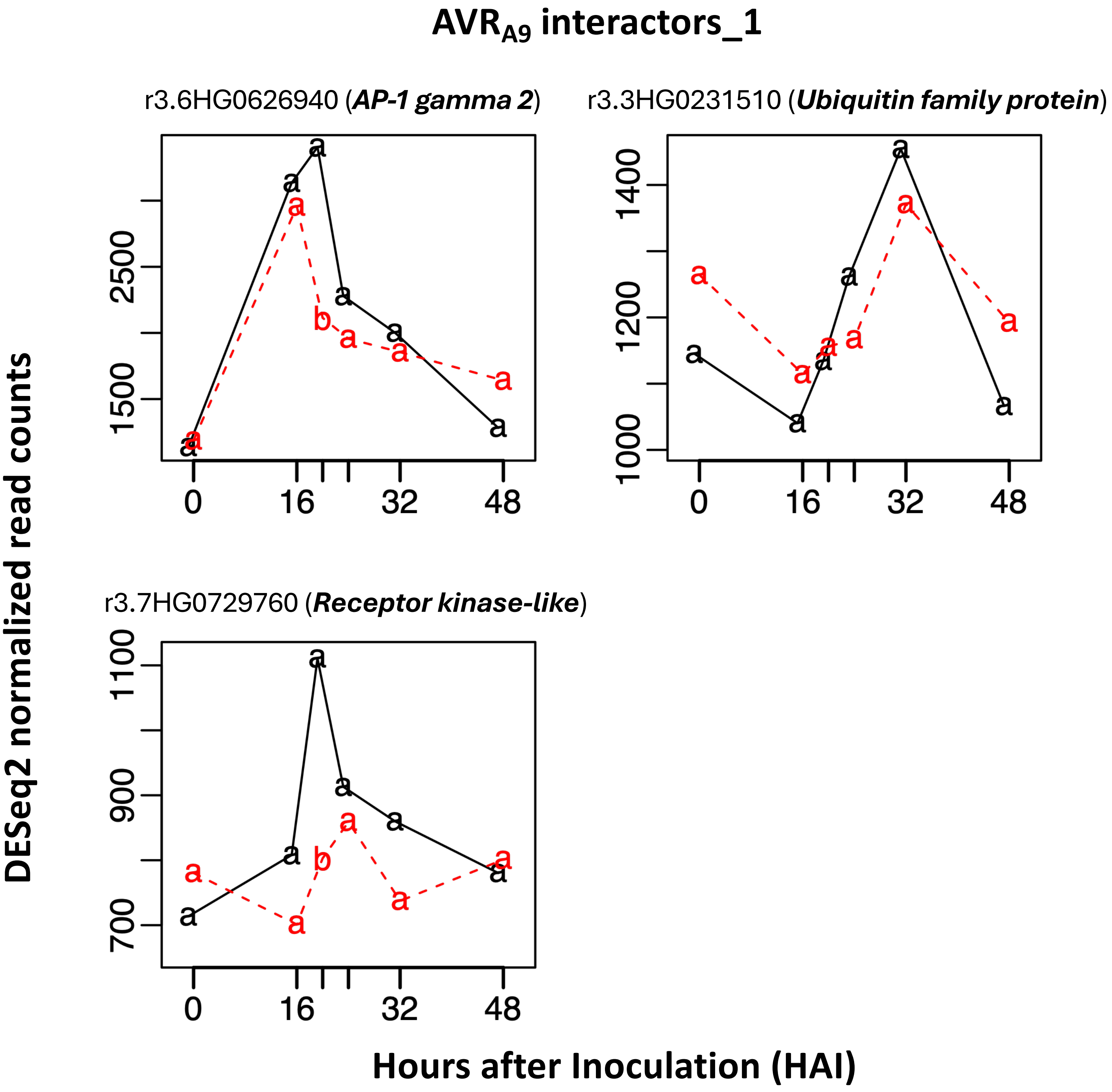


**Supporting Fig. S2-20.** Time-course expression of 48 *Blumeria hordei (Bh)* effector candidates, barley MLA, receptor-like kinase *Hv*SRF3_L1 (RLK5), and various host targets from the barley-*Bh* interactome. RNA-Sequencing data were extracted from an infection time course of barley CI 16151 (***Mla6*; black solid line**) and the derived fast-neutron susceptible mutant (***mla6*-m18982; red dashed line**), at 0, 16, 20, 24, 32 and 48 h after inoculation with *Bh* isolate 5874 ($AVR_{a1}$, $AVR_{a3}$, $AVR_{a6}$, $AVR_{a12}$; NCBI-GEO GSE101304) and analyzed as described in (Chapman *et al.*, 2021; Velásquez-Zapata *et al.*, 2022). Read counts were normalized using the taxon-specific method (Klingenberg and Meinicke, 2017). Genes differentially expressed at an adjusted *p* of <0.001 for barley and <0.003 for *Bh* were considered significant (Velásquez-Zapata *et al.*, 2024). Order of CSEP baits matches Figure 1 dendogram.

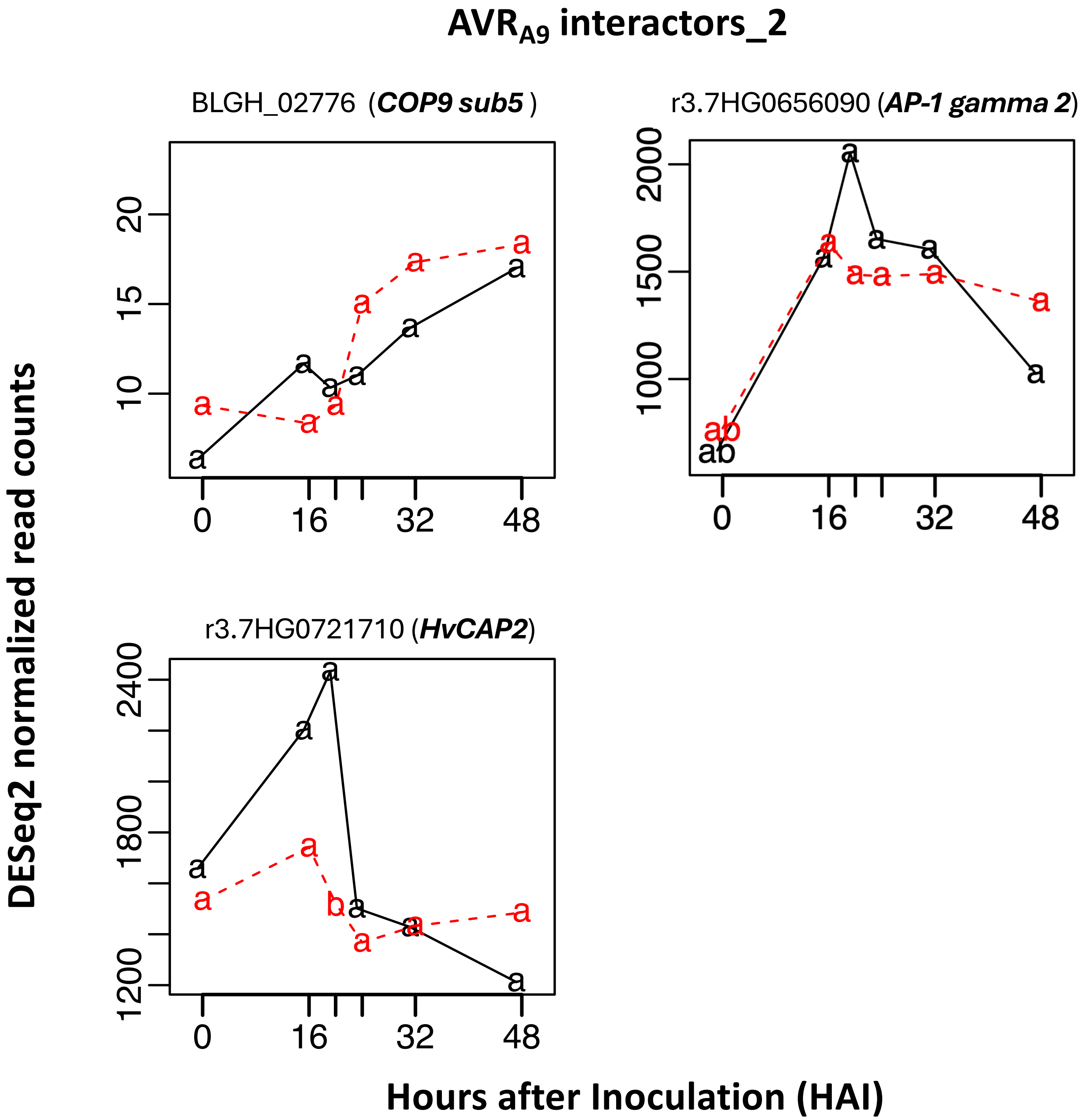


**Supporting Fig. S2-21.** Time-course expression of 48 *Blumeria hordei (Bh)* effector candidates, barley MLA, receptor-like kinase *Hv*SRF3_L1 (RLK5), and various host targets from the barley-*Bh* interactome. RNA-Sequencing data were extracted from an infection time course of barley CI 16151 (***Mla6*; black solid line**) and the derived fast-neutron susceptible mutant (***mla6*-m18982; red dashed line**), at 0, 16, 20, 24, 32 and 48 h after inoculation with *Bh* isolate 5874 (*AVR$_{a1}$, AVR$_{a3}$, AVR$_{a6}$, AVR$_{a12}$*; NCBI-GEO GSE101304) and analyzed as described in (Chapman *et al.*, 2021; Velásquez-Zapata *et al.*, 2022). Read counts were normalized using the taxon-specific method (Klingenberg and Meinicke, 2017). Genes differentially expressed at an adjusted *p* of <0.001 for barley and <0.003 for *Bh* were considered significant (Velásquez-Zapata *et al.*, 2024). Order of CSEP baits matches Figure 1 dendogram.

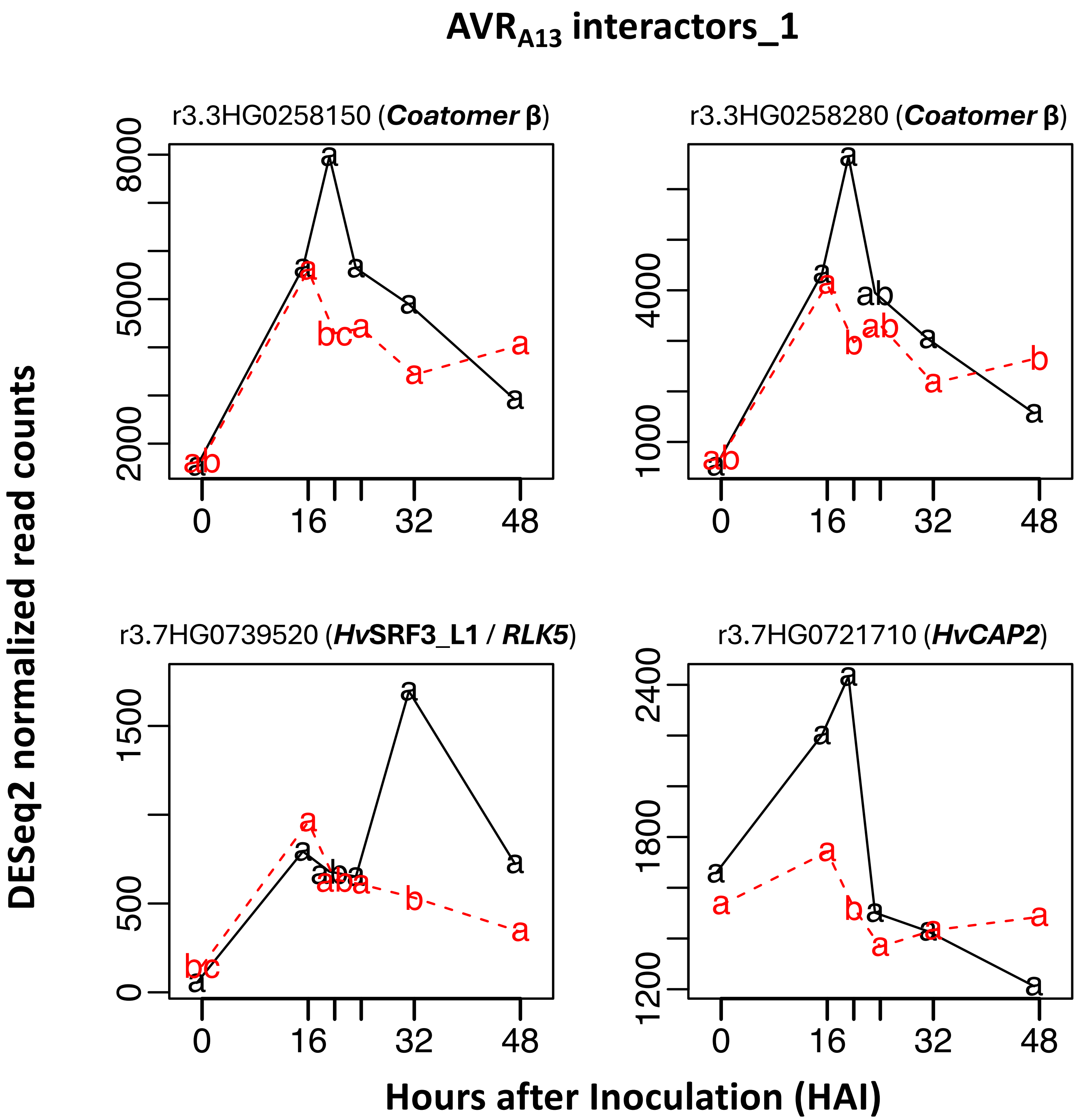


**Supporting Fig. S2-22.** Time-course expression of 48 *Blumeria hordei (Bh)* effector candidates, barley MLA, receptor-like kinase *Hv*SRF3_L1 (RLK5), and various host targets from the barley-*Bh* interactome. RNA-Sequencing data were extracted from an infection time course of barley CI 16151 (***Mla6*; black solid line**) and the derived fast-neutron susceptible mutant (***mla6*-m18982; red dashed line**), at 0, 16, 20, 24, 32 and 48 h after inoculation with *Bh* isolate 5874 (*AVR$_{a1}$, AVR$_{a3}$, AVR$_{a6}$, AVR$_{a12}$*; NCBI-GEO GSE101304) and analyzed as described in (Chapman *et al.*, 2021; Velásquez-Zapata *et al.*, 2022). Read counts were normalized using the taxon-specific method (Klingenberg and Meinicke, 2017). Genes differentially expressed at an adjusted *p* of <0.001 for barley and <0.003 for *Bh* were considered significant (Velásquez-Zapata *et al.*, 2024). Order of CSEP baits matches Figure 1 dendogram.

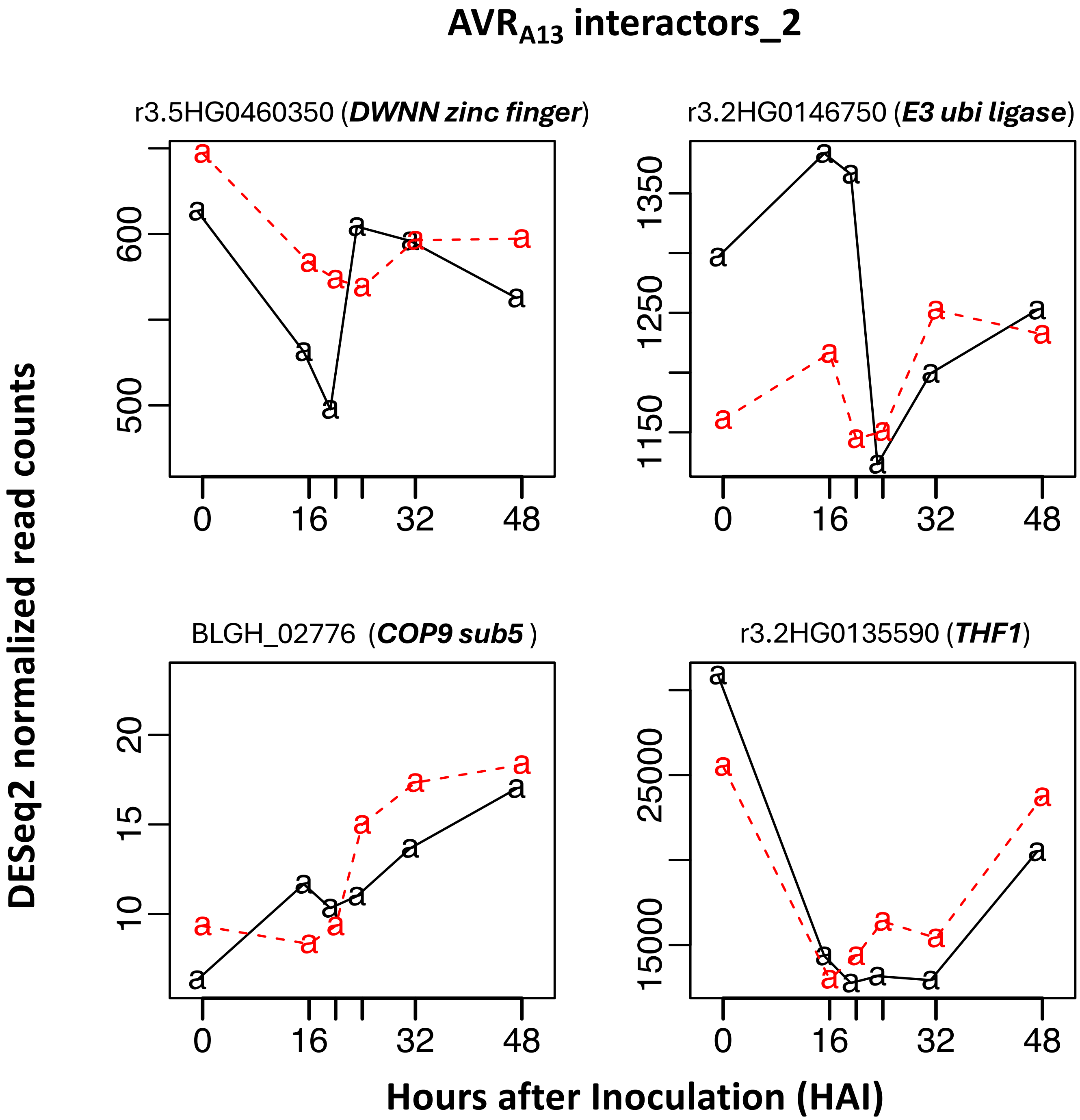


**Supporting Fig. S2-23.** Time-course expression of 48 *Blumeria hordei (Bh)* effector candidates, barley MLA, receptor-like kinase *Hv*SRF3_L1 (RLK5), and various host targets from the barley-*Bh* interactome. RNA-Sequencing data were extracted from an infection time course of barley CI 16151 (***Mla6*****; black solid line**) and the derived fast-neutron susceptible mutant (***mla6*****-m18982; red dashed line**), at 0, 16, 20, 24, 32 and 48 h after inoculation with *Bh* isolate 5874 ($AVR_{a1}$, $AVR_{a3}$, $AVR_{a6}$, $AVR_{a12}$; NCBI-GEO GSE101304) and analyzed as described in (Chapman *et al.*, 2021; Velásquez-Zapata *et al.*, 2022). Read counts were normalized using the taxon-specific method (Klingenberg and Meinicke, 2017). Genes differentially expressed at an adjusted *p* of <0.001 for barley and <0.003 for *Bh* were considered significant (Velásquez-Zapata *et al.*, 2024). Order of CSEP baits matches Figure 1 dendogram.

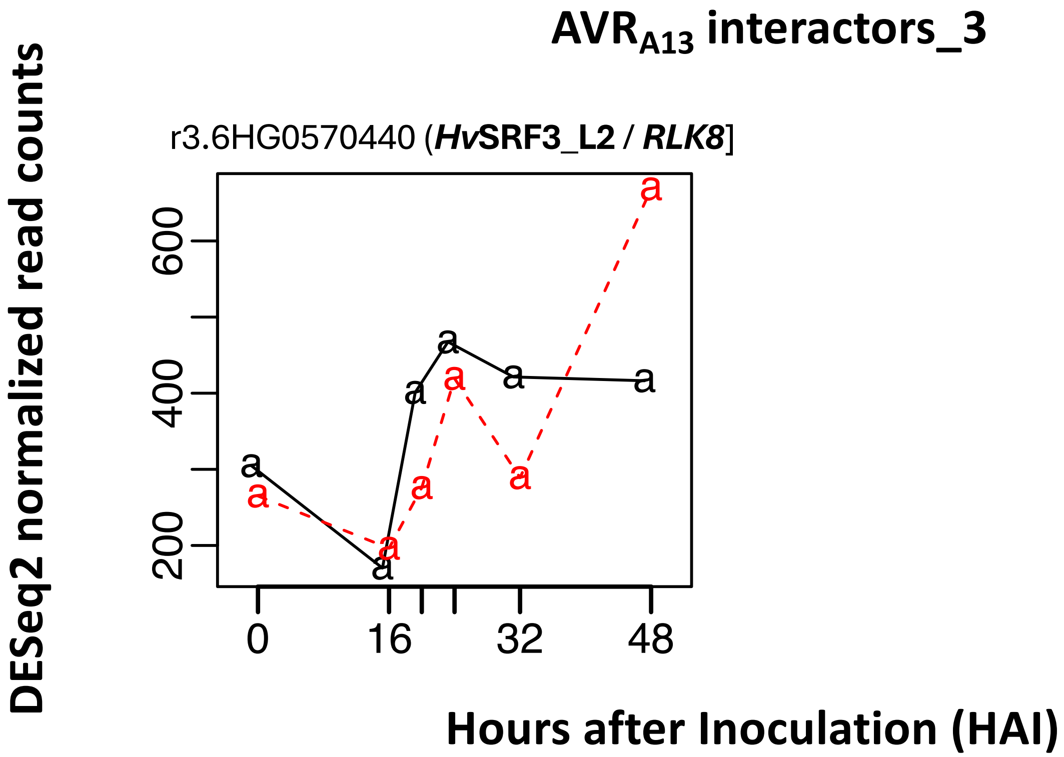


**Supporting Fig. S2-24.** Time-course expression of 48 *Blumeria hordei (Bh)* effector candidates, barley MLA, receptor-like kinase *Hv*SRF3_L1 (RLK5), and various host targets from the barley-*Bh* interactome. RNA-Sequencing data were extracted from an infection time course of barley CI 16151 (***Mla6*; black solid line**) and the derived fast-neutron susceptible mutant (***mla6*-m18982; red dashed line**), at 0, 16, 20, 24, 32 and 48 h after inoculation with *Bh* isolate 5874 ($AVR_{a1}$, $AVR_{a3}$, $AVR_{a6}$, $AVR_{a12}$; NCBI-GEO GSE101304) and analyzed as described in (Chapman *et al.*, 2021; Velásquez-Zapata *et al.*, 2022). Read counts were normalized using the taxon-specific method (Klingenberg and Meinicke, 2017). Genes differentially expressed at an adjusted *p* of <0.001 for barley and <0.003 for *Bh* were considered significant (Velásquez-Zapata *et al.*, 2024). Order of CSEP baits matches Figure 1 dendogram.

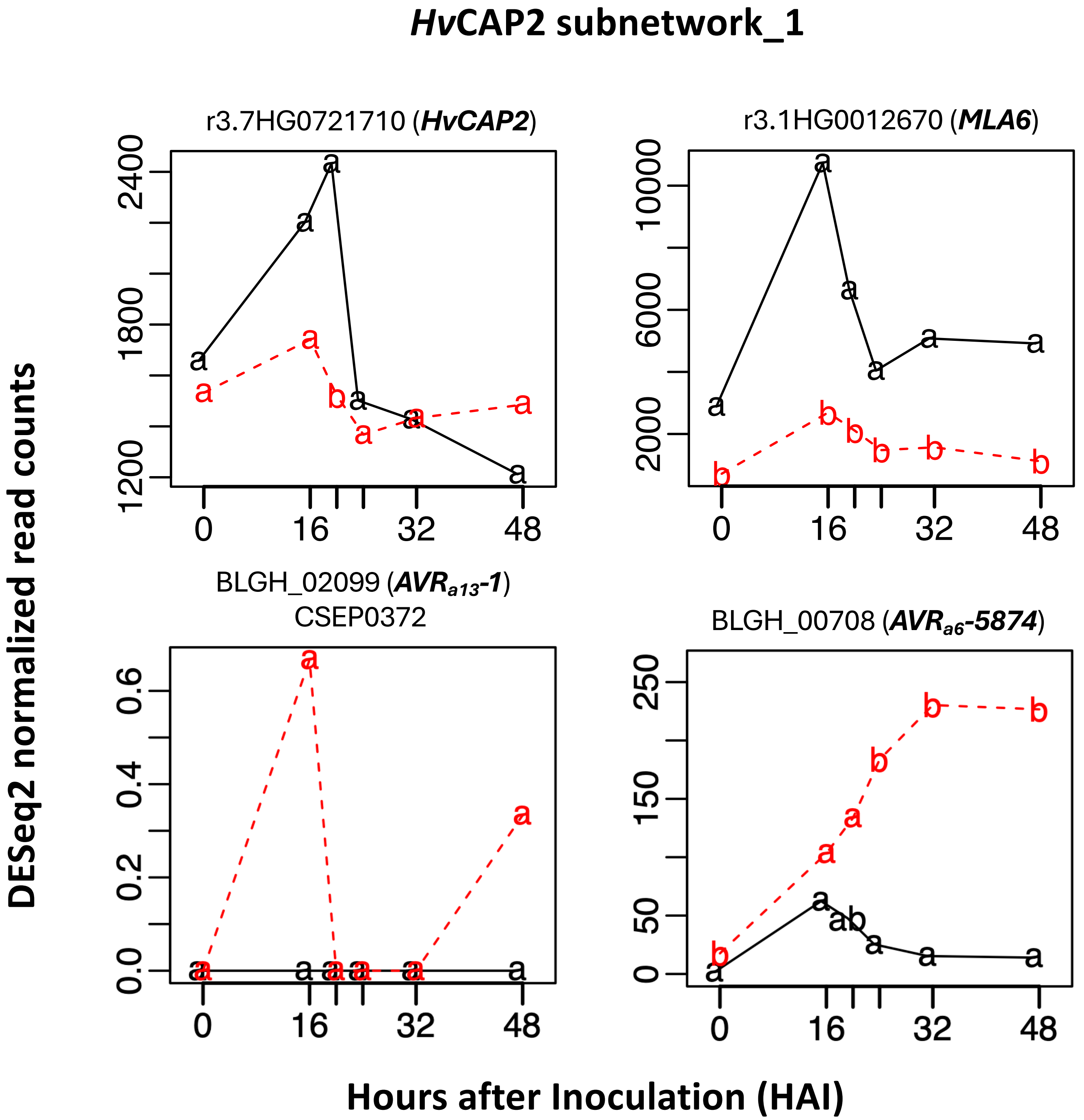


**Supporting Fig. S2-25.** Time-course expression of 48 *Blumeria hordei (Bh)* effector candidates, barley MLA, receptor-like kinase *Hv*SRF3_L1 (RLK5), and various host targets from the barley-*Bh* interactome. RNA-Sequencing data were extracted from an infection time course of barley CI 16151 (***Mla6*; black solid line**) and the derived fast-neutron susceptible mutant (***mla6*-m18982; red dashed line**), at 0, 16, 20, 24, 32 and 48 h after inoculation with *Bh* isolate 5874 ($AVR_{a1}$, $AVR_{a3}$, $AVR_{a6}$, $AVR_{a12}$; NCBI-GEO GSE101304) and analyzed as described in (Chapman *et al.*, 2021; Velásquez-Zapata *et al.*, 2022). Read counts were normalized using the taxon-specific method (Klingenberg and Meinicke, 2017). Genes differentially expressed at an adjusted *p* of <0.001 for barley and <0.003 for *Bh* were considered significant (Velásquez-Zapata *et al.*, 2024). Order of CSEP baits matches Figure 1 dendogram.

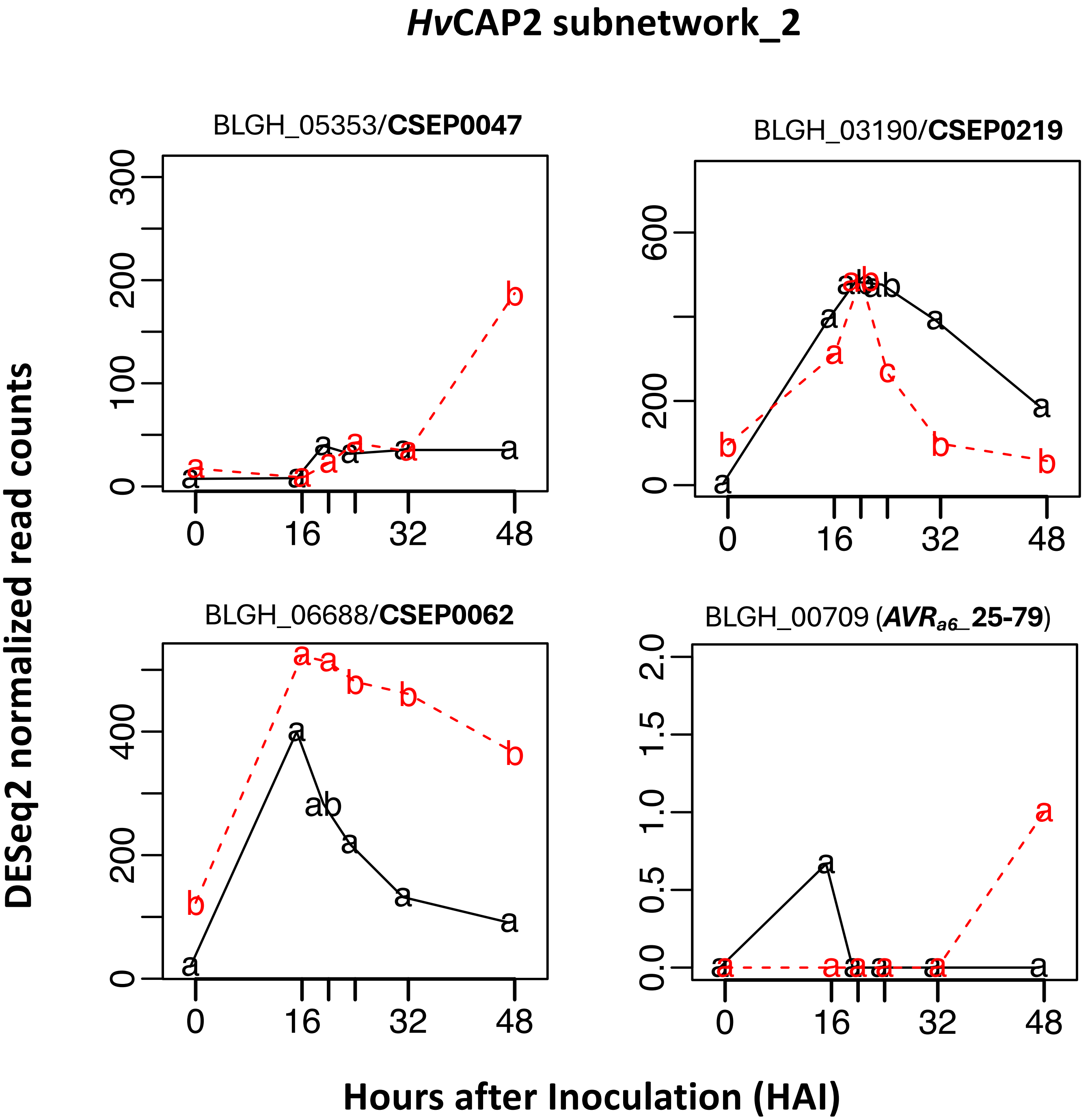


**Supporting Fig. S2-26.** Time-course expression of 48 *Blumeria hordei (Bh)* effector candidates, barley MLA, receptor-like kinase *Hv*SRF3_L1 (RLK5), and various host targets from the barley-*Bh* interactome. RNA-Sequencing data were extracted from an infection time course of barley CI 16151 (***Mla6*; black solid line**) and the derived fast-neutron susceptible mutant (***mla6*-m18982; red dashed line**), at 0, 16, 20, 24, 32 and 48 h after inoculation with *Bh* isolate 5874 ($AVR_{a1}$, $AVR_{a3}$, $AVR_{a6}$, $AVR_{a12}$; NCBI-GEO GSE101304) and analyzed as described in (Chapman *et al.*, 2021; Velásquez-Zapata *et al.*, 2022). Read counts were normalized using the taxon-specific method (Klingenberg and Meinicke, 2017). Genes differentially expressed at an adjusted *p* of <0.001 for barley and <0.003 for *Bh* were considered significant (Velásquez-Zapata *et al.*, 2024). Order of CSEP baits matches Figure 1 dendogram.

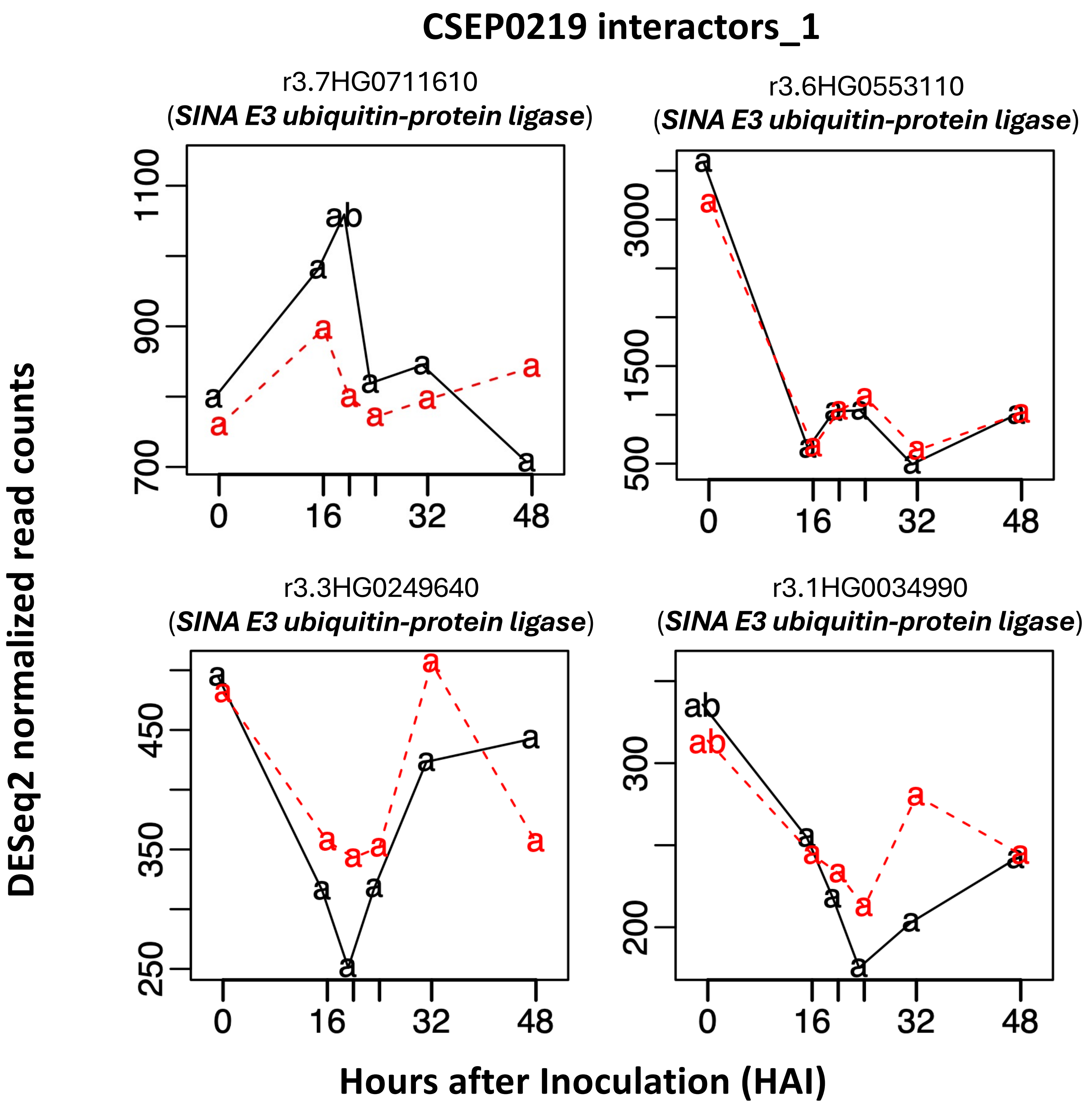


**Supporting Fig. S2-27.** Time-course expression of 48 *Blumeria hordei (Bh)* effector candidates, barley MLA, receptor-like kinase *Hv*SRF3_L1 (RLK5), and various host targets from the barley-*Bh* interactome. RNA-Sequencing data were extracted from an infection time course of barley CI 16151 (***Mla6*; black solid line**) and the derived fast-neutron susceptible mutant (***mla6*-m18982; red dashed line**), at 0, 16, 20, 24, 32 and 48 h after inoculation with *Bh* isolate 5874 ($AVR_{a1}$, $AVR_{a3}$, $AVR_{a6}$, $AVR_{a12}$; NCBI-GEO GSE101304) and analyzed as described in (Chapman *et al.*, 2021; Velásquez-Zapata *et al.*, 2022). Read counts were normalized using the taxon-specific method (Klingenberg and Meinicke, 2017). Genes differentially expressed at an adjusted *p* of <0.001 for barley and <0.003 for *Bh* were considered significant (Velásquez-Zapata *et al.*, 2024). Order of CSEP baits matches Figure 1 dendogram.

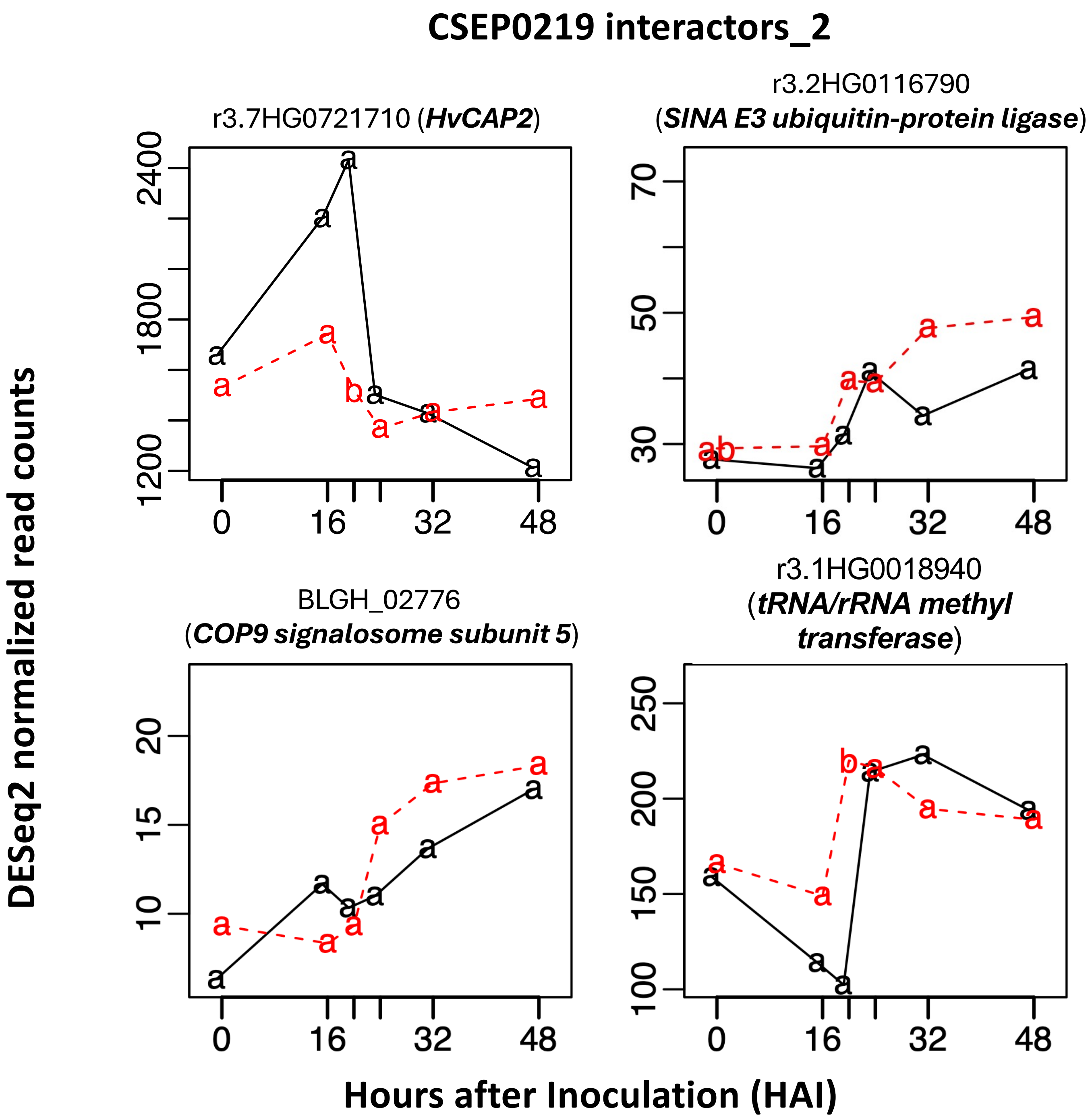


**Supporting Fig. S2-28.** Time-course expression of 48 *Blumeria hordei (Bh)* effector candidates, barley MLA, receptor-like kinase *Hv*SRF3_L1 (RLK5), and various host targets from the barley-*Bh* interactome. RNA-Sequencing data were extracted from an infection time course of barley CI 16151 (***Mla6*****; black solid line**) and the derived fast-neutron susceptible mutant (***mla6*****-m18982; red dashed line**), at 0, 16, 20, 24, 32 and 48 h after inoculation with *Bh* isolate 5874 ($AVR_{a1}$, $AVR_{a3}$, $AVR_{a6}$, $AVR_{a12}$; NCBI-GEO GSE101304) and analyzed as described in (Chapman *et al.*, 2021; Velásquez-Zapata *et al.*, 2022). Read counts were normalized using the taxon-specific method (Klingenberg and Meinicke, 2017). Genes differentially expressed at an adjusted *p* of <0.001 for barley and <0.003 for *Bh* were considered significant (Velásquez-Zapata *et al.*, 2024). Order of CSEP baits matches Figure 1 dendogram.
